# Blind prediction of complex water and ion ensembles around RNA in CASP16

**DOI:** 10.1101/2025.11.03.685595

**Authors:** Rachael C. Kretsch, Elisa Posani, Eugene F. Baulin, Janusz M. Bujnicki, Giovanni Bussi, Thomas E. Cheatham, Shi-Jie Chen, Arne Elofsson, Masoud Amiri Farsani, Olivia N. Fisher, M. Michael Gromiha, Ayush Gupta, Michiaki Hamada, K. Harini, Gang Hu, David Huang, Junichi Iwakiri, Anika Jain, Yuki Kagaya, Daisuke Kihara, Sebastian Kmiecik, Sowmya Ramaswamy Krishnan, Ikuo Kurisaki, Olivier Languin-Cattoën, Jun Li, Shanshan Li, Karim Malekzadeh, Tsukasa Nakamura, Wentao Ni, Chandran Nithin, Michael Z. Palo, Joon Hong Park, Smita P Pilla, Simón Poblete, Fabrizio Pucci, Pranav Punuru, Anouka Saha, Kengo Sato, Ambuj Srivastava, Genki Terashi, Emilia Tugolukova, Jacob Verburgt, Qiqige Wuyun, Gül H. Zerze, Kaiming Zhang, Sicheng Zhang, Wei Zheng, Yuanzhe Zhou, Wah Chiu, David A. Case, Rhiju Das

## Abstract

Biomolecules rely on water and ions for stable folding, but these interactions are often transient, dynamic, or disordered and thus hidden from experiments and evaluation challenges that represent biomolecules as single, ordered structures. Here, we compare blindly predicted ensembles of water and ion structure to the cryo-EM densities observed around the *Tetrahymena* ribozyme at 2.2-2.3 Å resolution, collected through target R1260 in the CASP16 competition. 26 groups participated in this solvation ‘cryo-ensemble’ prediction challenge, submitting over 350 million atoms in total, offering the first opportunity to compare blind predictions of dynamic solvent shell ensembles to cryo-EM density. Predicted atomic ensembles were converted to density through local alignment and these densities were compared to the cryo-EM densities using Pearson correlation, Spearman correlation, mutual information, and precision-recall curves. These predictions show that an ensemble representation is able to capture information of transient or dynamic water and ions better than traditional atomic models, but there remains a large accuracy gap to the performance ceiling set by experimental uncertainty. Overall, molecular dynamics approaches best matched the cryo-EM density, with blind predictions from bussilab_plain_md, SoutheRNA, bussilab_replex, coogs2, and coogs3 outperforming the baseline molecular dynamics prediction. This study indicates that simulations of water and ions can be quantitatively evaluated with cryo-EM maps. We propose that further community-wide blind challenges can drive and evaluate progress in modeling water, ions and other previously hidden components of biomolecular systems.

## 1 Introduction

It is a grand challenge in science to reach a predictive understanding of biomolecular structures accurate enough to enable the description and design of unseen biology and biomedicine. The tradition of structural biology has been to compare predictions to a single experimental model, yet the limitations of this approach are rapidly becoming apparent. The Critical Assessment of Structure Prediction (CASP) has been an important venue for enabling the field of structure prediction by organizing a biannual blind structure prediction challenge where groups across the world predict the 3D structures of proteins, and, since CASP15, nucleic acids^1–5^. Now the field aims to expand the goals of structure prediction towards a fuller understanding of biomolecular structure, including addressing four current limitations. First, biomolecules often function in concert with other molecules, such as proteins, nucleic acids, metabolites, or small-molecule drugs, so the interface between the molecules and binding-induced changes are important prediction challenges^3,6–8^. Second, an experimental structure is just a model of the underlying experimental data, which can introduce uncertainties^9^. Third, many biomolecules undergo conformational changes, which can be important for their function, for example, structural changes in a catalytic cycle^10,11^. Finally, many biomolecules contain disordered regions that are only well-described by an ensemble of structures^12,13^. In this study, we address the first, third, and fourth limitations by studying the ion and water structure around RNA by comparing directly to cryogenic electron microscopy (cryo-EM) density.

RNA is a fully solvated biomolecule because it has a negatively charged backbone and no positively charged moieties, precluding the possibility of forming a hydrophobic core buried from solvent, as in proteins. The importance of water and ions is particularly strong for nucleic acids since, without any positively charged moieties, electrostatic screening of backbone negative charge cannot be handled through intramolecular interactions such as salt bridges. Hence, interactions between RNA and water and ions, in particular cations like Mg^2+^, are important for the stable folding and function of RNA^14–16^. Yet, the majority of the electrostatic shielding does not come from highly ordered cations bound stably to the RNA backbone, but instead originates from a cloud of disordered ions surrounding the RNA^17,18^. Experimental measurements have been limited to highly ordered water and ions or biochemical measurements without 3D structural information. The study of solvation and ion interactions with RNA has therefore predominantly been limited to simulations, but the accuracy of these simulations is difficult to assess^15,19–21^.

Cryo-EM offers a new modality to compare simulations to experimental density. Unlike X-ray crystallography, the predominant source of experimental data on water and ion structures in RNA, cryo-EM does not isolate molecules in a crystal lattice but in vitreous ice, so the molecules are fully solvated. Additionally, cryo-EM enables the study of larger RNA molecules that may have more complex water and ion networks.

Prior to CASP16, two independent cryo-EM maps of the *Tetrahymena* ribozyme, an RNA enzyme, were obtained at high resolution, 2.2 and 2.3 Å^22^, where the core of the ribozyme was very similar. In these maps, ordered peaks were identified where water and ions could be modeled. However, the cryo-EM maps additionally contained diffuse density that was reproducible between the two independent reconstructions of the ribozyme core. The density in the solvent shell could not be fully modeled by ordered waters and ions, suggesting that an ensemble of solvent structures was better suited to describe the cryo-EM map. The CASP16 solvent ensemble prediction challenge took advantage of these cryo-EM maps – unpublished at the time of the challenge – to pilot the concept of comparing solvent ensemble prediction to the cryo-EM maps and to rigorously establish the state of the art in modeling water and ions.

## 2 Methods

### 2.1 Challenge set-up

A broad call for participation in this pilot ensemble challenge was sent to the computational structure prediction community, including members of the molecular dynamics (MD) community who have not typically participated in CASP single-structure prediction categories. This resulted in submissions from 26 prediction groups for the solvent ensemble challenge, labeled R1260 in CASP16 target labeling. Predictors were provided with the RNA sequence, in addition to information about the experimental conditions, what could be submitted, and how they would be assessed (**Box 1**). Importantly, the predictors were aware that the task was to predict a “cryo-ensemble” of water and ions by submitting up to 1,000 models and that the assessment would be against cryo-EM maps directly. The availability of high-quality reference coordinates for the RNA atoms in the *Tetrahymena* ribozyme, from the PDB ID 7EZ0, allowed predictors to focus on the water and ions that were not captured in previous maps. The submission portal was open for two months.

##### Information provided to predictors regarding R1260

Predictors are invited to submit an ensemble of conformations of this RNA with the solvent shell. The structure of the local solvent shell (water and ions) will be compared against cryo-EM data. We provide the following experimental information. These samples were folded and frozen in a solution of 50 mM Na-HEPES pH 8.0 and 10 mM MgCl2. The sample was folded for 30 min at 50°C, incubated at room temperature for 10 min, and then placed on ice. The sample was frozen on Vitrobot Mark IV with 4 s blot time, 4°C, and 100% humidity using liquid ethane as cryogen. A total electron dose of 57.25 electrons per Å2 was used during data collection. The cryo-EM maps are 2.2 and 2.3 Å according to gold standard FSC and were auto-sharpened using PHENIX. We require, for each conformation, that the RNA coordinates be present and that you submit at least a 5 Å shell of water and ions. Larger solvent shells are allowed but may not be assessed. The RNA conformation may change across the frames. The RNA motion will not be assessed but the predictors are welcome to submit RNA of different conformations. Predictors can assume that the RNA conformations are similar to 7EZ0 (e.g. < 4 Å RMSD). During assessment these models will be locally aligned before comparing solvent structure. The main assessment will be on the local water and ion structure surrounding each RNA region, based on local alignments compared to experimentally determined high-resolution maps. Given experimental uncertainty in some regions, the solvent structure around the following residues will not be assessed: 42-67, 208-224, 263-274, 311-314, 343-381 (7EZ0 numbering: 63-88, 229-245, 284-295, 332-335, 364-402). To reduce file size, please remove all hydrogen atoms. Please submit as a set of PDB files, these should be tarred, gzipped, and uploaded as one archive file through https://predictioncenter.org/casp16/predictions_submission_WATER.cgi. To create an archive, please run: tar -czf R1260TS234.tgz Folder_with_R1260_models. Archive file names should combine the target name R1260 and the group name (234 in the provided example). Names of individual models inside the archive should follow the CASP model naming scheme, e.g. R1260TS234_1, where _1 is the model number. You may submit up to 1,000 conformations, with a suggested minimum of 10. We will assume equal weighting of all models in the ensemble; you may repeat conformations in the ensemble if you wish to give those conformations greater weight.

### 2.2 Prediction of water and ion structure around RNA

All prediction groups were offered the opportunity to submit a written methods section for inclusion in this manuscript. The methods of the predicting groups are summarized in **Table 1** and sections describing the prediction of 18 prediction groups (from 12 laboratories) are found in **Supplemental Text 1-12**.

**Table 1:**
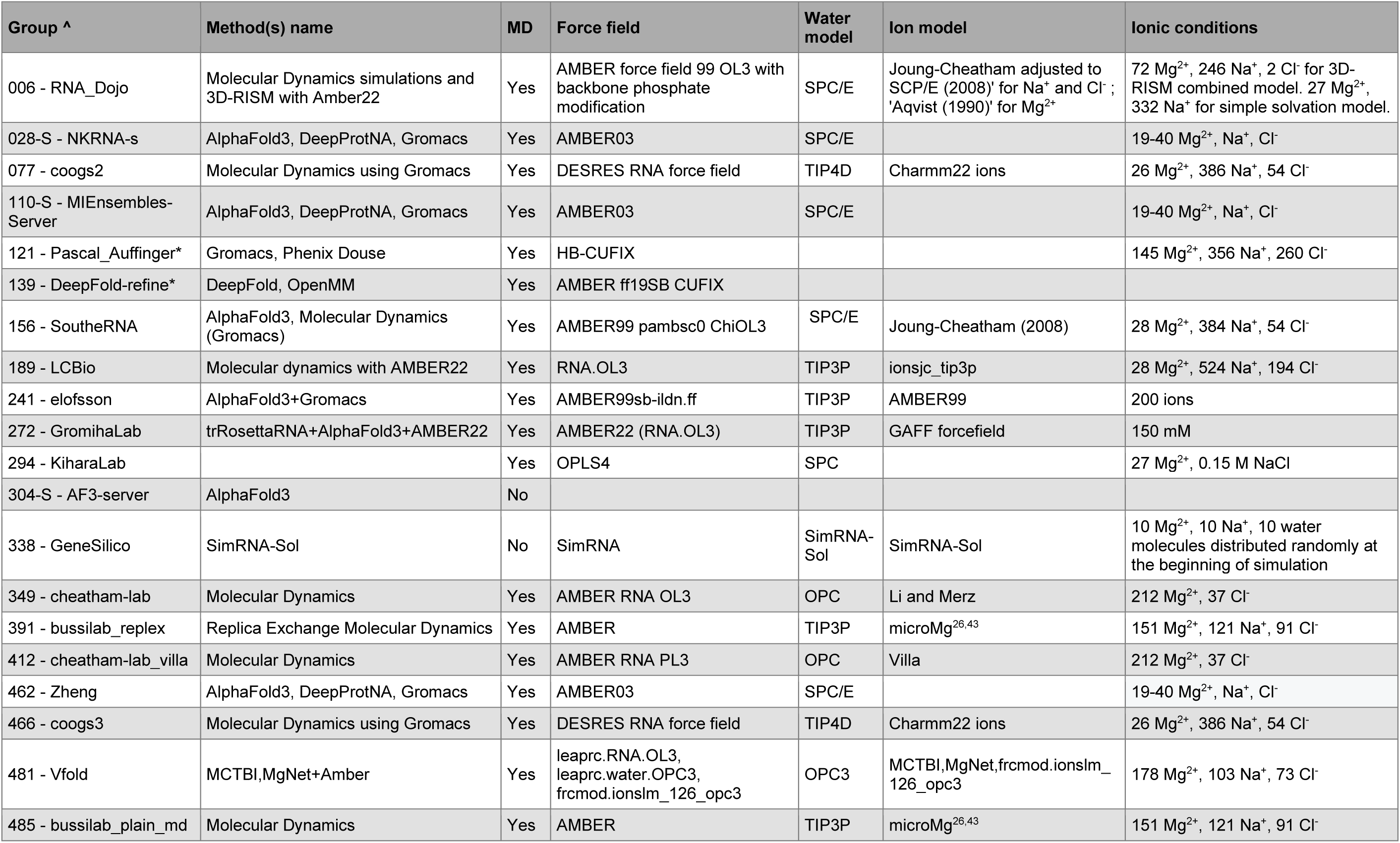
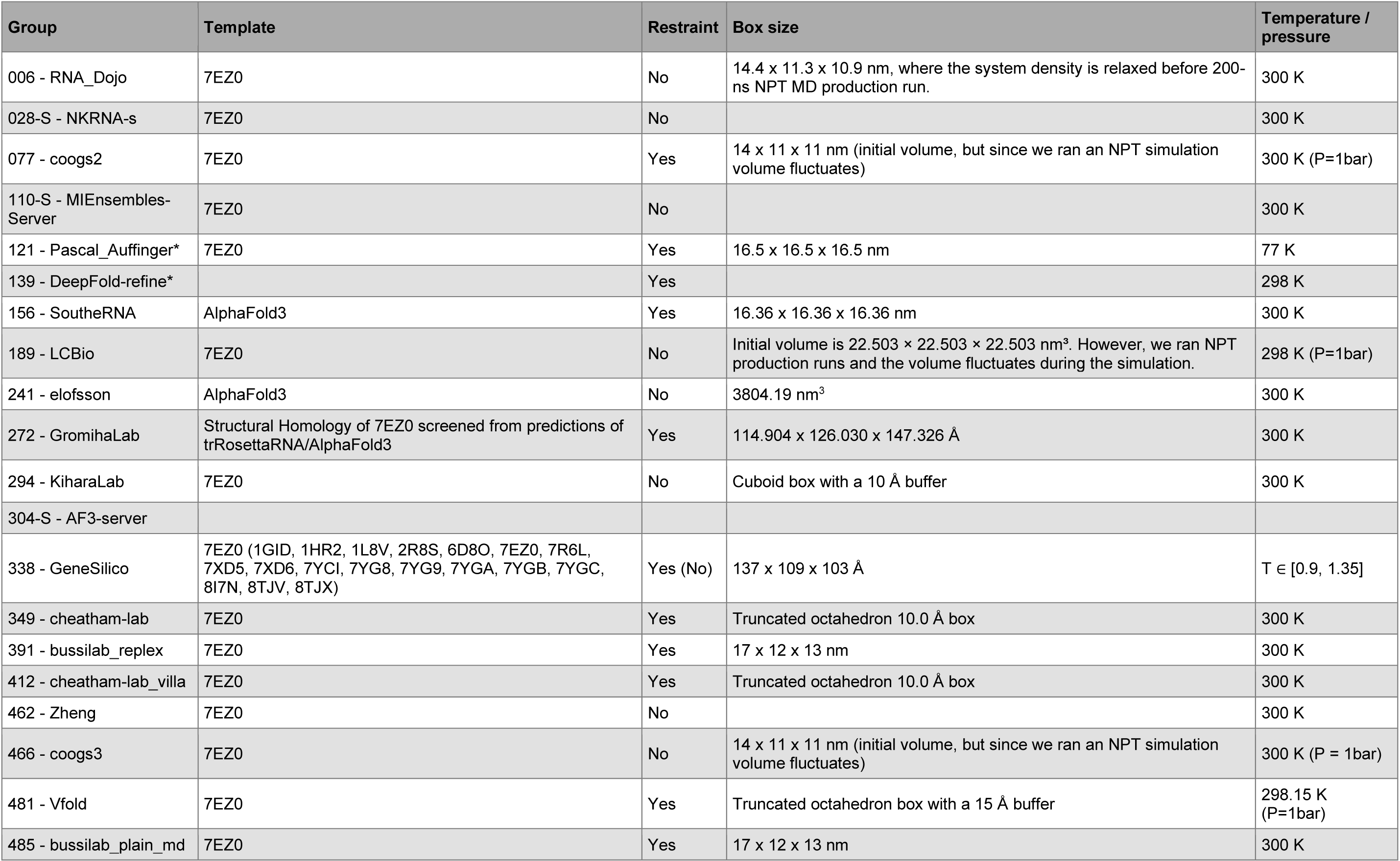
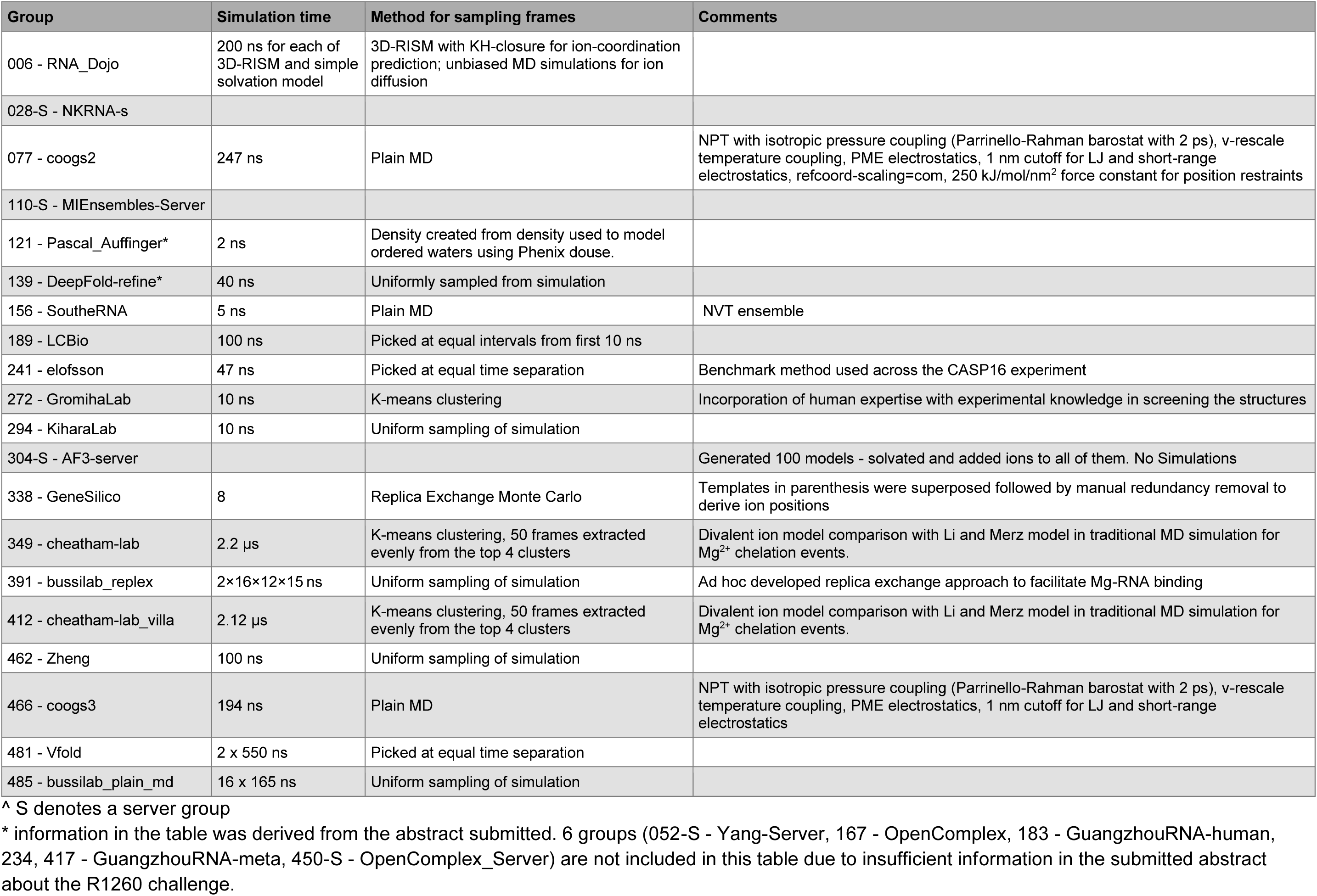
Group methods.

CASP16 predictions were compared against a variety of baselines and reference points. The performance ceiling was calculated by the comparison of the reference cryo-EM map (EMD-42499, solved at 2.2 Å resolution and deposited after the challenge) and an independent cryo-EM map (EMD-42498, solved at 2.3 Å and also deposited after the challenge)^22^. The deposited EMDB maps had similar sharpening B-factors (79.23 and 74.07 Å^2^, respectively)^22^. When sharpening of the independent map was varied, the maximal agreement to the reference map in the solvent shell occurred near the deposited B-factor. Hence, the deposited maps were used but sharpening remains an important factor to consider in future studies. The performance floor was calculated by randomly shuffling the 2.2 Å map solvent shell density values.

To compare to the static single model description of the density, we also included the PDB models derived from the cryo-EM maps (and released after the challenge); 9CBU and 9CBW contain the water and Mg^2+^ ions modeled by an automated cryo-EM map-guided algorithm, SWIM^23^, that were in consensus between the two maps, and 9CBX and 9CBY contain all the water and Mg^2+^ ions identified by SWIM in the 2.2 and 2.3 Å maps respectively. These are labeled as “Cryo-EM model con.” for 9CBU; “Ind. cryo-EM model con.” for 9CBW; “Cryo-EM model” for 9CBX; and “Ind. cryo-EM model” for 9CBY.

Finally, as further baselines, we extracted 1,000 uniformly sampled frames from the six simulation methods used in the manuscript describing these cryo-EM maps^22^. Briefly, these simulations were equilibrated from a starting pose derived from the 2.2 Å map with 20 Mg^2+^ ions placed for stability purposes; because of these experimental biases they do not serve as completely blinded baselines but offer a reference of performance. A total of 95 Mg^2+^ ions and 196 Na^+^ ions were used to neutralize the system; these were placed using two different methods, random placement and guided by Coulombic potential using “addIons” in LEaP^24^. Three force fields were used; “Baseline” is the DESRES modified RNA force field^25^, while “Baseline mMg” and “Baseline nMg” used parmBSC0χOL3 with the mMg and nMg parametrization of Mg^2+^, respectively^26–30^. This resulted in six simulation types where explicit TIP4P-D water^31^ was used for all.

These atomic ensembles were treated identically to the predicted ensembles from CASP16 participants in the next steps of creating a density representation and comparing that density to the cryo-EM map.

### 2.3 Conversion of atomic ensemble to densities

To compare the ensemble of atomic structure to the cryo-EM map directly, the atomic ensembles were converted to densities (**Figure 1**). First, the models were cleaned, ensuring the RNA, water, and ions were standardized: RNA, water, Mg^2+^, Na^+^, Cl^-^, and K^+^ molecules were all accounted for, and any other molecule was assigned as a water. Following the challenge guidelines, many groups removed water and ions more than 5 Å from the RNA. For consistency in evaluation, atoms beyond 5 Å from the RNA were removed before assessment for the other groups as well. It remains for future studies to determine the contributions of more distant water and ions on the solvent shell density.

**Figure 1:**
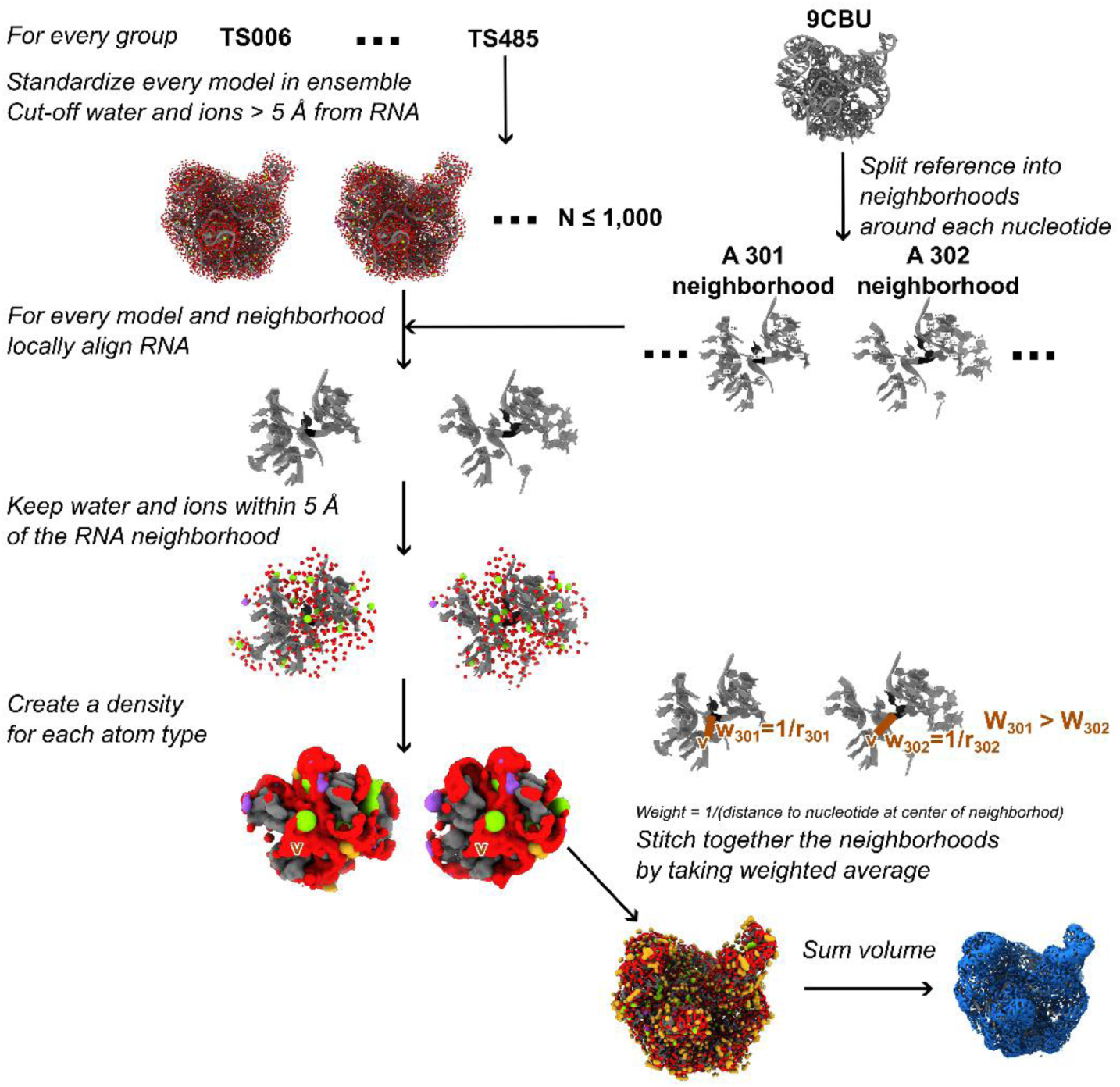
Conversion of atomic ensembles to densities.

If all models in the ensemble were aligned globally, deviations in RNA structure from, e.g., different frames of a molecular dynamics simulation, would dominate and blur out any densities from the water and ions. To ensure appropriate averaging of the local water and ion structure, regions around each residue were aligned individually, the local water and ion structure was determined, and then all the local water and ion structure was stitched back onto the reference RNA model. The neighborhood around each residue was defined as any RNA residue that had at least one RNA atom within 10 Å of the residue in the reference RNA structure. The analysis was also run with a 6, 12, and 20 Å radius for comparison of the effect of aligning neighborhood size. The alignment followed the Kabsch algorithm using all heavy atoms in the RNA. The analysis was also run using backbone atoms, a 5-atom selection (P, C4’, C2, and N9 and C6 (for G and A) or N1 and C4 (for C and U)), and a 3-atom selection (P, C4’, and N9 (for G and A) or N1 (for C and U)) for comparison. The water and ions within 5 Å of this local neighborhood were extracted and aligned according to the RNA alignment transformation.

The water and ions around every RNA were then converted to a density representation. This was done in two ways. First, Density_scat_ was applied, using atomic scattering amplitudes based on the approach of Hoff et al^32^, which uses a 5-Gaussian fit of the electron scattering factors from Peng et al.^33^. Second, Density_prob_ was applied, using a probability density calculated using the DensityAnalysis tool in MDAnalysis^34,35^. Atomic B-factors were ignored in this analysis because not all groups submitted B-factors, and it was assumed that thermal fluctuations would be represented within a 1,000-model ensemble. These local densities were then stitched together through weighted averaging, where for every voxel (0.82 x 0.82 x 0.82 Å), all local regions which contained that voxel were averaged using a weight of 1/r where r was the minimum distance from the voxel to the RNA residue at the center of that region. This weighting was used to create a smooth density while ensuring the local water and ion densities with the most accurate alignment for that position contributed most. Trilinear interpolation was used to calculate density values in between discrete voxel points when needed.

For both methods, the densities are additive (i.e., converting a model with water and ions to density is the same as converting the water to density, converting the ions to density, and adding the two densities), so the densities for RNA, water, and each ion type were converted separately to enable visualization and further analysis. All figures depicting the densities were created in ChimeraX^36^.

Additionally, five bootstrapped volumes were created for each predicted ensemble. Models of the ensemble were randomly selected with replacement to obtain a bootstrapped sample, and an RNA, water, and ion density was created for this sample following the same procedures as above. The standard deviation of scores from these bootstrapped samples was used as an indication of variance for each predicted ensemble score. These variances calculated within group predictions were smaller than the typical variances of scores between groups.

### 2.4 Comparison of densities to cryo-EM data

The predicted atomic ensembles were now represented as densities and could be directly compared to the cryo-EM density. First, the solvent shell was defined and only voxels within that shell were used for comparison, so density closer or farther to the RNA would not be assessed. As stated to predictors in the challenge announcement (**Box 1**), residues that were poorly resolved (63-88, 229-245, 284-295, 332-335, 364-402) were not considered; any voxel that was nearest to one of these residues was not used. To capture both directly coordinated water and ions we primarily analyzed the shell that was 1.8-3.2 Å away from the RNA. Beyond 3.2 Å there are important ions coordinating to RNA in their second shell, through a water, however, at 2.2 Å the uncertainty and noise in the cryo-EM data is not negligible: the Spearman correlation between the two independent maps falls with increasing distance from the RNA (**Supplemental Figure 1**). Hence, the second solvation shell and beyond were not the focus of this study and remain for future investigations with data with higher signal-to-noise ratios in those distal regions. For investigating the variation in accuracy based on location, regions were also isolated as follows: (1) solvent shells of increasing size from 1.8-1.9 Å up to 1.8-5.0 Å increasing by 0.1 Å, (2) 1 Å windows of increasing distance from the RNA from 1.8-2.8 Å to 4.0-5.0 Å, and (3) local regions around each residue; the 1.8-3.2 Å solvent shell was broken into residue regions where voxels were grouped by their closest RNA residue.

The isolated voxels were compared to the same voxels in the reference cryo-EM map (EMD-42499) using four metrics. All real-space measurements were chosen because any Fourier space measurement would be affected by the masking of the solvent region. First, the traditional measure of correlation coefficient used in cryo-EM^37^, sometimes called cross-correlation coefficient, is equivalent to the Pearson correlation coefficient *CC* (in the equation below x is the predicted density value, *x*_o_ is the reference density value, V is the set of voxels being compared):

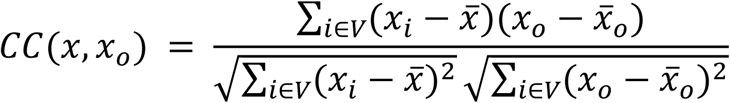

The Pearson correlation coefficient, although the most popular correlation measure used in cryo-EM, is most appropriate for Gaussian distributions, which are not perfect descriptions, of all predictions in the masked region. So, the Spearman correlation coefficient was also measured which calculates the correlation of the rank of density values, *R*(*x*):

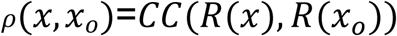

Third, the mutual information^38^ was calculated using sklearn mutual_info_regression with a neighborhood of 6^39–41^. For some individual residue masks, there was an insufficient number of voxels to calculate mutual information and those regions were not included in comparisons. Alternative measures that could be used to compare the distribution of density values, including Manders overlap coefficient (MOC), remain for future studies.

Finally, the assessment was restructured as a classification task asking how well the predictors were able to predict the location of high occupancy (density > 3 σ) in the cryo-EM map. The precision, recall, and false positive rate were then calculated:

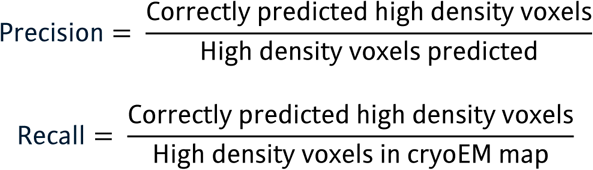

These values were calculated at various density thresholds for the predicted ensemble and a precision-recall curve was plotted. The area under these curves (AUC-PR) quantifies how predictors perform on identifying the high density voxels. There is an imbalance between the high and low density classes because cryo-EM density is predominantly empty, density peaks are sparse, so the area under the receiver-operating characteristic curve (AU-ROC) and a simple accuracy measurement are ill-suited metrics and were not used here ^42^. Alternative measures that could be calculated based on this restructuring as a classification task include Matthews Correlation Coefficient (MCC) and F_β_ scores. Due to the relatively low prediction accuracy, the comparison of the alternative metrics could not be investigated in this pilot study and remains for future studies.

As a measure of experimental uncertainty, the independent cryo-EM map (EMD-42498) was aligned to the RNA in PDB-9CBU and resampled onto the reference cryo-EM density voxels using trilinear interpolation. The same voxels were isolated and compared to the reference map using the same metrics, and the independent cryo-EM map (EMD-42498) was used as the reference instead of the original reference map (EMD-42499). The deviation of scores for each predicted ensemble when compared against the two independent maps (EMD-42498 and EMD-42499) provided an estimate of uncertainty due to statistical error in the cryo-EM maps. Finally, as a “floor” reference, the masked voxel values of the reference cryo-EM map were randomly shuffled and compared to the ordered vector.

To identify performance outliers, we took the Z-score (i.e., the number of standard deviations by which the model’s accuracy differed from the mean) of scores across groups.

### 2.5 Comparison to cryo-EM derived model

No systematic comparison of predictions to the cryo-EM derived atomic models was made due to the aforementioned limitations in the cryo-EM derived atomic model such as only modeling highly ordered water and ions and the uncertainty of peak identity (e.g., ion type) at 2.2 Å map resolution, where waters coordinating ions are not well resolved. Some prediction groups did conduct comparisons of their predictions and the atomic models, which can be found in **Supplemental Texts 13-20**.

## 3 Results

### 3.1 Group participation and methods

26 predictions were submitted to the R1260 challenge. The majority of groups used MD simulations to sample water and ion structure but differed in their force field, water and ion models, simulation conditions, and frame sampling (**Table 1**). Twelve laboratories described the methodologies used in 18 groups submitted in the Methods section, and the description of two additional group methodologies can be found in the CASP16 abstract book.

The majority of groups submitted ensembles of more than 100 coordinate models with low or moderate root mean squared deviation (RMSD) to the experimental RNA conformation (< 10 Å RMSD; **Figure 2A-B**). More than 12,000 models were submitted, comprising more than 4.5 million RNA residues and more than 100 million RNA atoms. All but one group submitted water and ions, for a total of more than 250 million water molecules and more than 2 million ions solvating the RNA (**Figure 2C-D**). The ionic composition of the submission varied between groups; cations Mg^2+^ and Na^+^ were most prevalent and were described to predictors as included in the experiment (**Box 1**). Cl^-^ and K^+^ were also included in some submissions (**Figure 2D**). Within most ensembles, particular water and ion molecules had characteristic coordination distances to the RNA, with Mg^2+^ ions bound the closest, followed by Na^+^ ions, and then water (**Supplemental Figure 2**). Additionally, some ensembles had a second Mg^2+^ peak, corresponding to Mg^2+^ coordinating RNA indirectly through an intermediate water molecule (**Supplemental Figure 2**).

**Figure 2:**
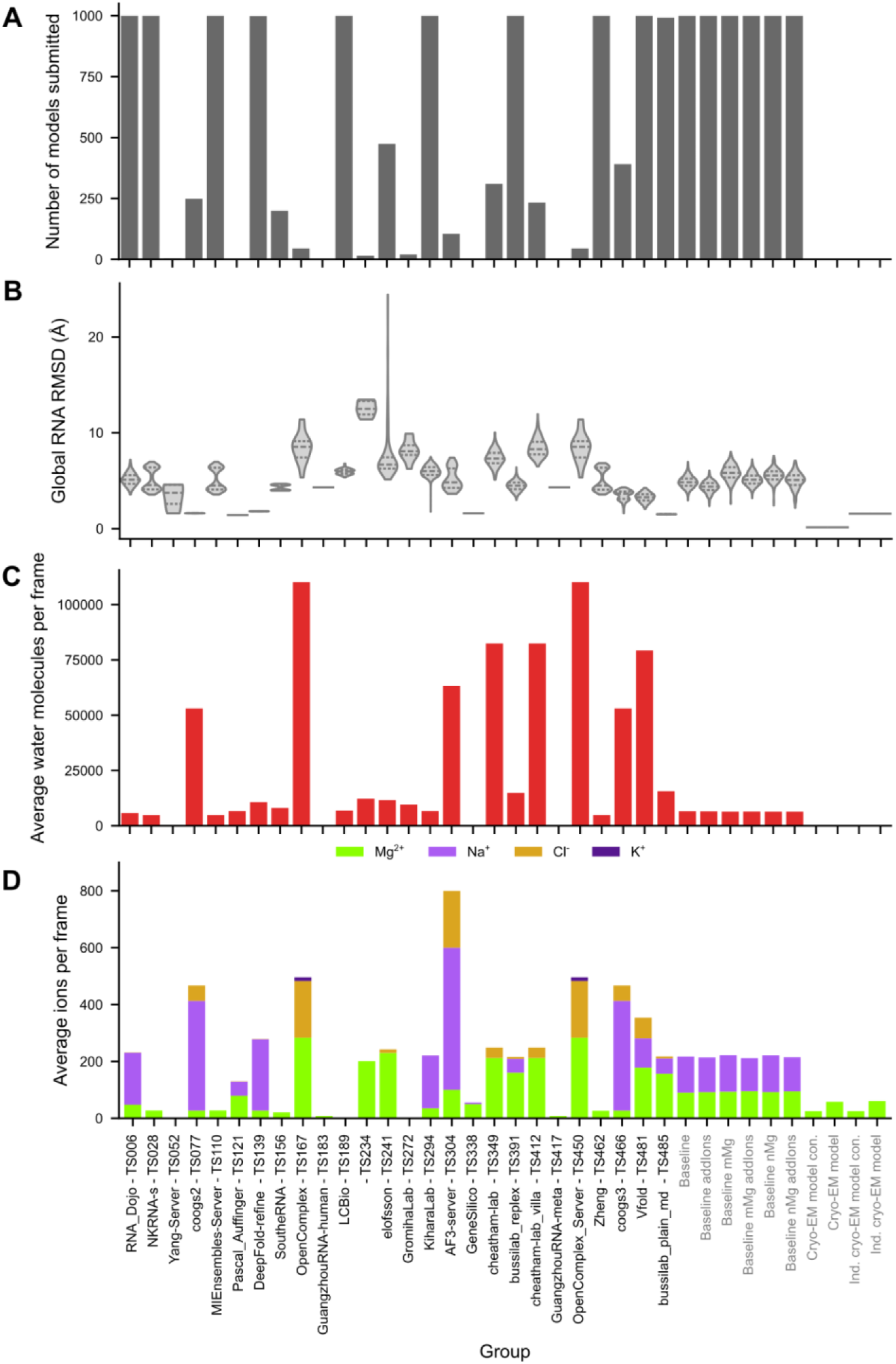
Predicting group participation. (**A**) Number of models submitted per group (maximum 1,000). (**B**) For every group, the distribution of global RMSD of the RNA heavy atoms of every submitted model in the ensemble to 9CBU is shown in a violin plot, the median and interquartile ranges are additionally marked with dashed lines. For every group, (**C**) the average number of water molecules submitted per model and (**D**) the average number of ions submitted per model are displayed. In (**D**) Mg^2+^ ions are colored green, Na^+^ are light purple, Cl^-^ are dark yellow, and K^+^ are dark purple. Some groups submitted water and ions beyond 5 Å from the RNA, explaining the very high counts for those groups in (**C**) and (**D**).

### 3.2 Generation of densities from atomic ensembles

All predicted, baseline, and reference ensembles were converted to density maps as described in **Methods** (**Figure 1**). Importantly, to obtain the water and ion structure around each residue, independent of RNA global motion, this procedure locally aligned water and ions around every residue. Even in ensembles that had large global motion (**Figure 2B**) the local RNA structure was very similar (**Supplemental Figure 3**). Results were similar when all heavy atoms were aligned or subsets of these atoms were aligned. The local alignment was affected by the size of the local region only when the local radius used for the alignment procedure was very large, 20 Å or greater (**Supplemental Figure 3**).

Based on the local water and ion structures, densities were generated and stitched together to obtain full water and ion shells around the RNA. Separate maps were generated for each molecule type enabling the visualization of structure around the RNA. Local regions of the ribozyme can be visualized to show similarities and differences between the predicted density and the cryo-EM density (**Supplemental Figure 4** and **Supplemental Figure 5**). These densities were generated in the two ways as described in **Methods**: using atomic scattering factors, where signal for each atom extended to 4 Å away from the atom, and using probability density representation, where the presence of a molecule at a location was simply counted at that location. These methods resulted in visually similar densities, particularly for predicted ensembles with many models, because the deviations in water and ion positions within the ensemble smoothed out the probability density (**Supplemental Figure 6**).

### 3.3 Correlation of solvent shell

The density representation of the predicted ensembles was directly compared to the cryo-EM map. To measure how similar the predicted solvent shell was to the cryo-EM map, the voxels surrounding the RNA (1.8-3.2 Å) were isolated, removing voxels near poorly resolved regions. These density values were then compared to the same region in the cryo-EM map (see **Methods** for description of comparison metrics).

Correlation coefficient (CC) is a standard metric used in cryo-EM to evaluate real-space similarity of two densities^37^; it is equivalent to the Pearson correlation coefficient between the sets of voxel values. To assess the variability in CC score on the models selected, each ensemble was sampled with replacement five times. The variation in CC scores for the bootstrapped models was lower than the variability between groups, indicating low variability due to sampling. To assess the variability in CC score on the cryo-EM map used as reference, we calculated the CC score when comparing it to the alternative independent cryo-EM map. The CC score was similar using either cryo-EM map (**Figure 3A**). These two conclusions held across scoring in this study. These results indicated that the differences in scores between groups were not dependent on the model selection or cryo-EM map selection (**Figure 3**).

**Figure 3:**
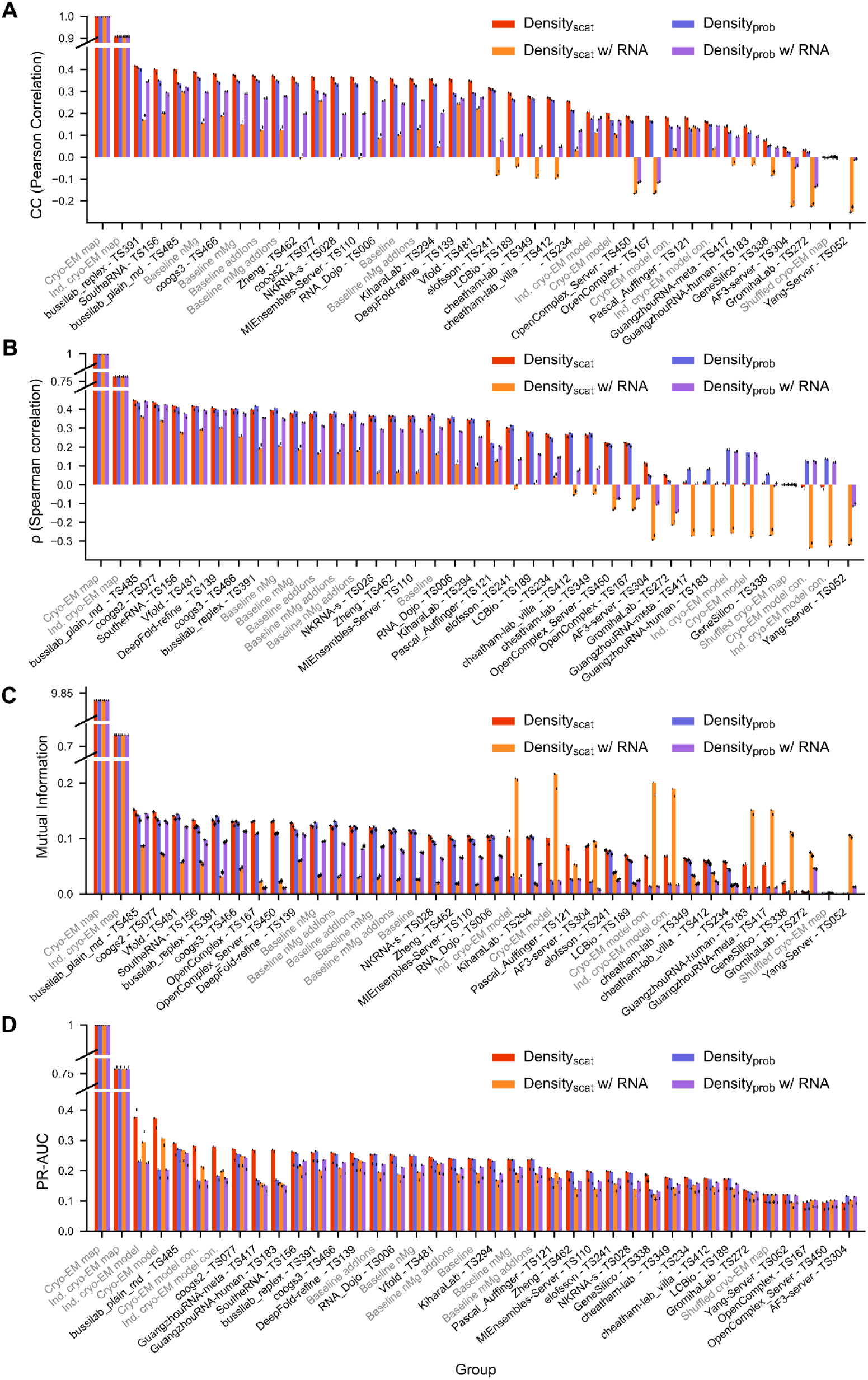
Comparison of predicted ensembles and the solvent shell of the cryo-EM map. For each ensemble, a density was calculated based on scattering amplitudes (density_scat_) or probability density (density_prob_) of the water and ions (red or blue, respectively) or of the water, ions, and RNA (orange or purple respectively). The densities were compared to the 2.2 Å cryo-EM map (EMDB-42498) in the shell 1.8-3.2 Å away from the well-resolved RNA residues. The density values were compared using multiple metrics: (**A**) standard cryo-EM cross correlation, equivalent to the Pearson correlation coefficient, (**B**) Spearman correlation coefficient, (**C**) mutual information, and (**D**) area under the precision-recall curve where voxels were classified as positive if their cryo-EM density was > 3 σ. For each bar, the score when comparing against the independently resolved cryo-EM map (EMDB-42499) is represented by a black dot. In each plot groups are ordered by performance with the density_scat_.

The Bussilab_replex, SoutheRNA, and bussilab_plain_md ensembles exhibited the highest Pearson correlation to the cryo-EM density when a scattering factor-based density procedure was used; their performance was higher than the baseline MD ensembles generated by challenge organizers with knowledge of the cryo-EM map (**Figure 3A**). Nevertheless, for all groups, correlation was significantly lower than the maximum achievable given cryo-EM experimental uncertainty (CC(bussilab_replex) = 0.42 vs. CC(independent cryo-EM map) = 0.91). All groups, except Yang-server, which did not submit water or ions, obtained higher correlation than the random shuffle (CC=0.000092) (**Figure 3A**). Most simulations also exhibited higher correlation with the cryo-EM density than the static SWIM reference atomic model (CC = 0.20), despite the latter being fit directly to the cryo-EM density map. This result suggests that predicted ensembles captured information in the cryo-EM map that was left unmodeled by the SWIM reference atomic model, which only contains ordered waters and Mg^2+^ ions (**Figure 3A**). This suggests that the current practice in deriving a single model from a cryo-EM map does not necessarily fully represent the dynamic nature of the surrounding water molecules and ions, which are indeed observed in the experiment but not reported in the PDB.

To reduce assumptions about the distribution of density values further, we measured the mutual information^38^ and Spearman correlation coefficient. Mutual information and Spearman correlation ranked groups in similar order (**Figure 3B-C**). Together with the Pearson correlation coefficient results, this analysis highlighted seven groups of interest (listed in numerical order by group number): coogs2, DeepFold-refine, SoutheRNA, bussilab_replex, coogs3, Vfold, and bussilab_plain_md (**Figure 4**).

**Figure 4:**
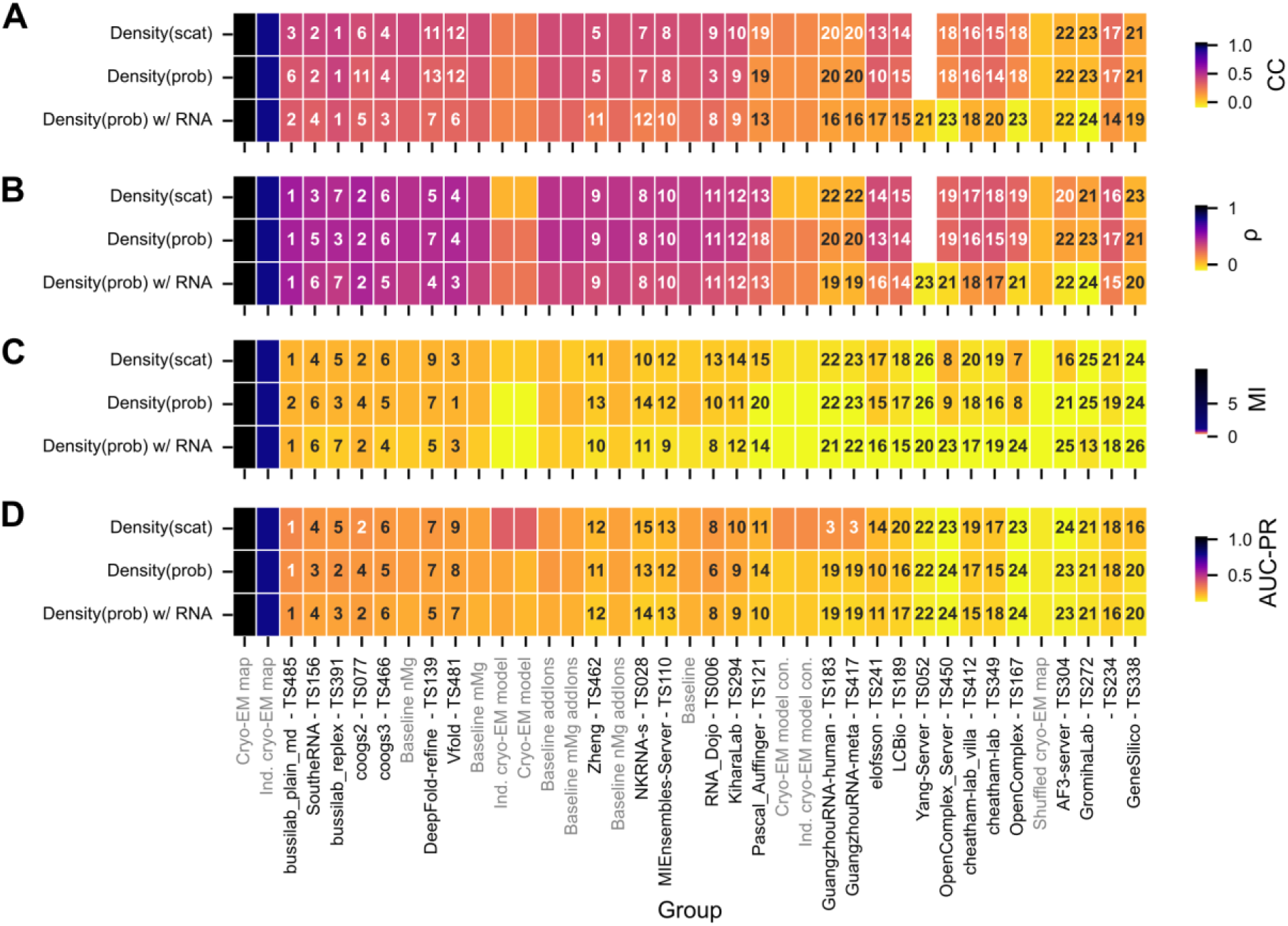
Comparison of ranking based on various metrics. For each metric the group is colored by score from experimental error (blue) to the lowest score (yellow). The ranking is labeled in text. The groups are ordered by the Z-score based ranking for Density_scat_ where the groups’ Z-score for each of the four metrics is summed. Z-scores are clipped at zero so if the group performed below average for a metric (Z<0) they are not penalized. References and baselines are colored in gray.

### 3.4 Impact of density generation methods

When a probability density-based procedure was used, all predictions exhibited a lower correlation to the cryo-EM map, and the bussilab_replex prediction scored noticeably higher than other groups (**Figure 3A**). This difference between scattering factor based-density generation and probability-based density generation was diminished when looking at the Spearman correlation coefficient, suggesting the difference was due to the scaling of density values and not to differences in accuracy of predicting the most dense positions (**Figure 3A-B**). The reference cryo-EM model, where water and ions were modeled into reference cryo-EM density peaks, received a higher correlation score when the density was generated with scattering factors. Accuracy metrics calculated from scattering factor-based densities are focused on in the following sections as more representative of the cryo-EM signal.

The predicted densities included water and ions only, but signals from the RNA extended into the solvent shell in both the experimental and predicted densities. When RNA was included in the predicted density, there was a decrease in correlation, particularly for the scattering factor-based densities (**Figure 3A**). This resulted in a negative correlation in some instances; all these groups predicted no or a limited amount of ions so this negative correlation upon including RNA was explained by underprediction. The reduction of correlation upon including RNA could be caused by two factors. First, the method of converting RNA into density may differ from the actual cryo-EM signal. Second, in each local alignment, the more distal RNA residues were not always well-aligned and were therefore placed into the solvent shell resulting in an aberrant signal. This effect should be reduced by the weighting when stitching together the densities, which would down-weight these distal RNA residues in the final volume.

Examining the probability density measurement helped to differentiate between these two factors, because the atomic signal is not spread beyond the voxel the atom is placed in (factor 1 is less apparent). With the probability density representation, we did not see major decreases in correlation of the top groups when RNA was included compared to when it was not. The groups that showed a change were generally groups that had RNA models that differed from the reference more, which would explain why they had more aberrant RNA signals (**Supplemental Figure 3**). There was a more significant drop in correlation for scattering factor-based densities. In this approach, the Gaussians from each atom were calculated out to 4 Å, and thus the RNA signal was overlapping with the solvent shell. The decrease in correlation suggested the density generation method did not match the cryo-EM map well, so the densities generated without RNA are focused on in following sections.

Given the potential influence of alignment accuracy, we returned to the various alignment methods tested above. The method of locally aligning each RNA residue generally did not change the resulting correlation with the cryo-EM density, except for when the alignment radius was set so high such that the alignment was no longer local in nature (**Supplemental Figure 7**). As previously suggested by the poor alignment accuracy, using a 20 Å radius for alignment reduced the overall agreement with the cryo-EM density. This suggested that it was important to conduct local refinement with a small radius (≤ 12 Å) to correctly align the water and ion network ensembles.

### 3.5 Impact of the definition of solvent shell

We also compared accuracy estimates across various definitions of solvent shells. The definition of the solvent shell of 1.8-3.2 Å was selected to contain both water molecules and ions directly bound to the RNA. When this shell radius was increased, we observed an increase in experimental error: Pearson correlation between the two cryo-EM maps only decreased slightly, still maintaining a CC > 0.8 until 5.0 Å; however, there was a large decrease in Spearman correlation between the two independent cryo-EM maps (**Supplemental Figure 8**). Finally, the AUC-PR between the experimental maps peaked at 3.2 Å, suggesting that this was a reasonable mask radius for the current analysis, but should be re-evaluated for future cryo-EM map challenges. Interestingly, many of the scores for the predicted ensembles also peaked around 3.2 Å.

To observe whether any group performed significantly better within a given distance of the RNA, we calculated the scores for 1 Å windows of the solvent shell. The performance of the reference atomic model, which had water and Mg^2+^ ions placed in the density, peaked at roughly 2.5-3.5 Å likely because that distance contains the most amount of modeled waters and ions; no waters and ions were modeled past 3.5 Å (**Supplemental Figure 1**). Otherwise, the predicted ensemble score dropped in a similar trend to the experimental error, suggesting there was not a particular shell of higher accuracy (**Supplemental Figure 1**).

### 3.6 Classification of high density regions

As can be seen in the ranking above, correlation-based metrics biased scores towards groups that fully predicted the solvent shell by submitting sufficiently large ensembles with a fully solvated shell. However, there were some groups that only submitted a few models, with a limited number of water and ions, some as few as in the cryo-EM reference model which contained highly ordered water and ions only.

The area under the precision-recall curve (AUC-PR) is a unique metric as it focuses on the differentiation of the low and high density values. This classification is an important goal for prediction, as the highest occupied sites (most dense) are expected to be the most energetically and functionally important. Unlike the other metrics where the SWIM cryo-EM reference model performed poorly because it left most density unmodeled, it performed best here because this metric focused on the high density regions where water and ions were modeled in this atomic model reference (**Figure 3D**). Hence, a group with a small, underpredicted solvent shell could score well in AUC-PR.

With AUC-PR, we observed improved relative performance by a group not performing well on correlation metrics, GuangzhouRNA (GuangzhouRNA-human - TS183 and GuangzhouRNA-meta - TS417), who submitted a very small ensemble of 1 model with only 189 waters and 8 Mg^2+^ ions (**Figure 2**, **Figure 3A**). Examining the precision-recall curve itself revealed more about these performance differences (**Figure 5A**). When tasked with predicting which voxels had a high (> 3 σ) density in the cryo-EM reference map, at the contour level of the predicted map leading to two-third precision, GuangzhouRNA was able to recover 9% of the high-density positions. This was in contrast to the 2% of positions that the next best ensemble, bussilab_plain_md, was able to recover. The cryo-EM-derived SWIM atomic models recovered 22% of high-density voxels when considering all the water and ions (9CBX) but only 13% when only the most confident water and ions were considered (9CBU). All of these performed worse than experimental reproducibility: 70% of high-density voxels were recovered by an independent solved cryo-EM map. At a contour level of the predicted map where half of the high-density prediction is wrong (precision = 0.5), GuangzhouRNA recovered 14% of the reference voxels, bussilab_plain_md 10%, 9CBX 26%, 9CBU 15%, and the independent map 83%. However, since the single model ensembles of GuangzhouRNA and the reference atomic models accounted for so little of the density, they hit a recall plateau and larger ensembles like that of bussilab_plain_md attained higher recall at contours with lower precisions. At a contour of the predicted map at which two-thirds of the predictions were allowed to be incorrect, GuangzhouRNA recovered 22% of the reference voxels, bussilab_plain_md 31%, 9CBX 33%, 9CBU 20%, and the independent map 92%.

**Figure 5:**
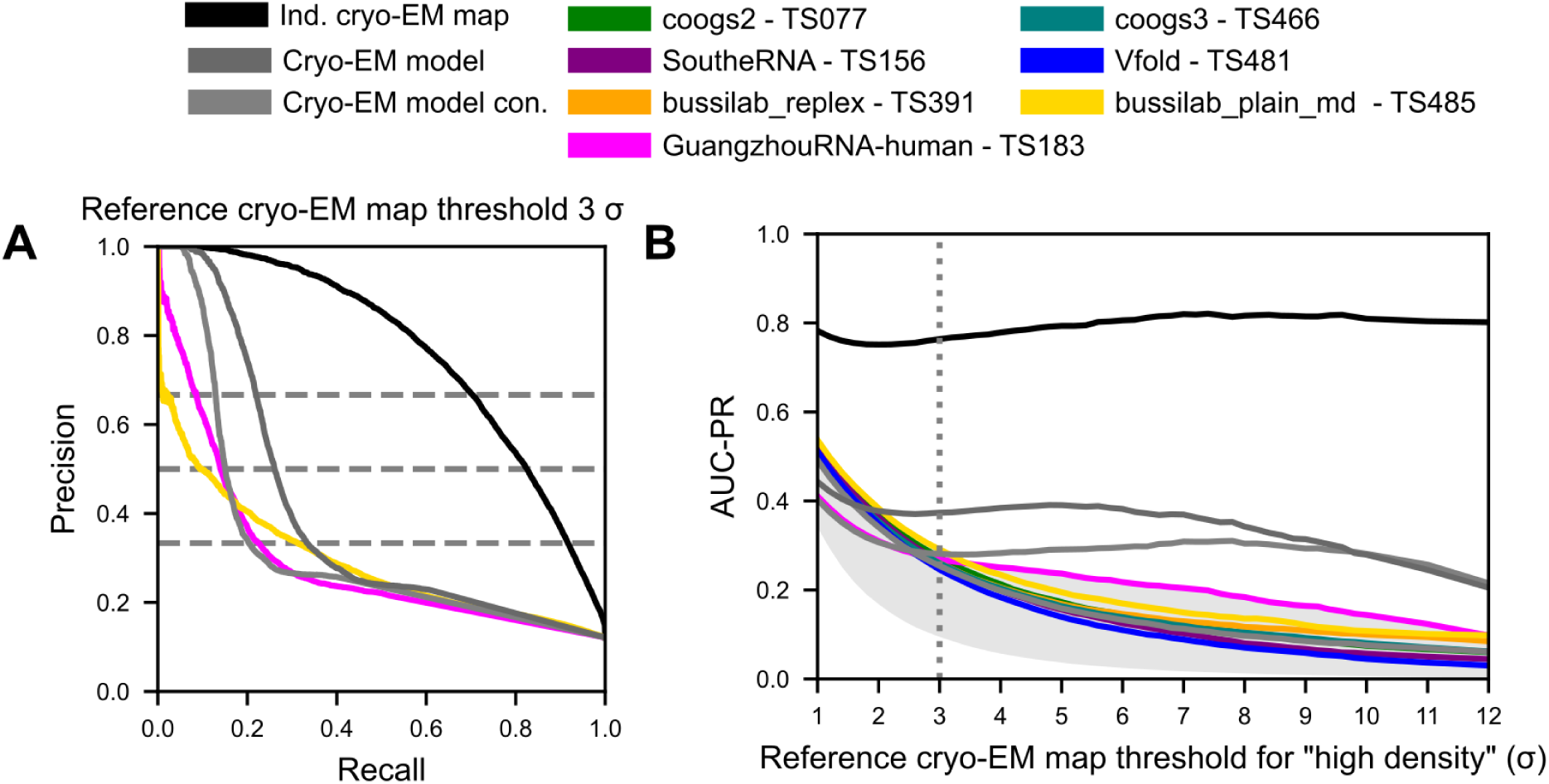
Precision-recall curve of predictions. (**A**) The precision-recall curve for predicting the voxel of high density in the cryo-EM map (> 3 σ) is calculated by varying the contour of the predicted density. (**B**) The area under the precision-recall curve (AUC-PR) is calculated for various thresholds of the reference cryo-EM map.

We additionally measured AUC-PR at various thresholds of the reference map to observe how accuracy changed when the focus was set on more or less ordered water and ions from the reference. The independent cryo-EM map and cryo-EM derived atomic model all had relatively stable AUC-PR across thresholds, but the MD methods all fell in AUC-PR with increasing levels of order, suggesting that they were not accurately capturing the most ordered water and ion positions (**Figure 5B**). The AUC-PR of GuangzhouRNA decreased with reference cryo-EM map threshold at a slower rate than other predictions, suggesting it was able to retrieve more accurate binding positions for highly ordered water and ions than MD simulations. We note that this analysis should not be used to interpret the identity (i.e., ion type) of the positions (see **Discussion**).

### 3.7 Prediction methodology and performance

The groups were ranked using a Z-score based ranking to combine all four metrics described above. Five groups ranked above baseline: bussilab_plain_md, SoutheRNA, bussilab_replex, coogs2, and coogs3 (**Figure 4**). These groups conducted MD simulations carried out near room temperature and sampled the simulation to submit a large ensemble. A variety of buffer conditions and box sizes were used, with the Bussi Lab entries having the largest box size. These groups simulated a range of time scales from a few ns to a few μs, suggesting that simulation time was not a differentiating factor for performance. This result also suggested that water and ion occupancy prediction was not improved with longer simulations, even though with increased force field accuracy in the future, longer simulations or enhanced sampling may eventually be important to sample ion occupancy. Additionally, ensembles on the order of 100 models appeared sufficient because top models ranged from 100 to 1000 models in the ensemble, and subsampling the baseline “addIons” model to smaller ensemble sizes led to saturating performance at and above 100 frames. Small ensembles were generally submitted by groups that aimed to predict a small selection of well-ordered water and ions well, a complementary but distinct task from this ensemble modeling challenge (**Box 1**). Hence, these small ensembles were not able to accurately predict the density fully. Unexpectedly, the top groups that provided large ensmebles used a diverse set of force fields, including diverse water and ion parameterizations (**Table 1**). This observation indicated either that our analysis was not able to significantly differentiate the accuracy of force fields or that other simulation variables are more important. Of note, the MD baselines, which had 3 different force field-ion model combinations but held other factors constant, were not significantly different from each other in accuracy. Additionally, the analysis by the SoutheRNA group showed that the choice of water model did not significantly affect their results (**Supplemental Text 15**).

Although this study focused on water and ion structures, we noted that changes in RNA conformation could influence the local water and ion ensembles. Hence, we examined the top groups’ treatment of RNA conformational fluctuations, which could be influenced by the starting structure, equilibration, and restraints. The starting model used in all cases was 7EZ0^44^ except for SoutheRNA which started from an AlphaFold3 based model (4 Å RMSD from 7EZ0; **Supplemental Text 4**). Many other MD groups also started from 7EZ0, suggesting the starting structure was not the major determinant of performance.

Some of the top groups restrained the RNA while others did not. However, even the unrestrained simulations of these top groups resulted in ensembles close, relative to other MD groups, to the reference RNA structure (**Supplemental Figure 3**). In fact, for groups who submitted large ensembles, performance was anti-correlated to the average global RMSD to the reference (**Figure 6A**). This result suggested that unrestrained simulations can rival restrained simulations for described water and ion structure, but only if they are sufficiently stable. A stable simulation requires proper placement and equilibration of cations to prevent the unfolding effect of backbone electrostatic repulsion, including maintaining known Mg^2+^ ions, which was done by all top groups. Some groups removed the Mg^2+^ ions in the starting 7EZ0 model (**Supplemental Text 10**), which may have led to instability or sampling of alternative structures, leading to a difference in performance despite similar simulation methodologies.

**Figure 6:**
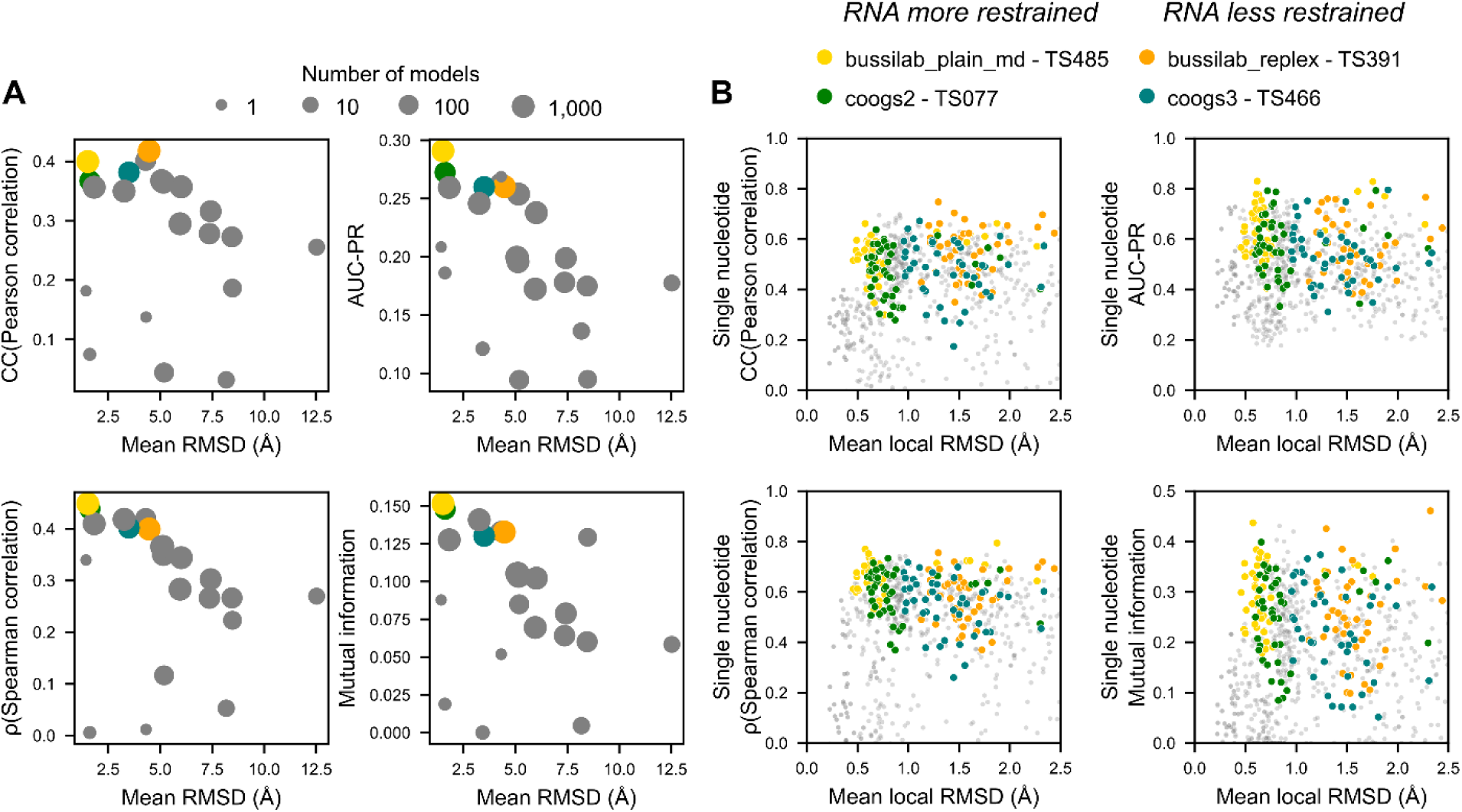
Variation in scoring with deviations in RNA structure. (**A**) For every group, the global mean all heavy atom RNA RMSD of all models submitted is plotted against various metrics measuring agreement to the reference cryo-EM density for density_scat_ in a 1.8-3.2 Å shell around the RNA. Four groups of interest are highlighted with colored dots; the remaining groups are gray. Symbol sizes convey the number of models submitted. (**B**) For every residue and group, the mean local all heavy atom RNA RMSD of all models submitted is plotted against various metrics measuring agreement to the reference cryo-EM density for density_scat_ in the region of the 1.8-3.2 Å shell around the RNA that is closest to the nucleotide. Only the nucleotides of highest experimental confidence (N = 48), nucleotides with top (80%) scores for all four metrics in the independent cryo-EM map are considered. A similar trend is observed for all residues. For visualization, the mean local RMSD is cut at 2.5 Å; the flat trend continues even at higher RMSD values.

However, our data indicated that restraints still play a role in accuracy. When comparing restrained and less restrained simulation from the same research laboratories (bussilab_plain_md vs. bussilab_replex and coogs2 vs. coogs3), we saw slightly better performance for the restrained simulations, implying a modest advantage to restrained simulations. This pattern is consistent with cryo-EM capturing a dominant RNA conformation whose associated water and ion ensemble is not preserved when the RNA deviates substantially in solution. An alternative explanation is that deviations in RNA structure interfered with our density-generation method. If so, we would expect, within the same group, lower scores in regions where the RNA was locally aligned less accurately, discussed in the next section.

Finally, the sampling strategy can influence the RNA conformations selected for the ensemble. The majority of groups, including all top groups, used uniform sampling across simulation time to select frames, which would reduce the likelihood of selecting rare RNA conformations. The alternative sampling strategy, K-means clustering, was used by GromihaLab and Cheatham Lab groups. This clustering strategy selects frames that have distinct RNA conformations; this resulted in an ensemble where the RNA diverged significantly from the cryo-EM structure (**Supplemental Figure 2**). This divergence in RNA structure, and hence potentially the water and ion structure, could explain the relatively poorer performance of these groups.

Overall, RNA restraints, ionic composition, initial ion placement, sampling strategy, and ionic equilibration appear to be important variables and are worthy of future investigation.

### 3.8 Local accuracy measurements

As described in **Methods**, we applied the same comparative scores to local regions around individual nucleotides (**Figure 7A**). When comparing the independent cryo-EM map to the reference cryo-EM map, we observed good correlation between these metrics (**Supplemental Figure 9A**). Mutual information was correlated to the Pearson and Spearman correlation but was more expressive (larger range of values) for nucleotides with high scores, suggesting that it may be an important metric for discriminating high accuracy predictions in the future. AUC-PR exhibited the weakest correlation to other metrics, perhaps because, with the small number of voxels around each nucleotide, the discretized nature of AUC-PR reduced its expressivity. Similar trends were observed for the comparisons of the predicted ensemble to the SWIM reference cryo-EM map (**Supplemental Figure 9B**), suggesting that these are robust metrics.

**Figure 7:**
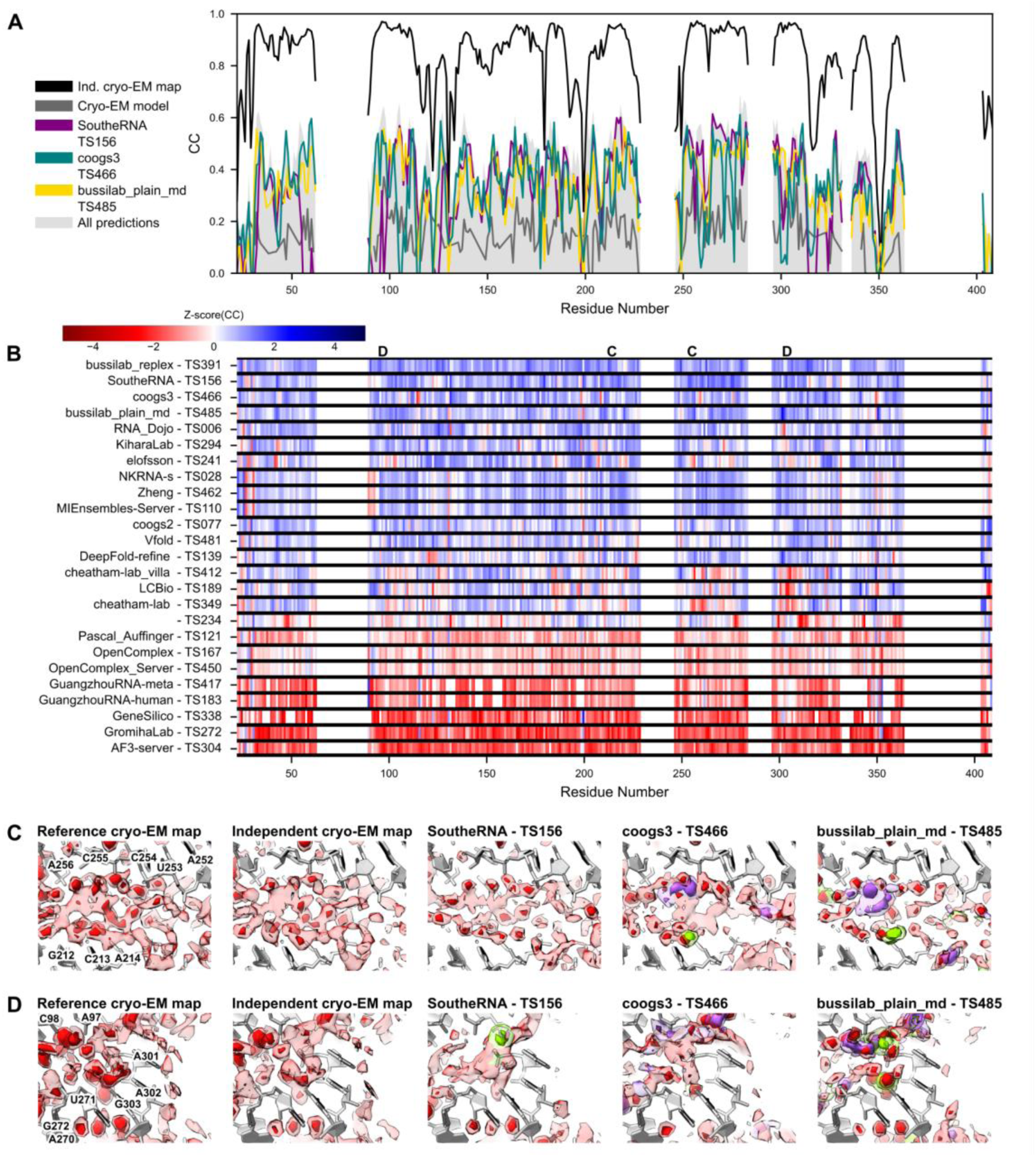
Correlation of predicted ensemble and cryo-EM map in the solvent shell around each nucleotide. (**A**) The Pearson correlation (CC) is shown for predicted water and ion density based on scattering factors in the solvent shell surrounding every residue. For residues poorly resolved in the cryo-EM map, the data is not shown. The shaded gray area is the range of scores for all CASP16 predicting groups. The black line represents the “ceiling” of each residue CC; the CC between the independent cryo-EM map (EMD-42498) and the reference map (EMD-42499). (**B**) For every residue, the Z-score of Pearson correlation from all predicting groups is displayed from red (below average performance) to blue (above average performance). The regions of focus in (C,D) are labeled at the top of the plot. The groups are sorted by the sum over all residues of Z-score(CC) > 0. (**C,D**) The local agreement between cryo-EM density and predicted water and ion density is shown for two regions. The cryo-EM maps are displayed at 2 σ (light red) and 5 σ (dark red) – the molecular identity for any peaks is not assumed. For the predicted models, features with low density (light color) and high density (dark color) are shown for water (red), Mg^2+^ ions (green), Na^+^ ions (purple), and Cl^-^ ions (yellow). See the density for all prediction groups, references, and baselines in **Supplemental Figure 4** and **Supplemental Figure 5**.

These measures of local agreement to the cryo-EM map were not correlated with local RMSD (**Figure 6B**), hence, RNA deviations are not influencing the scoring significantly. Instead, in simulations, these diverse RNA conformations appear to have distinct water and ion ensembles that do not match the cryo-EM data as well as simulated water and ion ensembles when the RNA structure is close to the reference.

We additionally used these local measurements to explore origins of differences in global scores between methods. For each metric and nucleotide, we computed the Z-scores across groups to quantify how far each group’s score deviated from the mean. Beyond the previously described difference in AUC-PR, there were no noticeable differences in scoring: groups that performed significantly above average on one metric or density-generation method tended to do so on the others (**Supplemental Figure 10**). This result suggested that global differences among correlation-based scores were largely due to overall simulation accuracy and not due to difference in specific idiosyncratic regions in the *Tetrahymena* ribozyme molecule. This conclusion was limited by current model accuracy and should be revisited when correlations approach experimental uncertainty.

The Z-score plot also highlighted regions for visual inspection (**Figure 7B**). For example, the solvent shell around residues 212-214 and 252-256 is a region where the top MD groups mentioned above outperformed other groups. This region is the junction between P4 and P6, near the catalytic core, where a network of ∼3 water molecules wide stitches together the junction. The cryo-EM density has a complex shape showing diffuse density with a few ordered (dense) features (**Figure 7C**). The independent cryo-EM map agrees well, although it has fewer well-resolved dense peaks (**Figure 7C**). SoutheRNA, coogs3, and bussilab_plain_md all predicted density similar to the cryo-EM map; notably, SoutheRNA’s density arose predominantly from water molecules, whereas coogs3 and bussilab_plain_md include coordinated cations (**Figure 7C**). This comparison revealed a limitation of density-only analysis: the molecular identity of cryo-EM map features can be ambiguous, and predictions that differ in what molecules produce the density cannot currently be discriminated.

Despite limited confidence in assigning molecular identity from a 2.2 Å map alone, our comparative analysis suggested that the cryo-EM density still carried clues to identity. For example, around residue 302, bussilab_plain_md performed better than other groups (**Figure 7B**). Examining the cryo-EM density we saw an octahedral shaped density suggesting that this feature was an ion, likely Mg^2+^, coordinating six waters at each terminus of the octahedron and binding RNA indirectly in its second coordination shell (**Figure 7D**). This region is another junction in the catalytic core, between P3 and P6, where water and ion structure mediate the junction interaction. The bussilab_plain_md group was the only group to predict a Mg^2+^ with high occupancy at this location, hence leading to a higher score (**Figure 7D**).

## 4 Discussion

This study demonstrates the potential of a cryo-EM density-based comparison to assess the accuracy of predicted water and ion structures. These comparisons provide a new avenue for describing the inherently flexible and complex ensemble of water and ions around macromolecules, which could complement the highly ordered water and ions described in single-structure PDB depositions. The data and predictions are all openly available as a resource for future explorations. Scoring metrics introduced here should be further developed, including considering map sharpening, power spectrum matching, filtering of noise, and alternative metrics. This community-wide study should be viewed as a pilot for comparisons of prediction methodologies and evaluating the utility of cryo-EM data and predictions for answering biophysical questions. Further blind challenges would open the door towards exploring more functionally motivated questions regarding water and ion ensembles, such as how they change upon ligand binding or though the catalytic cycle in this ribozyme or in other macromolecules.

The top-performing groups all used MD but employed different force fields and different water and ion parameterizations. There were a multitude of decisions that varied between these groups and affected the MD simulation, such as initial placement, box size, and equilibration. In particular, the number of Mg^2+^ ions and the ratio of divalent to monovalent ions, varied widely, as shown in **Table 1**. A correlation between lower deviations in RNA structure and higher agreement with solvent shell density highlights an important future direction of exploring the change in water and ion networks upon conformational changes or ligand binding. However, overall, no single parameter was shared by the top-performing groups, suggesting that at this level of accuracy, no individual parameter is the primary determinant of accuracy. With so many variables differing between groups, this study of 26 methods could not disentangle these multivariable dependencies. To compare methodologies at this level of accuracy, a systematic study varying one factor at a time will be needed. We also note that the majority of participating groups used MD approaches; more outreach to groups developing AI or hybrid approaches should be carried out in future challenges on water and ion ensemble prediction.

For the expansion beyond MD approaches, it is important to differentiate between the task posited here, prediction of solvent shell occupancy, and the prediction of a select set of ordered water and ions (e.g., ions and waters found in the PDB training data; see also ref^45^). AI algorithms have been developed to address the latter prediction tasks and could be subjects of an interesting complementary assessment similar to CASP ligand assessment with the potential addition of map fit metrics or geometric considerations^7^. This would enable the comparison of ion placement from deep learning algorithms like AlphaFold3 and MgNet (**Supplemental Text 12**) and more classical algorithms like 3D-RISM (**Supplemental Text 1**). In theory, these methods could be expanded to predict a large ensemble of ion and water placements, adapting them to the prediction task assessed here. The AlphaFold3 -server did participate in this challenge (**Supplemental Text 6**) and has been used as a baseline in other CASP challenges; however, the AlphaFold3-server was only used to predict the RNA conformation in this challenge. Na-HEPES and MgCl_2_ were randomly placed as ion clusters, not fully solvated separate ions, and then the system was solvated with water. Hence, while the AlphaFold3server performed poorly, the challenge did not assess the ability of the AlphaFold3-server to place ions. All methods, except for AlphaFold3, used time-evolving simulations, molecular dynamics, or the methods of GeneSilico (**Supplemental Text 9**). Machine-learning methods have not been developed specifically for this ensemble-modeling task due to the lack of training data. Potentially, the cryo-EM data can be utilized to train such algorithms in the future.

It is important to recognize the limitations and assumptions from the experimental data, as these affect the interpretation of the predictions. There are three predominant factors: radiation damage, the freezing process, and ambiguity of molecular identity. Radiation damage is a major concern because water and ion peaks are among the first densities to be affected in the initial frames. Bulk ions rapidly diffuse during collection^46^, but the effects of radiation on the molecules more tightly associated with the biomolecule are less well understood. A frame-by-frame analysis of damage and a dose-weighting procedure specifically tuned to consider water and ions may be important for future studies. Unfortunately, the cryo-EM data used herein were collected on carbon grids with high levels of beam-induced motion in the first few frames, so these data are ill-suited for that question. Collecting high-resolution data on HexAuFoil^47,48^ to obtain high-quality signals in the first frames of data collection will be important for investigating water and ions.

Water and ion ensembles are likely affected by the millisecond-scale freezing process, which is thought to influence dynamics with energy barriers less than 2.4 kcal mol^-1^ (10 kJ mol^-1^)^49^. Many ordered ions may not be affected significantly, but this could perturb the dynamics of water and ions in the second hydration shell. These effects could be investigated through simulation, such as the simulations at cryogenic temperature conducted by Pascal_Auffinger here. Additionally, these effects could be investigated in the context of rapid beam-induced melting and re-vitrification of the sample to approach a more liquid-like environment^50,51^.

Finally, this study was limited in its ability to discriminate between different models’ predictions for molecular identity. While predictions offered a detailed picture of the ion and water structure (e.g., identifying where Na^+^ was predicted to bind vs. where Mg^2+^ binds), the cryo-EM map is, at its root, a “black-and-white” picture. A view that distinguishes water and ions from each other is vital in investigating the ionic sensitivity of biomolecular function, which is commonly observed in RNA elements. Cryo-EM is not inherently blind to molecular identity. In fact, because cryo-EM maps represent electrostatic potential, they are sensitive to ionic charge^52–54^, but these effects are currently not well-modeled. Better modeling of this charge sensitivity, combined with coordination geometry enabled by higher resolution data, could pave the way towards more descriptive cryo-EM maps and more incisive testing and refinement of water and ion ensemble predictions.

The gold standard measure of predictive utility is whether predictions help answer biophysical questions. The impact of water and ion structure on questions such as relative small-molecule binding affinities remains poorly quantified. Challenges such as the one put forth in this study enable the investigation and quantification of water and ion structure. Future work should expand upon the ideas here towards determining the functional relevance of water and ion structure. For example, future high-resolution structures of the *Tetrahymena* ribozyme captured at different stages of its catalytic cycle could improve our ability to study and accurately predict changes in water and ion occupancy during catalysis.

## Data availability

Predictions can be downloaded from https://predictioncenter.org/download_area/CASP16/predictions/R1260/. Additional abstract describing group 121 - Pascal_Auffinger and 139 - DeepFold-refine can be found at https://predictioncenter.org/casp16/doc/CASP16_Abstracts.pdf. Volumes produced from the predicted ensembles and baseline ensembles can be accessed at https://doi.org/10.25740/vz022mq8177 and the models can be accessed at PDB 9CBU, 9CBW, 9CBX, and 9CBY^22^. The cryo-EM maps compared to in this study can be found at EMDB 42498 and 42499^22^. Code and scripts used in this study can be found at https://github.com/DasLab/CASP16_NA/tree/main/R1260_assessment.

## Author contributions

R.C.K., D.A.C., W.C., and R.D. conceptualized and designed the study. D.A.C, W.C., R.D., S.L., M.Z.P., and K.Z. determined the cryo-EM maps used in this study and advised on the creation of this challenge. All other authors made blind predictions of the water and ion solvation, contributed a methods section describing these predictions, and, in some cases, provided a supplemental text describing their analysis; the contributing authors are listed in the title of each section. R.C.K., E.P., D.A.C., and R.D. analyzed results. R.C.K. compiled the manuscript with input from all authors.

## Acknowledgments

We thank the CASP organization for enabling this pilot challenge, particularly Andriy Kryshtafovych for organizing the collection of these large ensembles. We thank Alissa Hummer for feedback on the manuscript. This work was supported, including the use of compute resources, by: Berzelius supercomputing resource (provided by the National Supercomputer Centre at Linköping University, the Knut and Alice Wallenberg Foundation, and SNIC, grant Nos. SNIC 2021/5-297 and Berzelius-2021-29 to A.E.); Bio-X Bowes Graduate Student Fellowship (R.C.K.); Cancer Prevention and Research Institute of Texas (CPRIT RR220008 to G.H.Z.); Core Research for Evolutional Science and Technology (CREST JPMJCR21F1 to M.H.); EURO-HPC (2023R03-136 to G.B., O.L.C.,and E.P.); European Molecular Biology Organization (ALTF 525-2022 to E.F.B.); European Union’s Horizon 2023 research and innovation programme (Marie Skłodowska-Curie 101152924 to O.L.C.); Financiamiento Basal para Centros Científicos y Tecnológicos de Excelencia de ANID (S.P.); Fondecyt Regular (1231071 to S.P.); Foundation for Polish Science and the EU European Regional Development Fund (POIR.04.04.00-00-3CF0/16 to J.M.B.); Fundamental Research Funds for the Central Universities (054-63253109 to W.Z.); Hewlett-Packard Enterprise Data Science Institute maintained computational resources at the University of Houston (G.H.Z.); Howard Hughes Medical Institute (to R.D.); Italian National Centre for HPC, Big Data, and Quantum Computing (CN00000013 to G.B.); Japan Agency for Medical Research and Development (AMED 23ae0121049h0003 to M.H.); Knut and Alice Wallenberg Foundation (A.E.); Next Generation EU initiative through PRIN 2022 (2022Z4FZE9 to G.B. and O.L.C.); National Institute of General Medical Sciences (T32 GM132024 to J.V.); National Institutes of Health (R01GM079429 to W.C., U24GM129541 to W.C., R35GM122579 to R.D., R35-GM134919 to S.J.C., U54-AI170660 to S.J.C., R01-GM-081411 to T.E.C., R01GM133840 and R35GM158267 to D.K.); National Natural Science Foundation of China (12426303 to W.Z., 92370128 to G.H., 12326611 to G.H.); National Science Centre, Poland (SHENG 2021/40/Q/NZ2/00078 to S.K. and 2017/26/A/NZ1/01083 to J.M.B.); National Science Foundation (2330652 to W.C. and R.D., CBET-2442006 to G.H.Z., CHE-2154924 to S.J.C., IIS2211598 to D.K., DMS2151678 to D.K., DBI2003635 to D.K., DBI2146026 to D.K.); Polish high-performance computing infrastructure PLGrid (HPC Center: ACK Cyfronet AGH PLG/2024/016931 to C.N.); Research Center for Computational Science, Okazaki, Japan (24-IMS-C123 to M.H.); SLAC Shared Scientific Data Facility (R.C.K.); Stanford Research Computing Center (R.C.K.); Tianjin Science and Technology Program (24ZXZSSS00320 to G.H. and W.Z.); Vetenskapsrådet (2021-03979 to A.E); Welch Foundation (E-2221 to G.H.Z., Catalyst Center for Advanced Bioactive Materials Crystallization V-E-0001 to G.H.Z.). This article is subject to HHMI’s Open Access to Publications policy. HHMI lab heads have previously granted a nonexclusive CC BY 4.0 license to the public and a sublicensable license to HHMI in their research articles. Pursuant to those licenses, the author-accepted manuscript of this article has been made freely available under a CC BY 4.0 license on BioRxiv.

