## Supplemental Figures, Tables, and Text for "Blind prediction of complex water and ion ensembles around RNA in CASP16"


[1]Rachael C. Kretsch https://orcid.org/0000-0002-6935-518X
[2]Elisa Posani https://orcid.org/0000-0002-0217-5125
[3,4]Eugene F. Baulin https://orcid.org/0000-0003-4694-9783
[3]Janusz M. Bujnicki https://orcid.org/0000-0002-6633-165X
[2]Giovanni Bussi https://orcid.org/0000-0001-9216-5782
[5]Thomas E. Cheatham III. https://orcid.org/0000-0003-0298-3904
[6,7,8]Shi-Jie Chen https://orcid.org/0000-0002-8093-7244
[9]Arne Elofsson https://orcid.org/0000-0002-7115-9751
[3]Masoud Amiri Farsani https://orcid.org/0000-0001-5116-0483
[5]Olivia N. Fisher https://orcid.org/0000-0002-0862-8367
[10]M. Michael Gromiha https://orcid.org/0000-0002-1776-4096
[11]Ayush Gupta https://orcid.org/0009-0006-1702-6946
[12,13]Michiaki Hamada https://orcid.org/0000-0001-9466-1034
[10]K. Harini
[14]Gang Hu https://orcid.org/0000-0002-7134-3380
[15]David Huang
[16]Junichi Iwakiri https://orcid.org/0000-0002-4611-2142
[15]Anika Jain https://orcid.org/0000-0002-0249-875X
[15]Yuki Kagaya https://orcid.org/0000-0003-0146-1709
[15,17]Daisuke Kihara https://orcid.org/0000-0003-4091-6614
[18]Sebastian Kmiecik https://orcid.org/0000-0001-7623-0935
[10]Sowmya Ramaswamy Krishnan https://orcid.org/0000-0001-5404-3266
[19]Ikuo Kurisaki https://orcid.org/0000-0003-4519-1093
[2]Olivier Languin-Cattoën https://orcid.org/0000-0002-0486-9841
[20]Jun Li https://orcid.org/0000-0002-7011-1991
[21]Shanshan Li https://orcid.org/0000-0002-7041-5960
[11]Karim Malekzadeh https://orcid.org/0009-0004-8185-4995
[15]Tsukasa Nakamura https://orcid.org/0000-0002-6312-3070
[14]Wentao Ni https://orcid.org/0009-0002-7355-0073
[18]Chandran Nithin https://orcid.org/0000-0001-8212-6093
[22]Michael Z. Palo https://orcid.org/0000-0003-4883-6766
[17]Joon Hong Park
[18]Smita P Pilla https://orcid.org/0000-0002-4520-3210
[23,24]Simón Poblete https://orcid.org/0000-0001-7793-0570
[25]Fabrizio Pucci https://orcid.org/0000-0003-2916-022X
[15]Pranav Punuru

[26]Anouka Saha
[27]Kengo Sato https://orcid.org/0000-0001-6744-7390
[10]Ambuj Srivastava
[15]Genki Terashi https://orcid.org/0000-0002-5339-909X
[15]Emilia Tugolukova
[15]Jacob Verburgt https://orcid.org/0000-0002-8524-1342
[28]Qiqige Wuyun https://orcid.org/0000-0002-7228-903X
[11]Gül H. Zerze https://orcid.org/0000-0002-3074-3521
[21]Kaiming Zhang https://orcid.org/0000-0003-0414-4776
[6]Sicheng Zhang https://orcid.org/0009-0009-9851-5451
[14]Wei Zheng https://orcid.org/0000-0002-2984-9003
[6]Yuanzhe Zhou https://orcid.org/0000-0003-1858-3067
[1,29,30,31,*]Wah Chiu https://orcid.org/0000-0002-8910-3078
[32]David A. Case https://orcid.org/0000-0003-2314-2346
[1,33,34,*]Rhiju Das https://orcid.org/0000-0001-7497-0972

[1]Biophysics Program, Stanford University School of Medicine, Stanford, California, USA.
[2]Scuola Internazionale Superiore di Studi Avanzati (SISSA), via Bonomea 265, 34136 Trieste, Italy.
[3]Laboratory of Bioinformatics and Protein Engineering, International Institute of Molecular and Cell Biology in Warsaw, ul. Ks. Trojdena 4, 02-109 Warsaw, Poland.
[4]Laboratory of RNA Algorithms, IMol Polish Academy of Sciences, ul. M. Flisa 6, 02-247 Warsaw, Poland.
[5]University of Utah, Medicinal Chemistry Department, Salt Lake City, Utah, USA.
[6]Department of Physics and Astronomy, University of Missouri, Columbia, MO 65211, USA.
[7]Department of Biochemistry, University of Missouri, Columbia, MO 65211, USA.
[8]MU Institute for Data Science and Informatics, University of Missouri, Columbia, MO 65211, USA.
[9]Dep of Biochemistry and Biophysics and Science for Life Laboratory, Stockholm University Box 1031, 171 21 Solna, Sweden.
[10]Department of Biotechnology, Bhupat and Jyoti Mehta School of Biosciences, Indian Institute of Technology Madras, Chennai 600036, Tamil Nadu, India.
[11]William A. Brookshire Department of Chemical and Biomolecular Engineering, University of Houston, 4226 Martin Luther King Boulevard, Houston, TX, 77204, USA.
[12]Faculty of Science and Engineering, Waseda University, 169-8555, Tokyo, Japan.
[13]AIST-Waseda University Computational Bio Big-Data Open Innovation Laboratory (CBBD-OIL), National Institute of Advanced Industrial Science and Technology, 169-8555, Tokyo, Japan.
[14]NITFID, School of Statistics and Data Science, AAIS, LPMC and KLMDASR, Nankai University, Tianjin 300071, China.
[15]Department of Biological Sciences, Purdue University, West Lafayette, IN, 47907, USA.
[16]Graduate School of Frontier Sciences, The University of Tokyo, 277-8561, Chiba, Japan.
[17]Department of Computer Science, Purdue University, West Lafayette, IN, 47907, USA.

[18]University of Warsaw, Biological and Chemical Research Centre, Faculty of Chemistry, Warsaw, Poland.
[19]Waseda Research Institute for Science and Engineering, Waseda University, 169-8555, Tokyo, Japan.
[20]School of Sciences, Great Bay University, Great Bay Institute for Advanced Study, Guangdong Provincial Key Laboratory of Mathematical and Neural Dynamical Systems, Dongguan, Guangdong 523000, China.
[21]Division of Life Sciences and Medicine, University of Science and Technology of China, Hefei 230001, China.
[22]Department of Structural Biology, Stanford University School of Medicine, CA USA.
[23]Facultad de Ingeniería, Universidad San Sebastián, Bellavista 7, 8420524 Santiago, Chile.
[24]Centro BASAL Ciencia & Vida, Universidad San Sebastián, Av. del Valle Norte 725, 8580704 Santiago, Chile.
[25]Computational Biology and Bioinformatics, Université Libre de Bruxelles, Brussels, Belgium.
[26]Department of Mathematics, The University of Texas at Austin, Austin, TX, 78712, USA.
[27]School of Life Science and Technology, Institute of Science Tokyo, 152-8550, Tokyo, Japan.
[28]Department of Computer Science and Engineering, Michigan State University, East Lansing, Michigan 48824, USA.
[29]Department of Bioengineering and James Clark Center, Stanford University , CA USA.
[30]Department of Microbiology and Immunology, Stanford University School of Medicine, CA USA.
[31]Division of CryoEM and Bioimaging, SSRL, SLAC National Accelerator Laboratory, Menlo Park, CA, USA.
[32]Department of Chemistry & Chemical Biology, Rutgers University, Piscataway, New Jersey, USA.
[33]Department of Biochemistry, Stanford University School of Medicine, CA USA.
[34]Howard Hughes Medical Institute, Stanford University, CA USA.

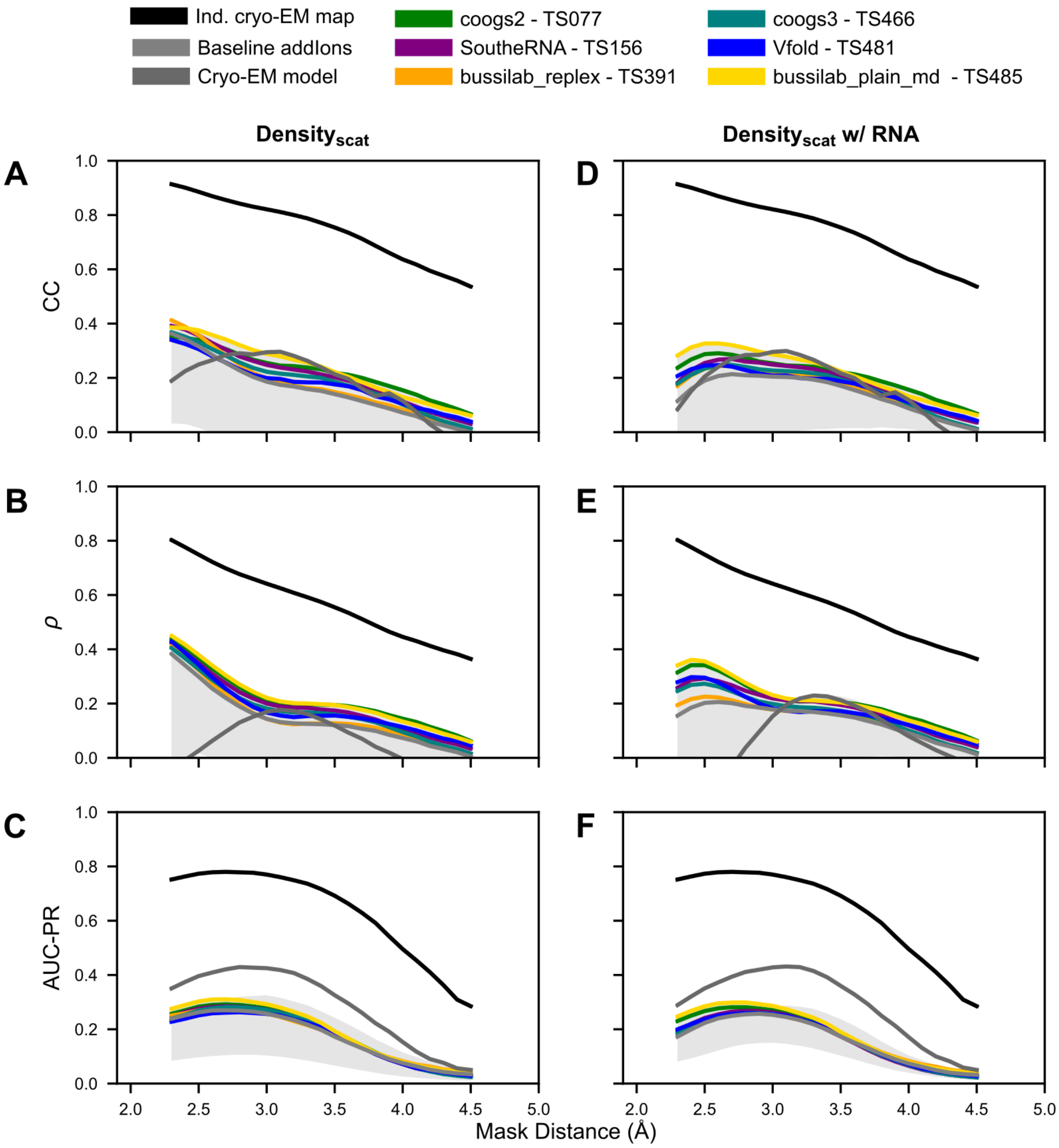


**Supplemental Figure 1: Prediction accuracy for 1 Å windows of the solvent shells.** The shaded grey area represents the range of score for all CASP16 predictors.

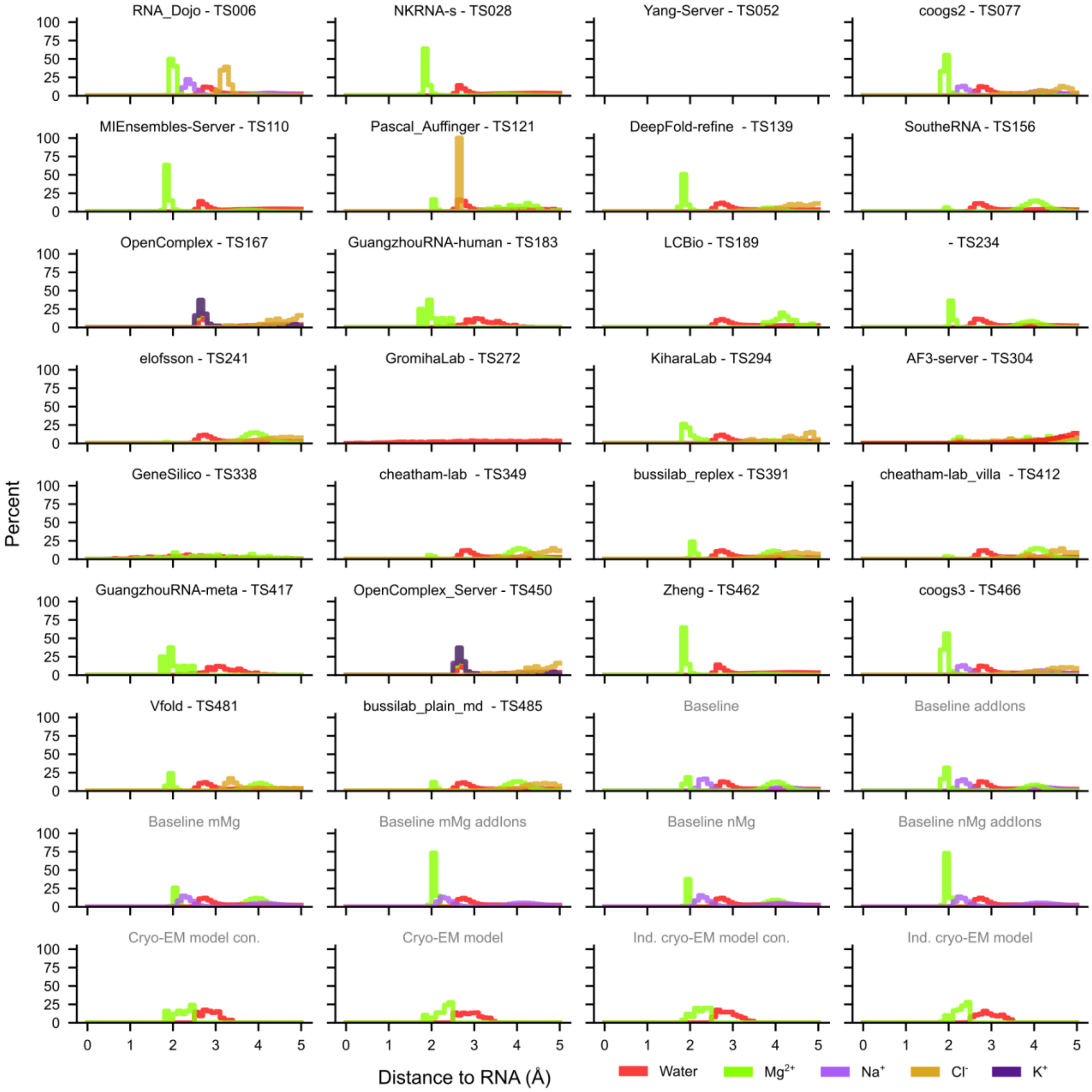


**Supplemental Figure 2: Proximity of water and ions to RNA.** For each ensemble, the distance between the water or ion and the closest RNA heavy-atom is calculated and the distribution of distances is displayed in histograms after binning by 0.1 Å. Water is colored red, $Mg^{2+}$ ions are green, $Na^{+}$ are light purple, $Cl^{-}$ are dark yellow, and $K^{+}$ are dark purple. Flat histograms with values near 0% across the distance range indicate that the molecules do not have a modal distance to RNA in the ensemble; they are evenly distributed at all distances from the RNA.

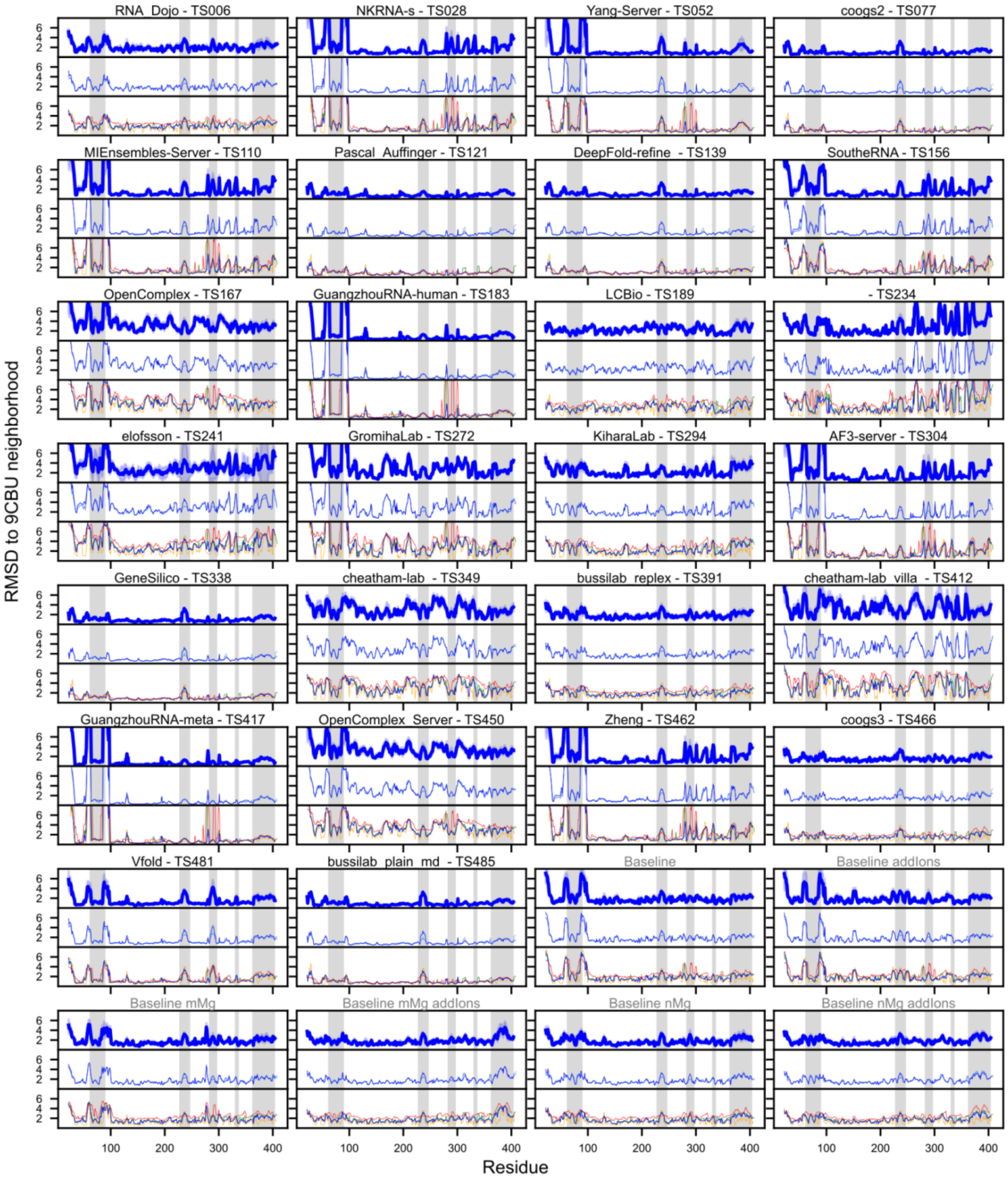


**Supplemental Figure 3: Local alignment of RNA residues.** For each model, each residue, including its local neighborhood, was aligned to the reference separately. The mean RMSD for each residue's local alignment is displayed for every group's ensemble. On the top in dark blue is the main alignment – a 10 Å neighborhood aligning with all RNA heavy atoms. The width of the line shows the standard deviation across models submitted. The second graph displays the mean local RMSD when the alignment is done with all heavy atoms, select 5 atoms, select 3 atoms, or just backbone atoms. These lines overlap as these alignment methods do not produce significantly different results. The bottom plot shows the local RMSD when aligning on a 6, 10, 12, or 20 Å neighborhood around the residue.

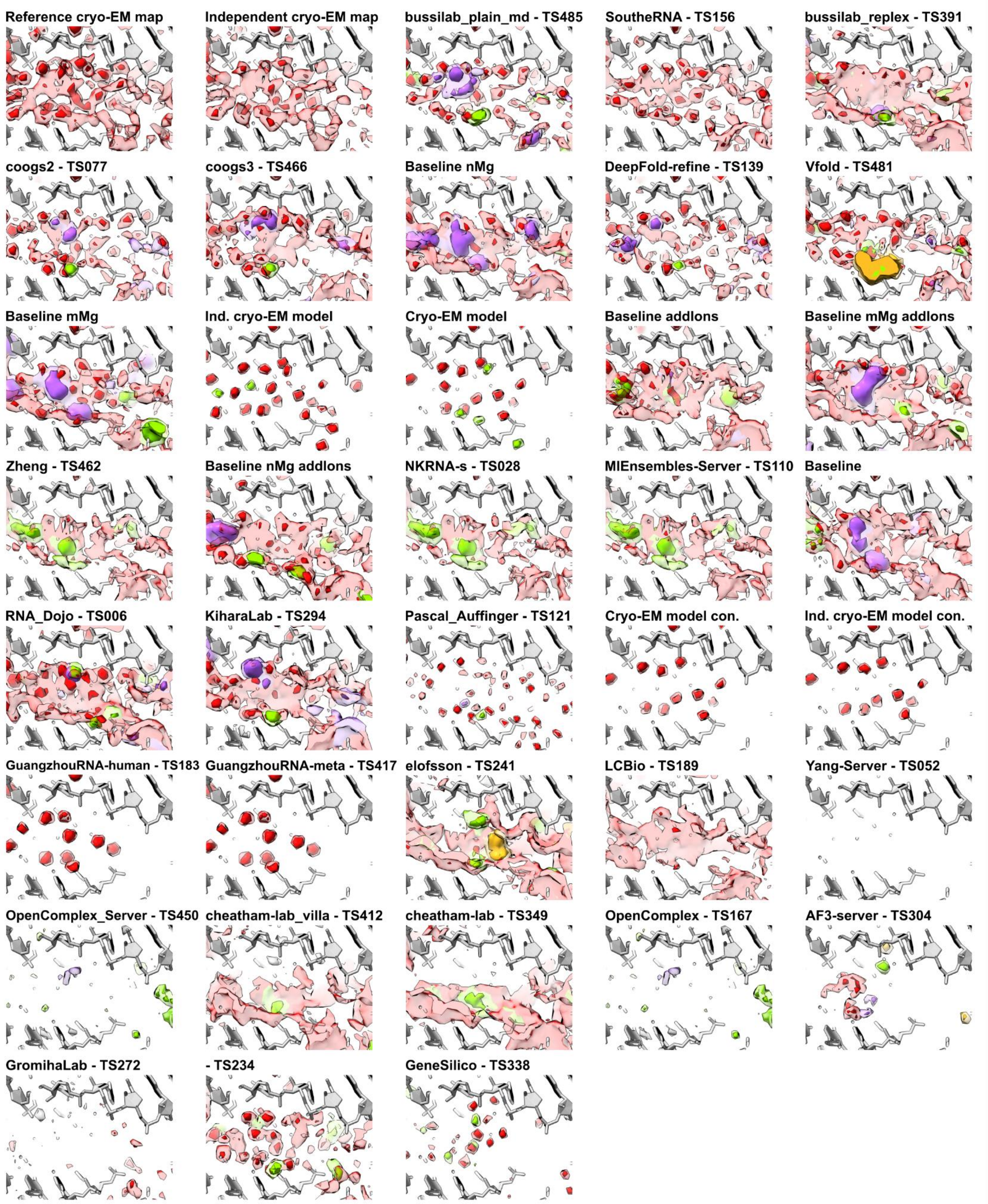


**Supplemental Figure 4: The local agreement between cryo-EM density and predicted water and ion density is shown for regions in Figure 7C.** The cryo-EM maps are displayed at 2 σ (light red) and 5 σ (dark red) – the molecular identity for any peaks is not assumed. For the predicted models, features with low density (light color) and high density (dark color) are shown for water (red), $Mg^{2+}$ ions (green), $Na^{+}$ ions (purple), and $Cl^{-}$ ions (yellow). The groups are ordered by ranking in **Figure 4**.

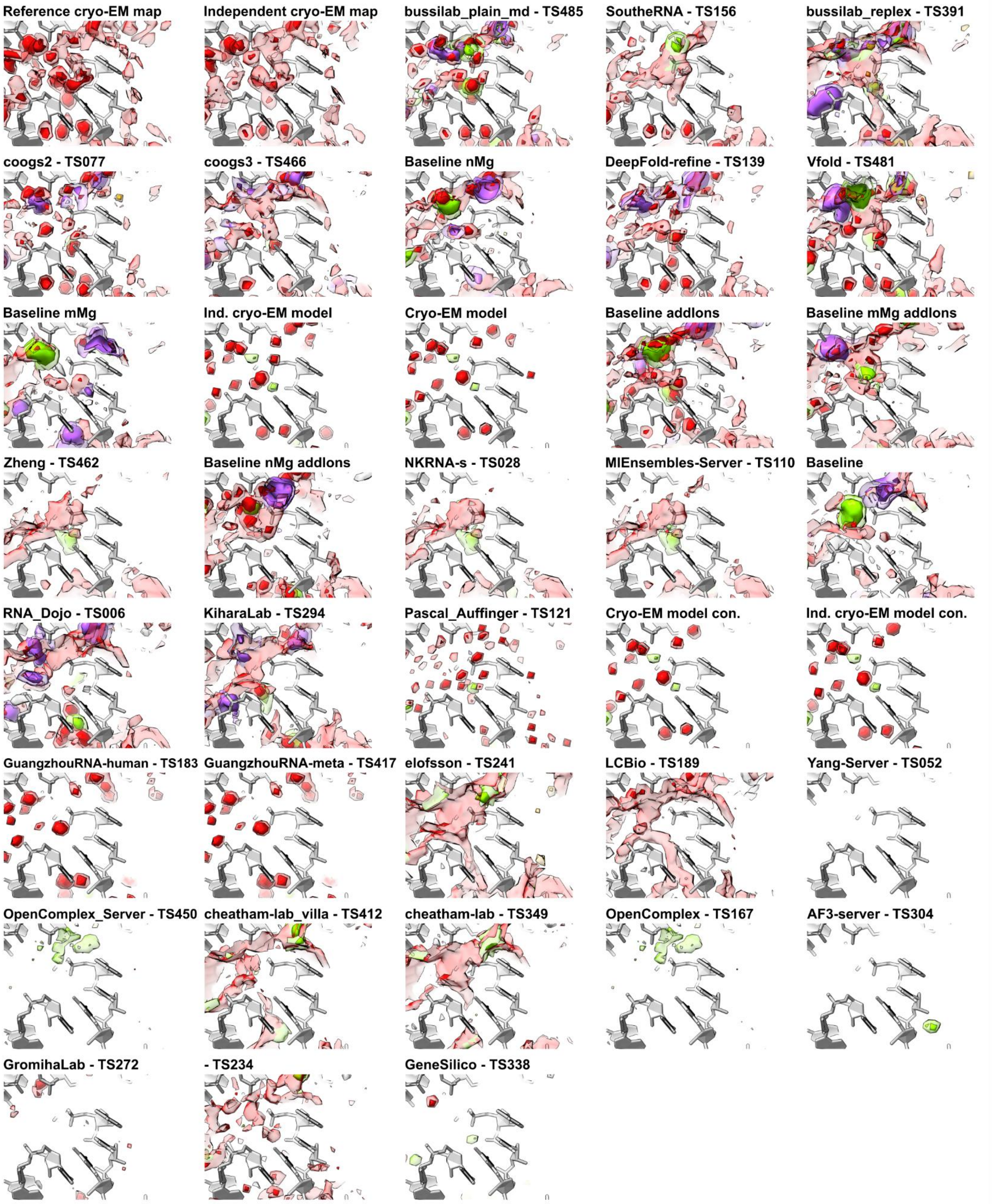


**Supplemental Figure 5: The local agreement between cryo-EM density and predicted water and ion density is shown for regions in Figure 7D.** The cryo-EM maps are displayed at 2 σ (light red) and 5 σ (dark red) – the molecular identity for any peaks is not assumed. For the predicted models, features with low density (light color) and high density (dark color) are shown for water (red), $Mg^{2+}$ ions (green), $Na^{+}$ ions (purple), and $Cl^{-}$ ions (yellow). The groups are ordered by ranking in **Figure 4**.

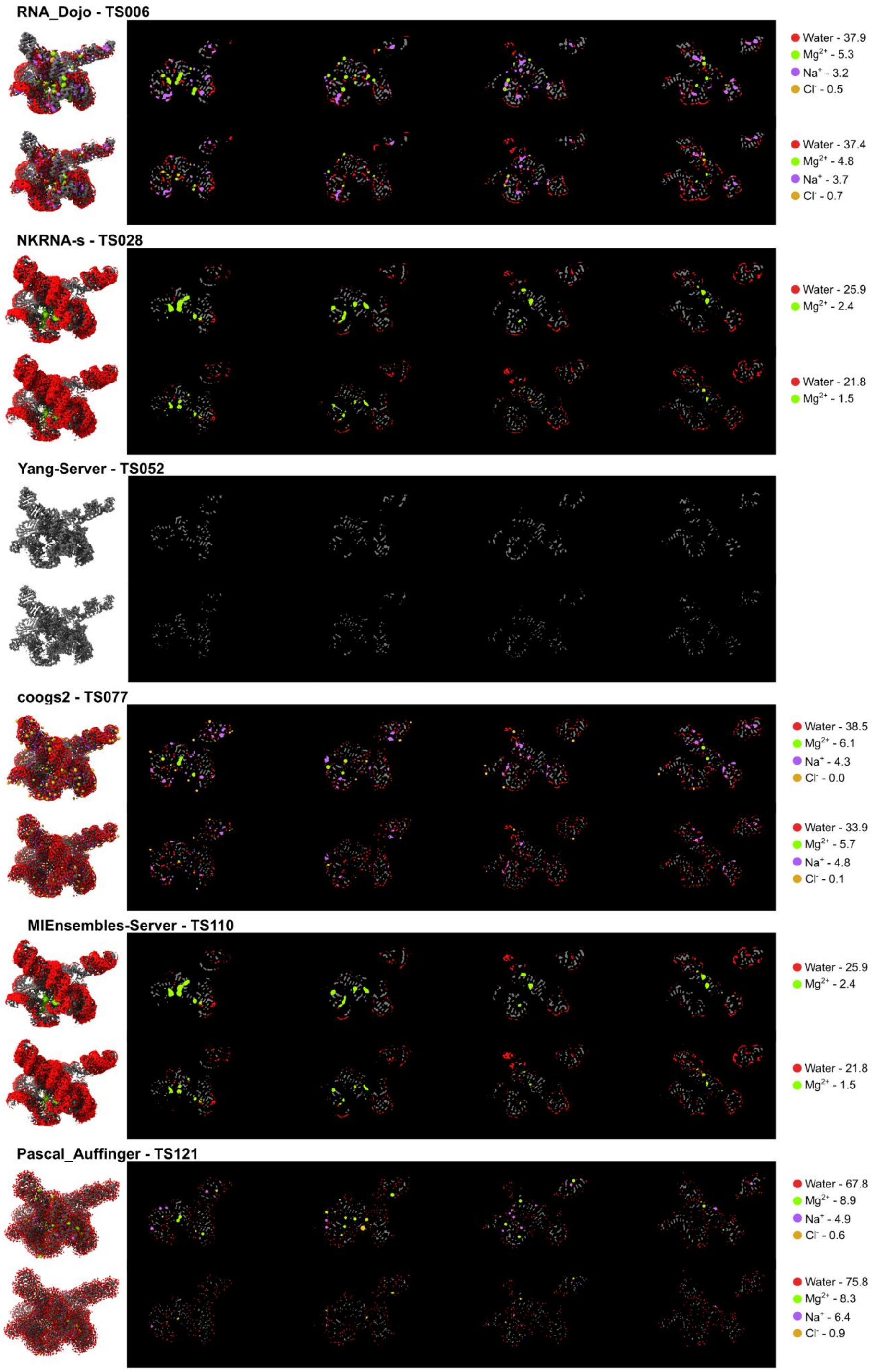


**Supplemental Figure 6: Densities derived from local alignment of atomic ensembles.** For each group the density of RNA (grey), water (red), $Mg^{2+}$ ions (green), $Na^{+}$ ions (light purple), $Cl^{-}$ ions (dark yellow), and $K^{+}$ ions (dark purple) are shown on the left as a 3D volume and in the center as a range of Z-slices. The top row for every group displays the $\text{Density}_{\text{scat}}$ and the bottom row the $\text{Density}_{\text{prob}}$. The densities for each molecule entity are displayed separately at 10 σ in order to effectively visualize the location of these densities even when low occupancy relative to other molecules. On the right of the image, the level of each displayed density is reported as a percentage of the RNA density level.

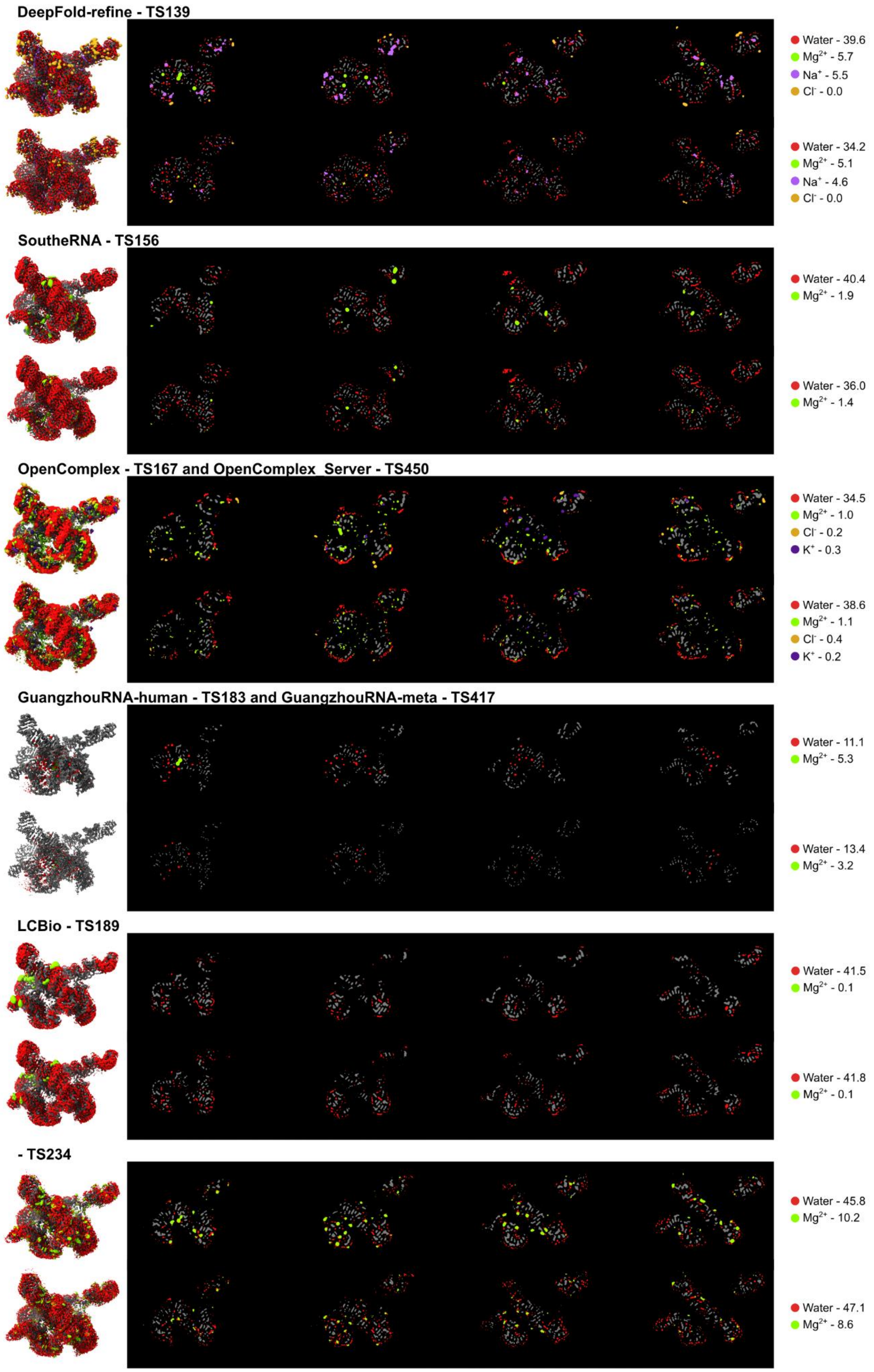


**Supplemental Figure 6 continued**

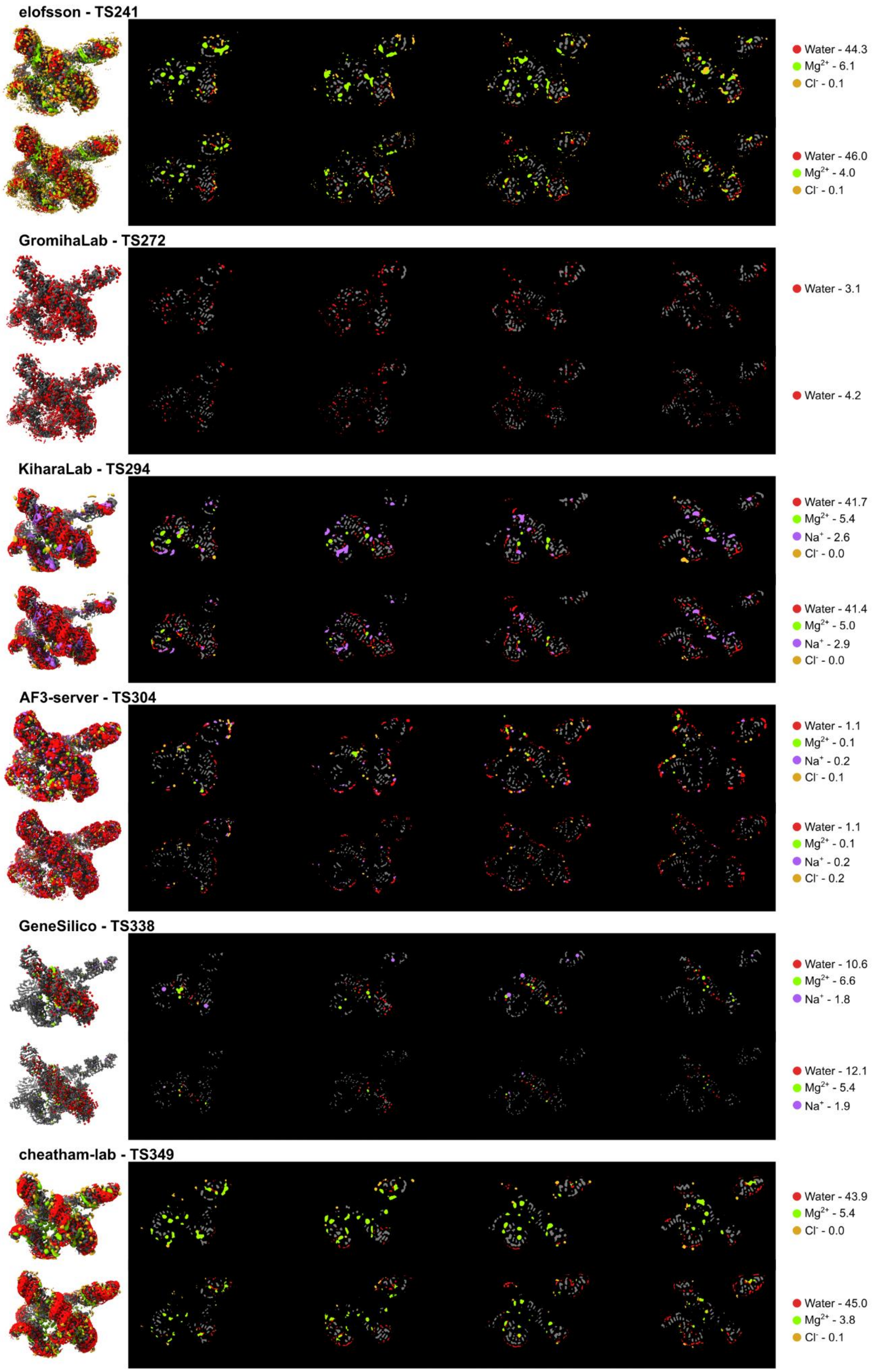


**Supplemental Figure 6 continued**

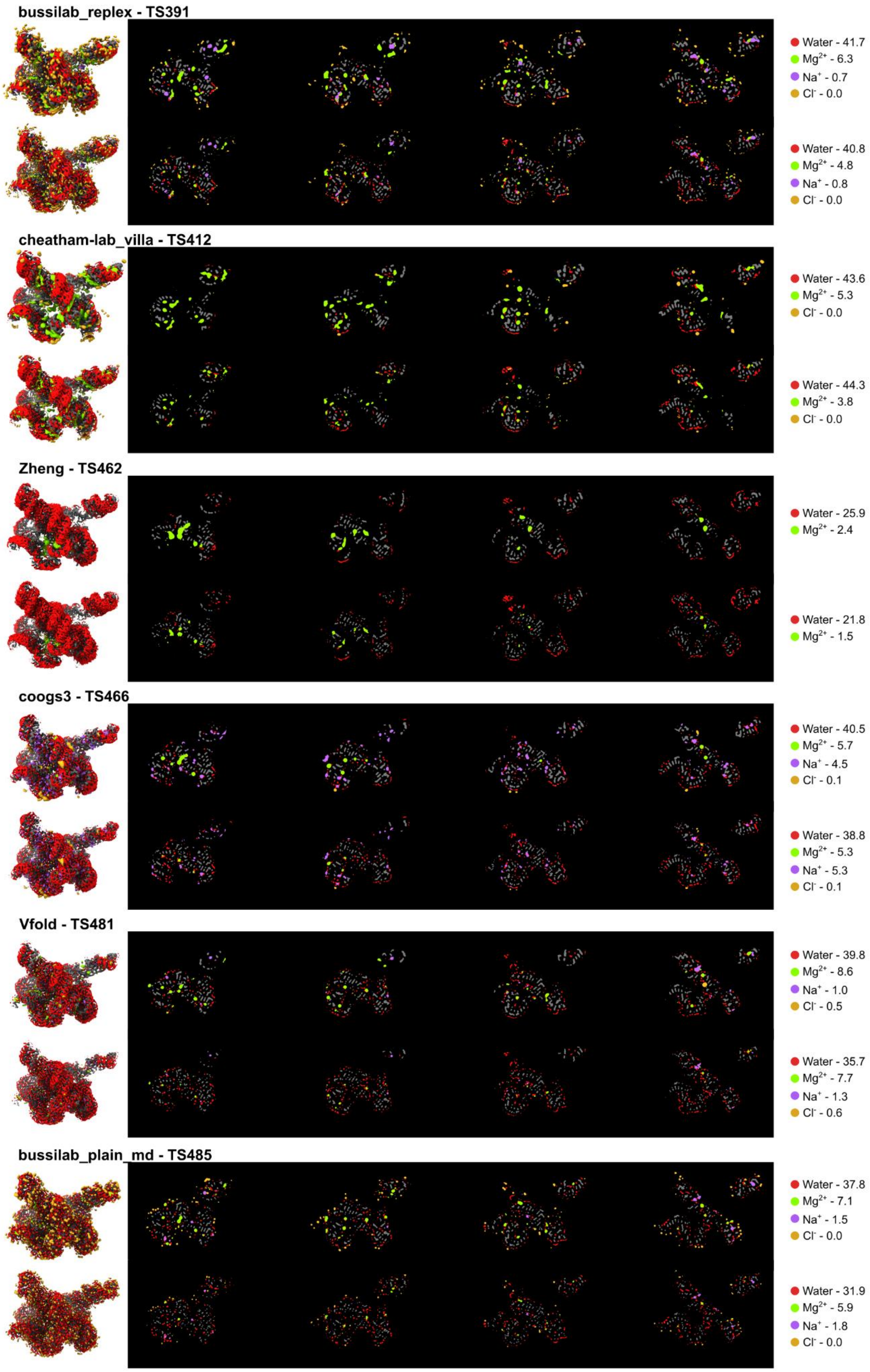


**Supplemental Figure 6 continued**

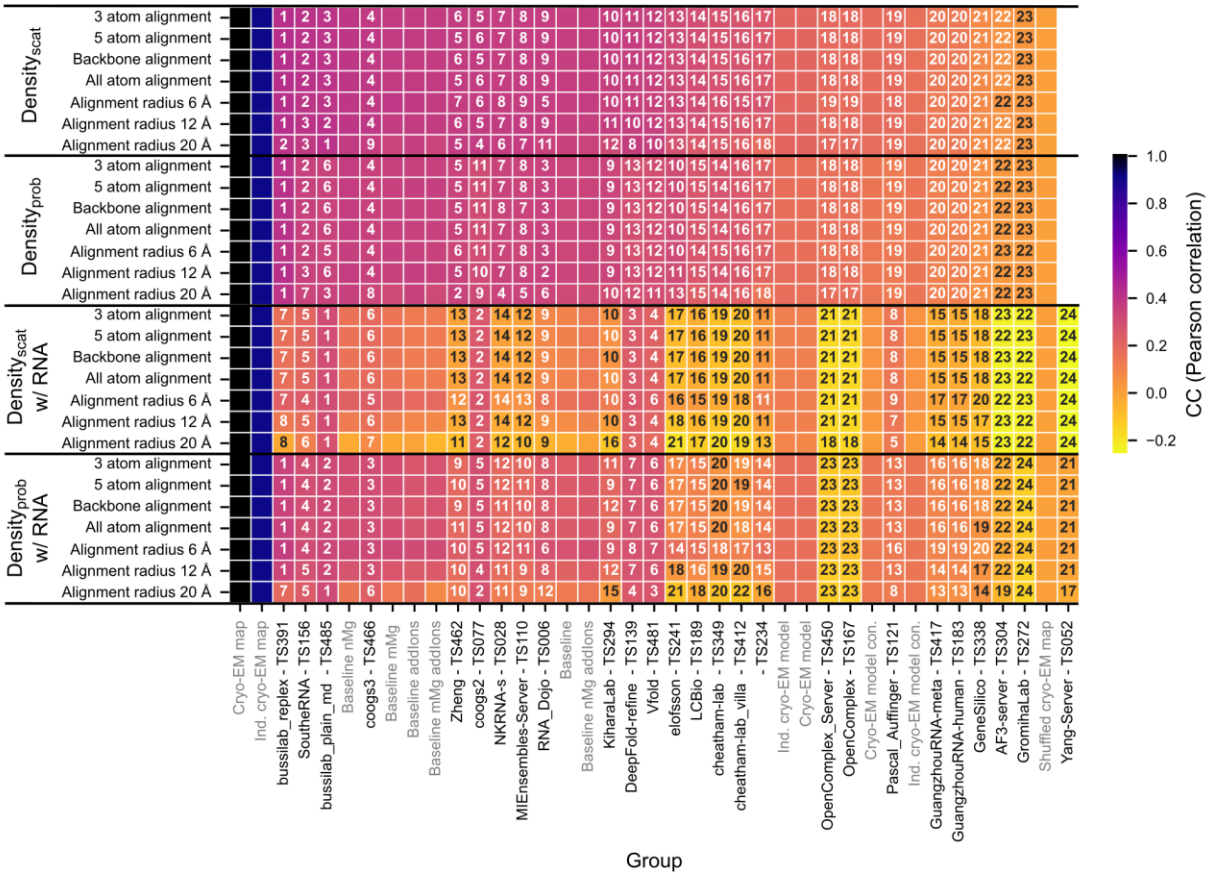

**Supplemental Figure 7: RNA local alignment method effects on density comparison.** The heatmap compares the 2.2 Å cryo-EM map (EMDB-42498) in the shell 1.8-3.2 Å away from the well-resolved RNA residues to the density derived from the predicted ensemble using various alignment methods. The plot is annotated with the rank the group would receive using that alignment method and colored by the CC (Pearson correlation). Groups are ordered by performance with the density$_{scat}$ for the alignment of all heavy atoms within 10 Å radius. A similar trend is observed for other metrics.

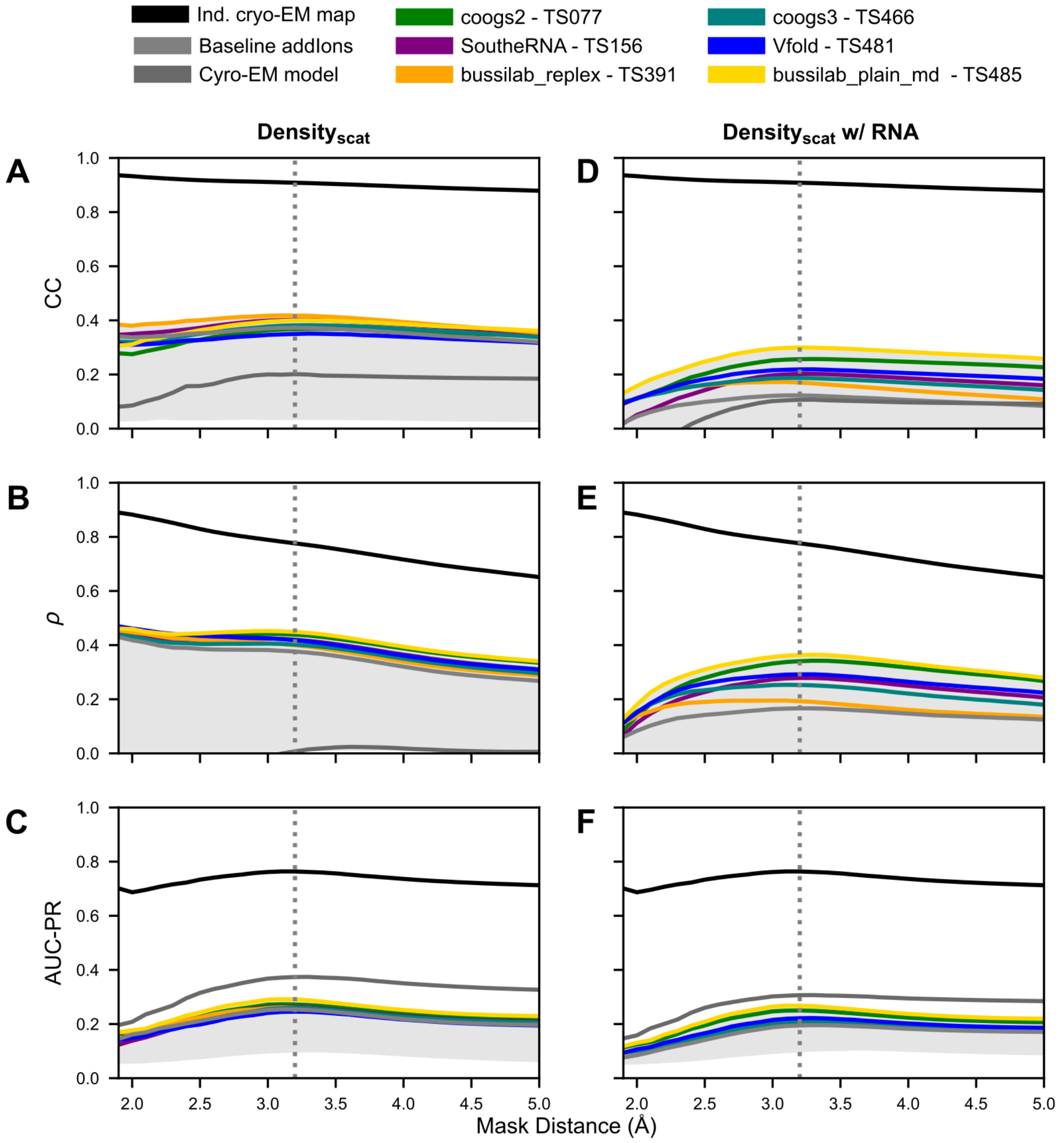


**Supplemental Figure 8: Prediction accuracy for the solvent shell.** The solvent shell originates 1.8 Å from the RNA and is extended to the mask distance. The shaded grey area represents the range of score for all CASP16 predictors.

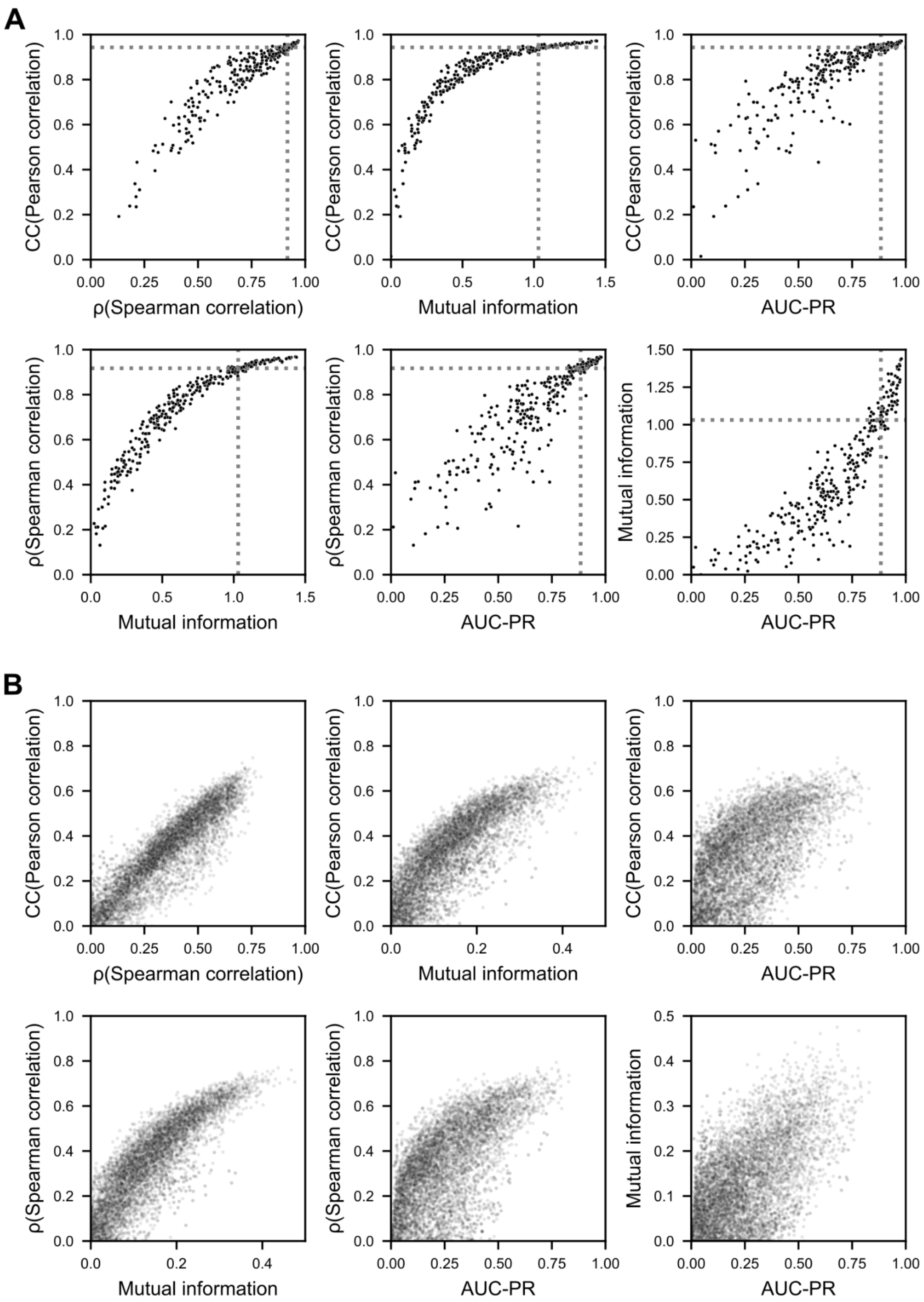


**Supplemental Figure 9: Correlation between metrics studied.** (**A**) For each nucleotide the local scores for the comparison between the independent cryo-EM map and the reference cryo-EM map are plotted. The grey dotted lines represent the boundaries used to define the nucleotide plotted in **Figure 6B**. (**B**) The local scores for every prediction group, for every nucleotide are plotted.

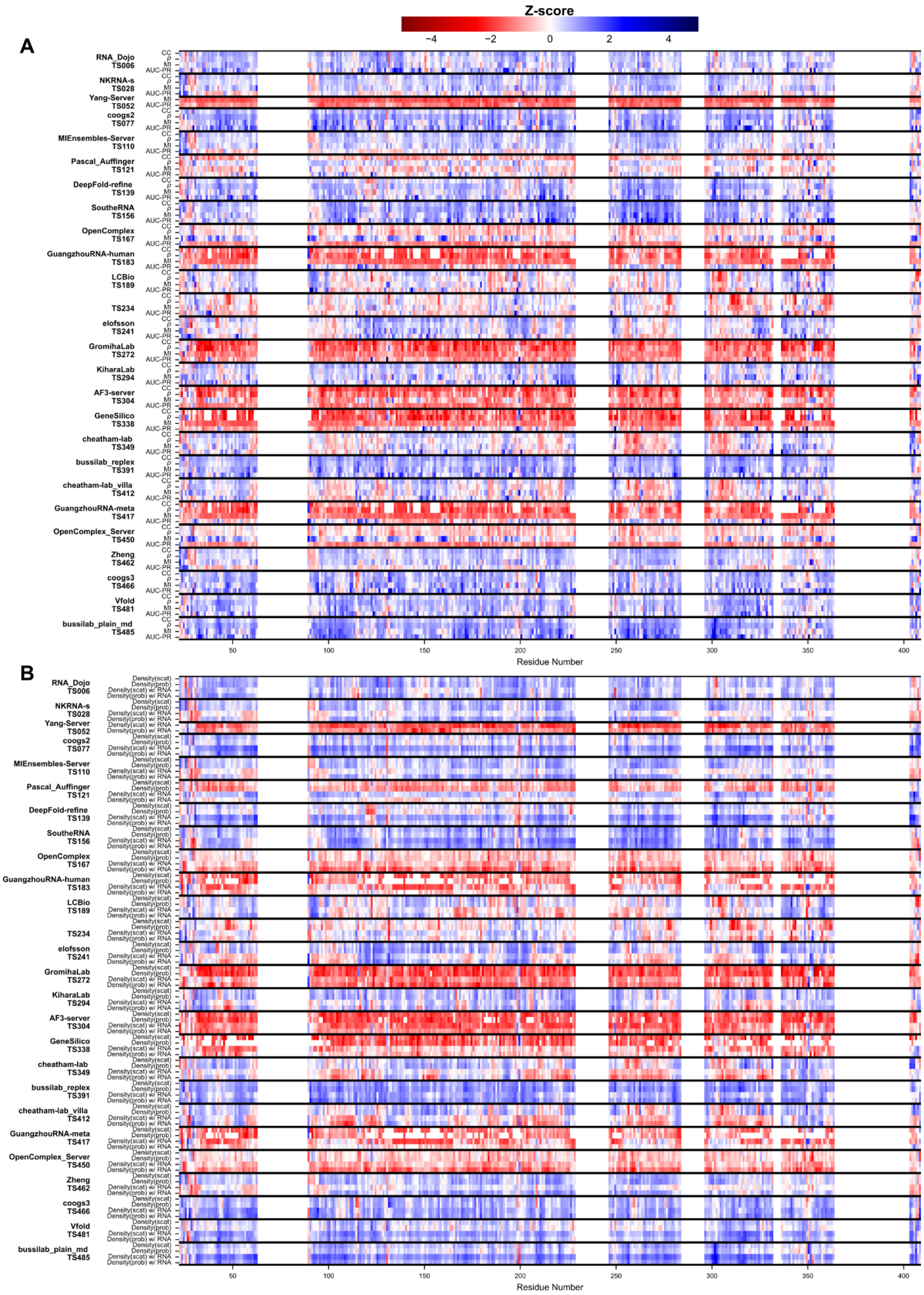


**Supplemental Figure 10: Comparison of per-residue scoring metrics.** The Z-score, standard deviation above the mean for all groups, was calculated for: (**A**) the four scoring metrics on Density$_{scat}$ predicted volumes (**B**) CC for the four density generation methods.

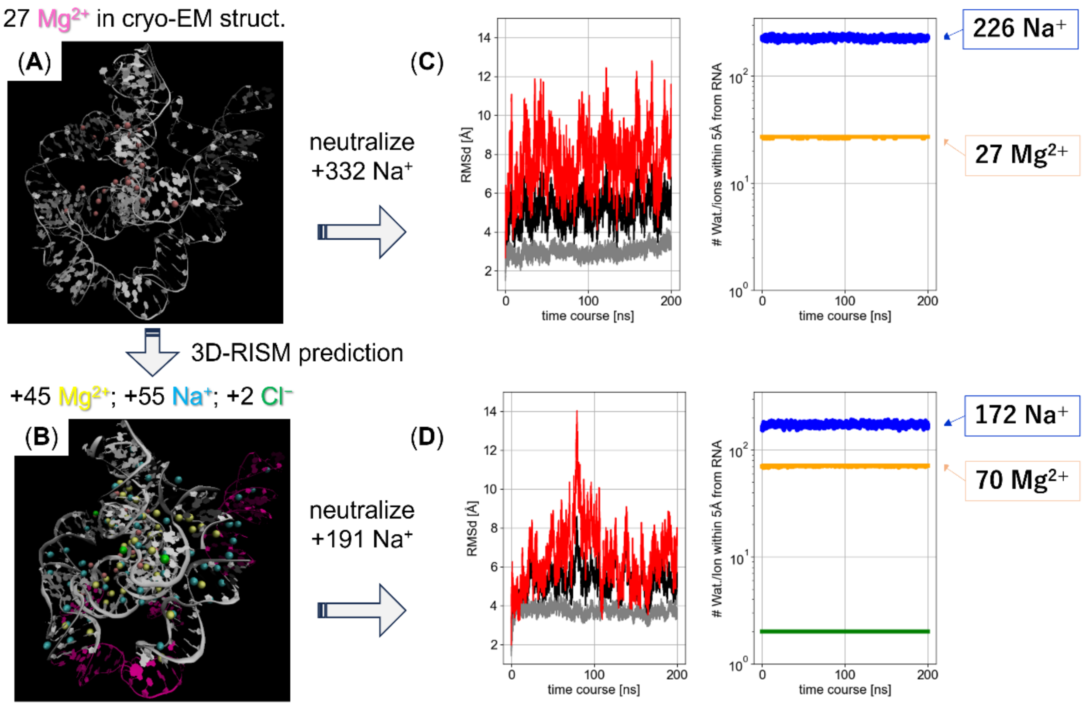


**Supplemental Figure 11: Analysis of RNA Dojo predictions.** (**A**) Cryo-EM structure. (**B**) Cryo-EM structure with additional ions predicted by 3D-RISM. (**C**) and (**D**) are MD trajectory analyses for RNA systems without and with 3D-RISM predicted ions. In panels (**A**) and (**B**), experimentally-derived and 3D-RISM-derived $Mg^{2+}$ ions are colored by pink and yellow, respectively. In panel (**B**), the flexible regions (see **Methods**) are colored by magenta. The left panels in (**C**) and (**D**) display the temporal profile of RMSD, where black, grey and red denote whole, core and fluctuating regions of RNA, respectively. The right panels in (**C**) and (**D**) are for ions coordinated inside a 5 Å solvation shell around the whole RNA, where blue, orange and green denote $Na^+$, $Mg^{2+}$ and $Cl^-$, respectively.

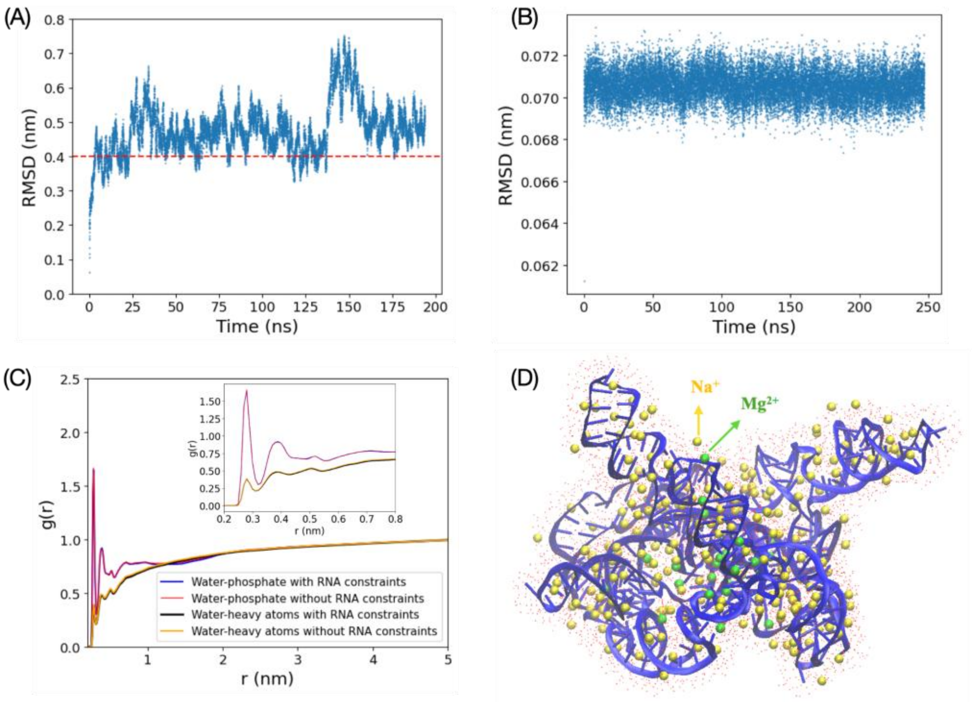


**Supplemental Figure 12: Analysis of Coogs2 and Coogs3 predictions.** (**A**) RMSD of RNA heavy atoms during the simulation without positional restraints (Coogs 3) in the production run. (**B**) RMSD of RNA heavy atoms during the simulation with positional restraints in the production run (Coogs 2). (**C**) Radial distribution function, g(r), of oxygen atoms of water relative to RNA phosphate groups and all RNA heavy atoms (i.e., non-hydrogen) for both simulations; the inset is a zoomed-in g(r) at low r for the visualization of the first few solvation shells. (**D**) A representative snapshot of RNA and its solvation shell within 5 Å, with red dots representing water molecules and green and yellow beads representing $Mg^{2+}$ and $Na^{+}$ ions, respectively.

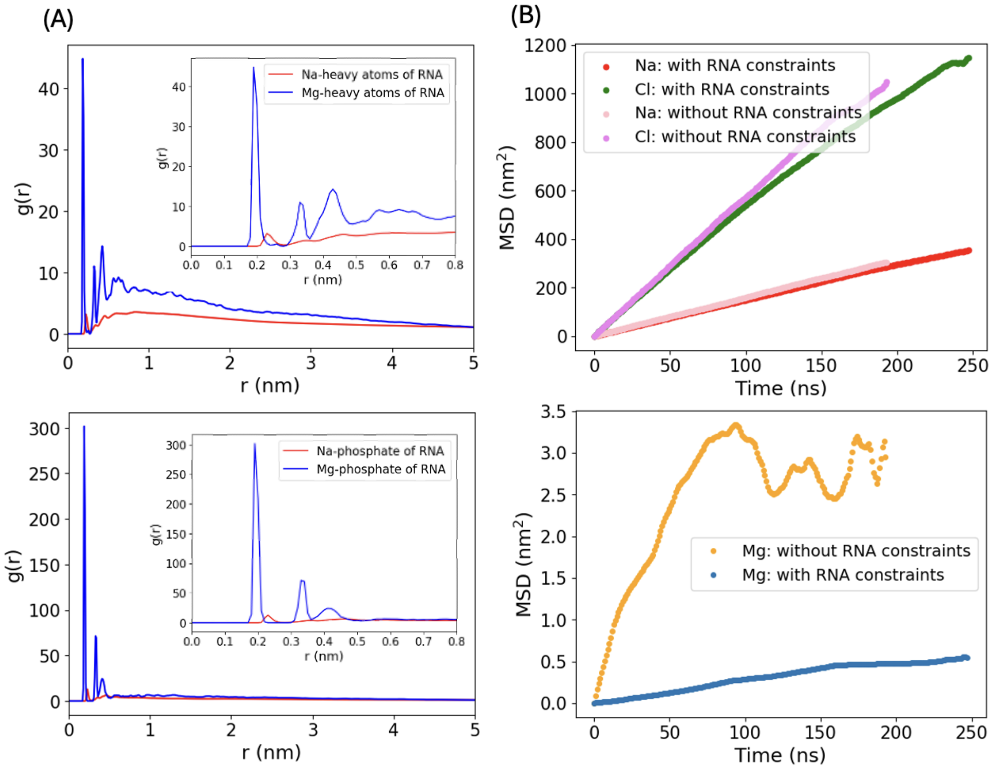


**Supplemental Figure 13: Analysis of Coogs2 and Coogs3 predictions.** (**A**) Radial distribution function, g(r), of $Mg^{2+}$ and $Na^{+}$ ions with respect to the heavy atoms (top) and phosphate atoms (bottom) of the RNA for simulation with constraints on RNA atoms (Coogs2); the inset is a zoomed-in g(r) at low r. (**B**) Mean square displacement (MSD) of ions during the simulations with and without positional restraints on RNA atoms (Coogs 3) for $Na^{+}$ and $Cl^{-}$ ions (top) and $Mg^{2+}$ ions (bottom).

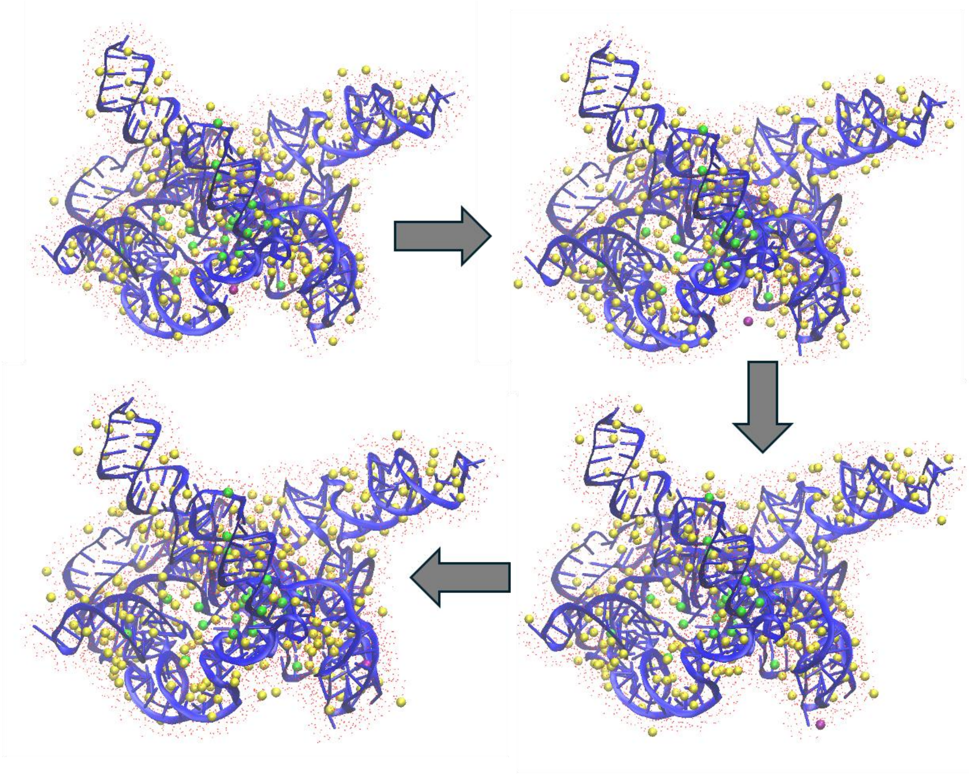

**Supplemental Figure 14: Analysis of Coogs2 predictions.** Snapshots showing the movement of the $Mg^{2+}$ ion (magenta color) from one RNA pocket (between nucleotides A21, U22, A150, and U146) in the native structure (7EZ0[1]) to another pocket (between nucleotides A102, C103, G173, G174, and A175). All other $Mg^{2+}$ ions are shown in green, and all $Na^{+}$ ions are shown in yellow.

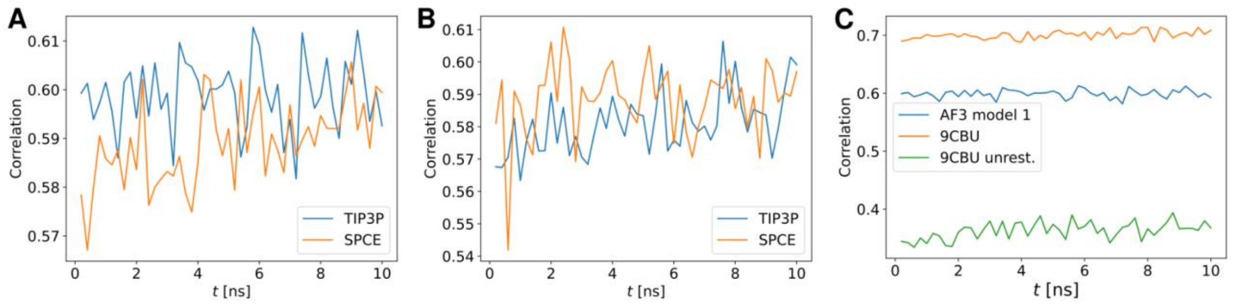


**Supplemental Figure 15: Analysis of SoutheRNA predictions.** Correlation between electron density EMD-42499 and values generated by Chimera, using different water models for (**A**) AF3 model 1 and (**B**) AF3 model 2. The comparison between AF3 model 1 and the simulation of reference structure 9CBU with and without restraints is shown in (**C**) using the TIP3P water model in all cases . Simulations had a length of 10 ns.

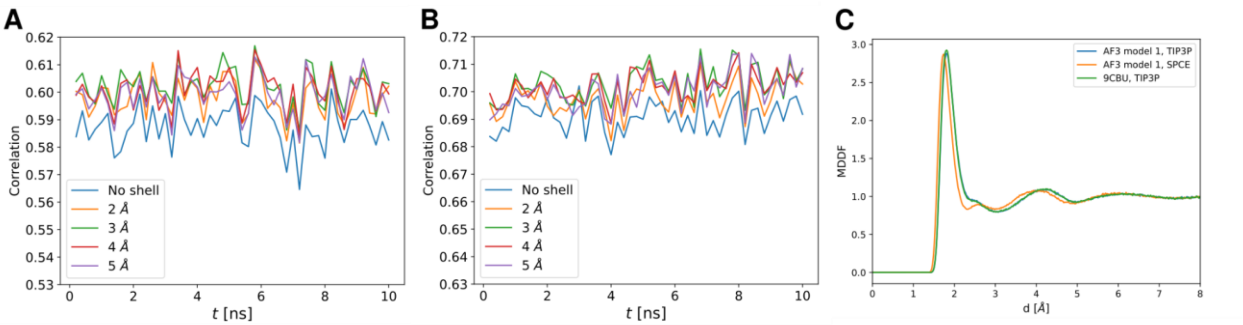


**Supplemental Figure 16: Analysis of SoutheRNA predictions.** Correlation between cryo-EM map EMD-42499 density values and values generated by Chimera for atomic models, as a function of time using a water shell of different thickness for (**A**) AF3 model 1 and (**B**) reference structure 9CBU. In (**C**) we can see the Minimum Distance Distribution Function (MDDF) for three different simulations of 10 ns.

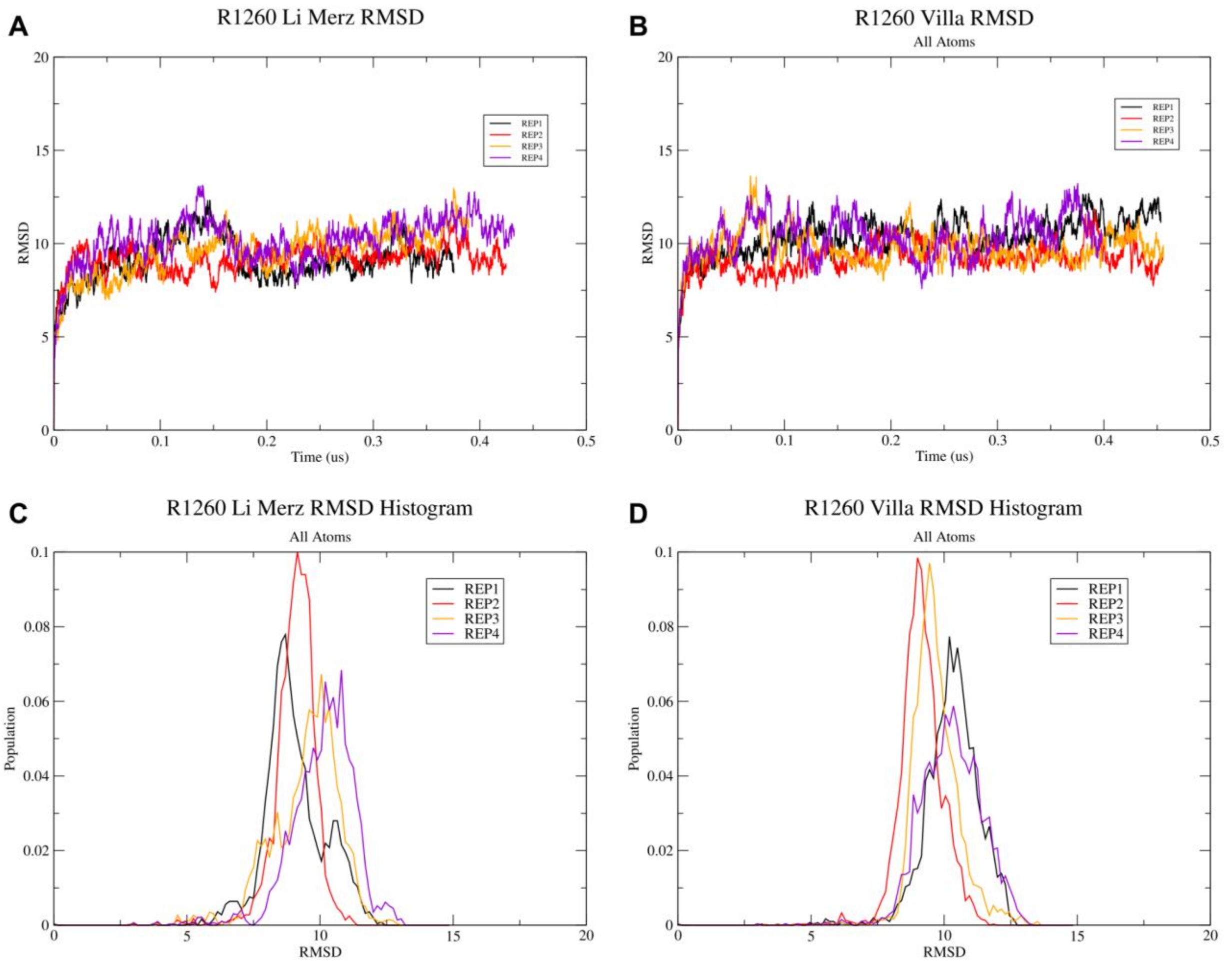


**Supplemental Figure 17: Analysis of Cheatham Lab predictions.** RMSD vs. Time plots (top) and RMSD histogram plots showcasing the population of structures compared to the first frame reference structure. RMSD calculations contained all non-hydrogen atoms and were calculated for each replica using every ten frames.

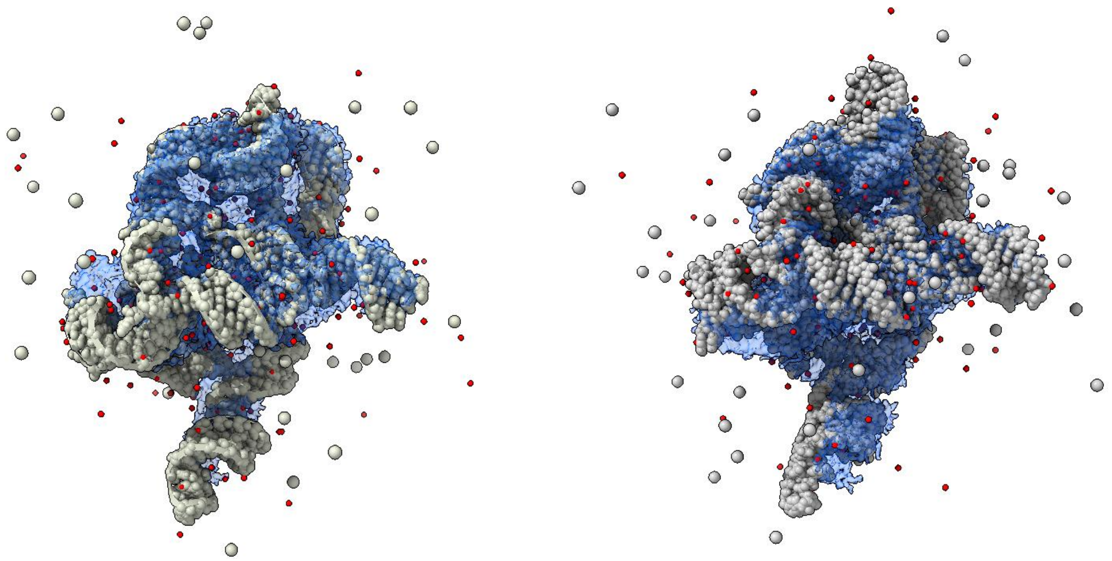

**Supplemental Figure 18: Analysis of Cheatham Lab predictions.** Superimposition of Li Merz (left) and Villa (right) representative structures (grey) with EMD-42498 map (blue). $Mg^{2+}$ ions are colored red, and $Cl^{-}$ ions in white.

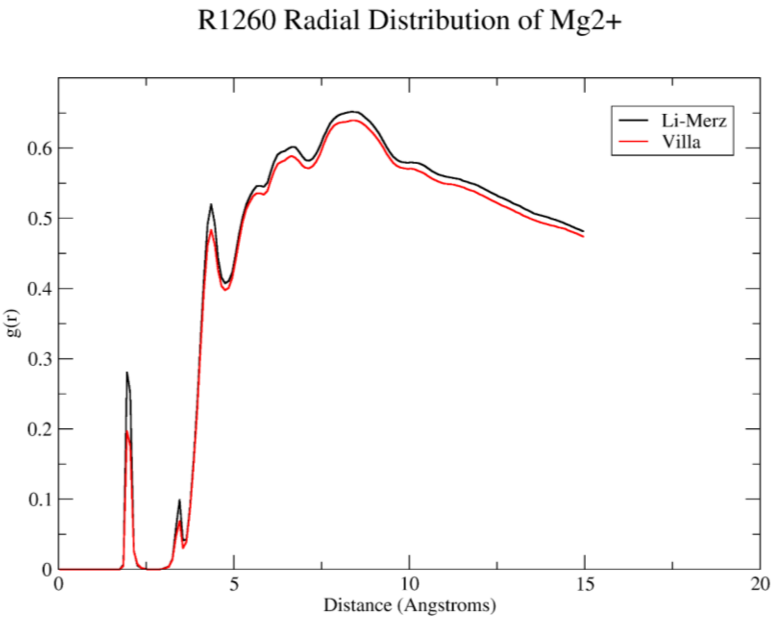


**Supplemental Figure 19: Analysis of Cheatham Lab predictions.** Graphed radial distribution function of distance between $Mg^{2+}$ ions and RNA. Li Merz divalent ion model shown in black and Villa divalent ion model shown in red.

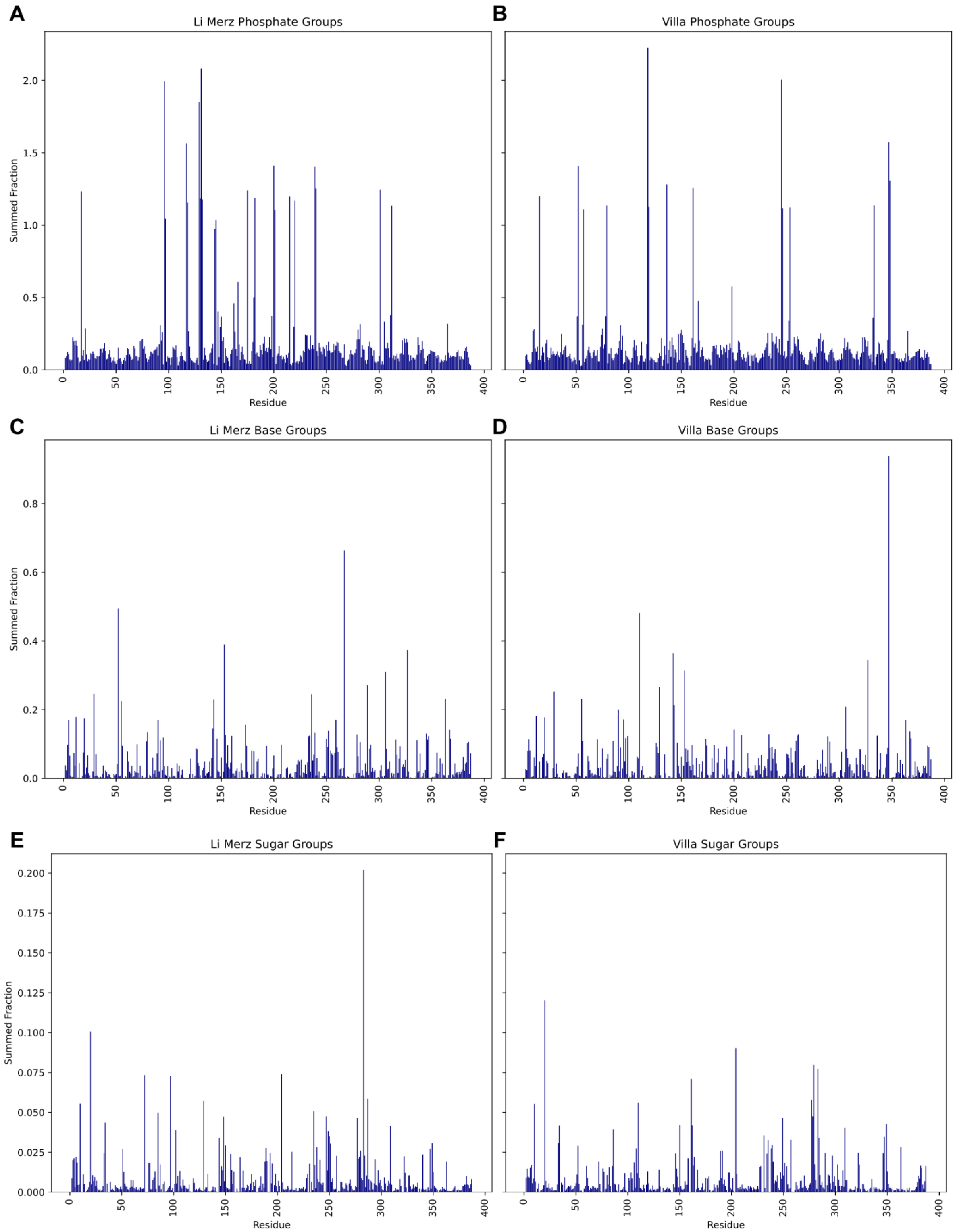


**Supplemental Figure 20: Analysis of Cheatham Lab predictions.** Summed fraction occupancies of binding grouped by (**A,B**) phosphate atom interactions (O3’, O5’, OP1, and OP2), (**C,D**) base atom interactions (N9, N7, O6, N1, N2, N3, N4, O2, N6, and O4), and (**E,F**) sugar atom interactions (O4’ and O2’) for (**A,C,E**) Li Merz and (**B,D,F**) Villa simulations.

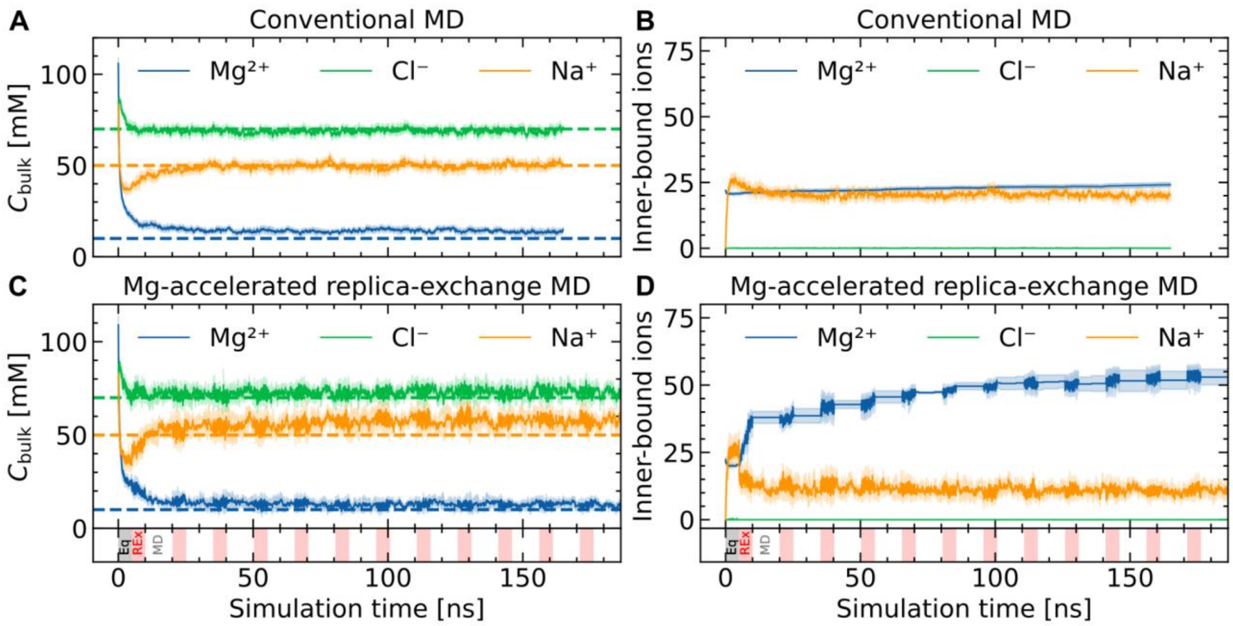


**Supplemental Figure 21: Analysis of BussiLab predictions.** Concentration of ions in the bulk (**A,C**) and number of ions bound to a phosphate non-bridging oxygen (**B,D**) as a function of simulation time, obtained using conventional MD (**A,B**) or enhanced sampling (**C,D**). The horizontal axis reports the simulated time per replica. Shaded regions in panels (**c**) and (**d**) indicate the replica-exchange segments. Dashed lines correspond to the experimental concentration.

**Supplemental Table 1: Molecular components of RNA systems for RNA Dojo prediction.**

| Molecule | Reference | 3D-RISM prediction* |
|---|---|---|
| $Mg^{2+}$ | 27 | 72 (45) |
| $Na^{+}$ | 332 | 246 (55) |
| $Cl^{-}$ | 0 | 2 (2) |
| Water | 54204 | 54204 |

* Number in parentheses denotes 3D-RISM derived ion's number.

**Supplemental Table 2: Analysis of GeneSilico predictions.** Prediction quality of the simulated (model1) and manual (model2) models in capturing magnesium ions and water atoms.

$Mg^{2+}$

| query | reference | TP (F1,RC) < 1.0 Å | TP (F1,RC) < 2.0 Å | TP (F1,RC) < 3.0 Å | TP (F1,RC) < 4.0 Å | TP (F1,RC) < 5.0 Å |
|---|---|---|---|---|---|---|
| model1 | 9CBX | 2 (0.05,0.04) | 4 (0.10,0.07) | 7 (0.18,0.13) | 8 (0.20,0.15) | 10 (0.25,0.19) |
| model1 | 9CBU | 2 (0.09,0.09) | 4 (0.17,0.18) | 5 (0.21,0.23) | 5 (0.21,0.23) | 5 (0.21,0.23) |
| model1 | model2 | 6 (0.12,0.08) | 14 (0.28,0.19) | 15 (0.3,0.2) | 19 (0.38,0.25) | 20 (0.4,0.27) |
| model2 | 9CBX | 7 (0.11,0.13) | 11 (0.17,0.20) | 13 (0.20,0.24) | 20 (0.31,0.37) | 23 (0.36,0.43) |
| model2 | 9CBU | 4 (0.08,0.18) | 7 (0.14,0.32) | 7 (0.14,0.32) | 9 (0.19,0.41) | 11 (0.23, 0.5) |

Water

| query | reference | TP (F1,RC) < 1.0 Å | TP (F1,RC) < 2.0 Å | TP (F1,RC) < 3.0 Å | TP (F1,RC) < 4.0 Å | TP (F1,RC) < 5.0 Å |
|---|---|---|---|---|---|---|
| model1 | 9CBX | 3 (0.02,0.01) | 5 (0.03,0.02) | 6 (0.04,0.02) | 10 (0.07,0.04) | 11 (0.07,0.04) |
| model1 | 9CBU | 3 (0.04,0.02) | 5 (0.07,0.04) | 6 (0.08,0.04) | 7 (0.09,0.05) | 8 (0.1,0.06) |
| model1 | model2 | 0 (0.0,0.0) | 3 (0.01,0.01) | 6 (0.02,0.01) | 7 (0.02,0.01) | 9 (0.03,0.02) |
| model2 | 9CBX | 9 (0.02,0.03) | 51 (0.12,0.18) | 77 (0.18,0.28) | 103 (0.25,0.37) | 118 (0.28,0.42) |
| model2 | 9CBU | 4 (0.01,0.03) | 25 (0.07,0.19) | 38 (0.11,0.28) | 54 (0.16,0.40) | 62 (0.18,0.46) |

TP – True Positives, F1 – F-score, RC – Recall.

**Supplemental Table 3: Analysis of BussiLab ion predictions.** Results of the replica-exchange simulation were averaged for 2 replicates × 5 unbiased replicas, skipping the initial equilibration and sampling round (20 ns). Results of the conventional MD were averaged for 16 replicates, skipping the initial 10 ns. Standard errors are smaller than the reported precision.

| Ion | Total ions | Target bulk concentration [mM] | Method | Number inner ions d(ion-OP) < 2.5 Å | Number outer ions 2.5 Å < d(ion-OP) < 5 Å | Number bulk ions d(ion-OP) > 20 Å | Bulk concentration [mM] |
|---|---|---|---|---|---|---|---|
| $Mg^{2+}$ | 178 | 10 | replex | 48.0 | 87.7 | 12.2 | 13.0 |
| | | | plainmd | 22.8 | 121.3 | 13.6 | 14.4 |
| $Cl^{-}$ | 91 | 70 | replex | 0.0 | 0.5 | 68.6 | 72.7 |
| | | | plainmd | 0.0 | 1.0 | 64.9 | 68.9 |
| $Na^{+}$ | 121 | 50 | replex | 11.3 | 14.8 | 53.5 | 56.8 |
| | | | plainmd | 20.4 | 20.0 | 46.6 | 49.4 |

**Supplemental Text 1: RNA Dojo TS006 - Ikuo Kurisaki, Junichi Iwakiri, Kengo Sato, and Michiaki Hamada**

In the challenge for CASP16 target R1260, the target RNA structure was previously resolved with 27 $Mg^{2+}$ molecules by cryo-EM (PDB 7EZ0[1]). This RNA has 386 negative charges from the phosphate groups in total. Recalling apparent violation of electrostatic neutrality, a substantial number of cations are probably *overlooked* through the tertiary structure modeling, which may be due to technical challenge to reliably identify ion species interacting with RNA[2,3]. We predicted experimentally invisible ions around RNA by using atomistic simulations.

MD simulations can theoretically examine spontaneous ion diffusion around RNA and subsequent ion-binding to RNA. However, simple use of this approach would not necessarily work well due to two concerns. First, ion-binding sites, which may be formed in an experimentally resolved RNA structure, may be deformed or missing in MD simulations because of thermal fluctuation. Such perturbation would interfere with the correct prediction of $Mg^{2+}$ binding to RNA. Second, it is not trivial to estimate how many ions should be added to a given RNA system. Since ions condense around RNA beyond ion's bulk concentration[4], the actual numbers of ions cannot be simply estimated by using information of the ion's concentration and system volume. Considering the above two concerns, we first predicted the ions bound to the experimentally-resolved RNA structure. We used three-dimensional Reference Interaction Site Model (3D-RISM)[5,6] to predict an initial model of experimentally-invisible but possibly-present ions around the RNA.

The 3D-RISM calculation was performed under 100 mM NaCl and 10 mM $MgCl_2$ (**Supplemental Figure 11B**). Kovalent-Hirata closure is employed to solve RISM equations. RISM-predicted ions are seemingly abundant ($Mg^{2+}$: 529, $Na^+$: 750, $Cl^-$: 39) and then were clustered by using Density Based Spatial Clustering of Applications with Noise (DBSCAN) algorithm[7], where $\varepsilon$ (inter-atomic distance [Å] as radius to search neighboring points) is set to 4 and minimum number of core points is set to 3 (these parameters are here chosen as test). The representative ions derived from DBSCAN calculations are retained for the following MD simulation. The final molecular components are summarized in **Supplemental Table 1**. This system is referred to as *3D-RISM prediction*, hereafter. Meanwhile, we also constructed a *reference system* with a minimum number of $Mg^{2+}$ by simply adding $Na^+$ molecules to the experimentally-resolved structure to electronically neutralize the system (**Supplemental Figure 11A**).

For each of the two systems, we performed 200-ns NPT MD simulations after temperature and density relaxation simulations and then 500 snapshot structures were extracted from the last 50-ns time domain sampling uniformly in a 100-ps interval. In total, the 1,000 snapshot structures are deposited as prediction candidates for the R1260 target of CASP16. Computational details are summarized below. To calculate the forces acting among atoms, AMBER force field 99 OL3 with backbone phosphate modification[8], SPCE water model[9], Aqvist's $Mg^{2+}$ parameter[10], and Joung/Cheatham ion parameters adjusted for the SPCE[11] water model were applied for RNA residues, water molecules, $Mg^{2+}$, and monovalent ions ($Na^+$ and $Cl^-$), respectively. The system temperature and pressure were regulated with a Berendsen thermostat with a 5 ps of coupling constant[12] and Monte Carlo barostat with volume change by each 100 steps, respectively. 3D-RISM computations were carried out with sander.MPI module of Amber22[13] on CPU machines (AMD E7763) of the Research Center for Computational Science. The MD trajectories were analyzed with the Cpptraj module of Amber22[13]. MD simulations were performed using the Amber 22 GPU-version PMEMD module based on SPFP algorithm[14] with NVIDIA GeForce RTX 3090.

**Supplemental Text 2: NKRNA-s TS028, MIEnsembles-Server TS110, Zheng Group TS462 - Wentao Ni, Qiqige Wuyun, Gang Hu, and Wei Zheng**

The Zheng Group, NKRNAs, and MIEnsembles-Server employed deep learning methods, including AlphaFold3 and DeepProtNA, to predict the structure of R1260. Additionally, GROMACS MD simulations were utilized to model the water shell and ion distribution around the RNA.

*RNA structure prediction*. The AlphaFold3 server[15] was employed to generate 1,000 models for R1260, with each model comprising the RNA structure alongside a variable number of $Mg^{2+}$ ions, ranging from 19 to 40. The distribution of $Mg^{2+}$ ions followed an approximately normal pattern, with a peak at 27 at the center, gradually decreasing in both directions. This pattern aligns with the experimental structure (PDB 7EZ0[1]), which contains 27 $Mg^{2+}$ ions. To ensure the reliability of the predictions, only models with a TM-score[16] greater than 0.8, relative to the experimental structure, were retained. For models where the RNA had a TM-score below 0.8, we replaced the RNA component with a high-quality structure predicted using our in-house nucleic acid structure prediction method, DeepProtNA, while maintaining the original $Mg^{2+}$ ions. DeepProtNA is an end-to-end deep learning algorithm developed specifically for the prediction of nucleic acid-related structures. It integrates pre-trained language model embeddings, multiple sequence alignment (MSA) information, predicted secondary structures, and structural templates to directly generate three-dimensional coordinates. ESM[17] and RNA-FM[18] were utilized to generate high-dimensional sequence embeddings for protein and RNA sequences, respectively. MSAs were constructed using a modified version of DeepMSA2[19] for proteins and rMSA[20] for nucleic acids. The core architecture of DeepProtNA is inspired by AlphaFold2[21], incorporating a similar attention-based mechanism to process multi-dimensional features. This network translates sequence information directly into spatial coordinates for each residue and nucleotide, ultimately generating the final structure.

*Water shell and ion creation*. MD simulations were performed using GROMACS[22] on the selected models to create a water shell. The system was solvated by adding water molecules, and $Na^{+}$ and $Cl^{-}$ ions were introduced to neutralize the system and maintain charge balance. The SPC/E water model was applied for solvation, and the AMBER03 force field was utilized. Energy minimization was then performed to relax the system using the steepest descent algorithm, with a maximum force threshold of 1000 kJ/mol/nm, running for 50,000 steps. The system was subsequently equilibrated in two phases under NVT and NPT conditions. Following equilibration, production MD simulations were carried out for 100 ns. Afterward, the trajectory was processed to remove periodic boundary conditions and align the system. Only the RNA, water molecules within 5 Å of the RNA structure, and $Mg^{2+}$ ions were selected. Finally, PDB files were generated from the trajectory, saving one model every 10 ns.

For the Zheng Group (#462), the complete process, including energy minimization, equilibration, and subsequent steps, as described above, was followed. In contrast, for NKRNA-s (#028) and MIEnsembles-Server (#110), only energy minimization was performed.

**Supplemental Text 3: Coogs2 TS077, Coogs3 TS446 - Ayush Gupta, Karim Malekzadeh, and Gül H. Zerze**

We submitted two predictions using conventional MD simulations, one with restrained RNA (TS077) and the other (TS446) without restraints. Further details of the simulations are given below.

The starting structure for the *Tetrahymena* ribozyme RNA was taken from the PDB (7EZ0[1]). This PDB contained 27 $Mg^{2+}$ ions around the RNA molecule. We maintained $Mg^{2+}$ ions at positions given in the PDB file and added the corresponding number of $Cl^{-}$ ions (54) to adjust the salt as $MgCl_2$. We modeled the RNA sequence using the nucleic acid force field DESRES[23] and combined it with the TIP4PD[24] water model. A single copy of the RNA molecule was solvated in a cuboidal box of volume 2060 $nm^3$. 386 $Na^+$ ions were added to maintain electroneutrality. The ions were modeled using CHARMM22 parameters[25]. After solvation and salt addition, we energy-minimized the system using the steepest descent algorithm. The simulation box was then equilibrated with a 100-ps NVT simulation (T = 300 K) followed by a 2-ns NPT simulation (T = 300 K, P = 1 bar). In both the equilibration steps, the positions of all the heavy atoms of RNA were restrained using a force constant of 500 kJ/(mol $nm^2$). We used GROMACS 2023.5[26] to perform all the production runs at 300 K and 1 bar, with a time step of 2 fs. Electrostatic interactions were calculated using the particle-mesh Ewald method[27] with a real space cutoff distance of 1 nm. A cutoff distance of 1 nm was also used for the van der Waals interactions. The final production run was done using two routes (submitted as two separate predictions): in the first submission (Group number: 077) the positions of all the heavy atoms of RNA were restrained using a force constant of 250 kJ/(mol $nm^2$), while in the second (Group number: 446), we didn't apply any restraints to the RNA atoms.

**Supplemental Text 4: SoutheRNA TS156 - Fabrizio Pucci and Simón Poblete**

Our procedure consisted of solvating RNA structures generated with AlphaFold3 (AF3)[15] with different water models and sampling their conformations using short MD simulations at constant temperature. The energy minimizations, solvation procedures, and simulations employed in the method were performed using the Gromacs2021.2 package[22].

Firstly, we generated 5 RNA models using the AlphaFold3 webserver starting from the sole sequence of the target. We selected the first and second AF3 models, which differed mainly in the conformation of the first 6 nucleotides, giving a total RMSD of 2.2 Å. The RMSD between our models and the reference structure (PDB 9CBU) was of 4 and 4.6 Å respectively. We also checked the absence of topological artifacts[28,29] in the selected models.

The systems were neutralized with Na atoms to be later solvated in SPC/E water with $MgCl_2$ at a concentration of 10 mM. The chosen force field was AMBER99 with parmbsc0 and ChiOL corrections[30–32], using the description of Joung and Cheatham[11] for the ion interactions.

The structures were submitted to an energy minimization procedure and then equilibrated by a short simulation with constant pressure of 1atm (using Berendsen barostat) and constant temperature of 300K (using a stochastic velocity rescale thermostat with a constant of 0.1ps) for 1 ns. Finally, a production run at constant temperature of 5 ns was generated using RMSD restraints with the predicted structure as a reference, with Plumed 2.9[33] as a harmonic moving restraint with a linearly increasing spring constant of $10^6$ kcal/mol·$nm^2$ every 200 ps, starting from 0. The final ensemble included 100 frames of each system.

**Supplemental Text 5: LCBio TS189 - Chandran Nithin, Smita P Pilla, and Sebastian Kmiecik**

The initial model of the target RNA was prepared based on a template from PDB (PDB 7EZ0[1]). The structure was energy minimized with 5000 steps of QRNAS[34], followed by MD simulations. MD simulations were performed using the Amber 22 package[13]. Initial structures were prepared using tleap, enclosed in a truncated octahedral box with a 10 Å buffer using tleap[35]. The system was neutralized with $Na^+$ ions. Since the samples were folded and frozen in a solution in 50 mM Na-HEPES and 10 mM $MgCl_2$, we added 50 mM NaCl and 10 mM $MgCl_2$ to the system. Solvation was performed using the TIP3P water model[9]. The χOL3 force field[32,36], specific for RNA, was applied. Energy minimization was conducted in two stages of 10,000 cycles each. In the first stage, positional restraints were applied to the backbone phosphorus (P) and oxygen atoms (OP1, OP2) of the RNA backbone with a restraint weight (restraint_wt) of 20.0 kcal/mol·$Å^2$. The second stage was performed without any restraints. Energy minimization and all subsequent simulation steps were facilitated using the CUDA-accelerated PMEMD[14,37,38]. The system was gradually heated from 100 K to 300 K over 500 ps (250,000 steps) with a time step of 2 fs. During heating, positional restraints with a weight of 20.0 kcal/mol·$Å^2$ were maintained on the RNA backbone atoms. The heating process was controlled using the Langevin thermostat (ntt=3)[39] with a collision frequency (gamma_ln) of 5.0 ps−1, and random seed (ig=-1) for stochastic dynamics. The temperature was ramped linearly from 100 K (tempi) to 300 K (temp0) during the simulation. Other key parameters included setting the nonbonded cutoff to 12 Å (cut=12.0), and enabling SHAKE constraints on bonds involving hydrogen (ntc=2, ntf=2). Following heating, the system underwent a four-phase equilibration process under NVT conditions. Initially, the system was equilibrated for 200 ps with positional restraints applied to the backbone atoms (O, OP1, OP2, P) using a restraint weight of 10.0 kcal/mol·$Å^2$. The temperature was maintained at 300 K using the Langevin thermostat (gamma_ln=5.0), with a 12 Å cutoff for non-bonded interactions and SHAKE constraints on hydrogen bonds (ntc=2, ntf=2). This was followed by an additional 200 ps equilibration with the restraint weight reduced to 5.0 kcal/mol·$Å^2$, and another 200 ps with the restraint weight further lowered to 1.0 kcal/mol·$Å^2$. Finally, a 2 ns equilibration was conducted without restraints, under the same non-bonded interaction cutoff and SHAKE constraints, ensuring the system's stability before proceeding to the production phase. The production phase of the simulation was conducted under constant pressure conditions using the NPT ensemble. A total of 100 ns of simulation time was performed with a 2 fs time step, resulting in 50 million steps (nstlim=50000000). The system was maintained at 298 K (temp0=298.0) using the Langevin thermostat (ntt=3) with a collision frequency (gamma_ln) of 1.0 ps-1. The pressure was set to 1.0 atm (pres0=1.0) with isotropic position scaling (ntp=1) and a pressure relaxation time (taup) of 2.0 ps. The Particle-Mesh Ewald (PME)[27] method was used for calculating electrostatic interactions with a 12 Å cutoff for non-bonded interactions (cut=12.0). SHAKE constraints were applied to bonds involving hydrogen (ntf=2, ntc=2). The simulation was set to restart from previous coordinates and velocities (irest=1) with random seed initialization (ig=-1). After the simulations 1000 models were selected from the trajectory of initial 10 ns. Only water molecules within 5 Å distance from at least one of the atoms from the RNA chain were retained.

**Supplemental Text 6: Elofsson TS241 and AF3-server TS304 - Arne Elofsson**

For the target R1260, we initiated our process by utilizing the AlphaFold3 server[15] to generate default predictions of the RNA structure. In total, we performed 100 predictions across 20 different seed numbers to ensure a comprehensive exploration of possible structural conformations. The primary aim of these AF3 server predictions was to establish a robust baseline for our analyses. To prepare the models for simulation, we solvated all 100 generated structures using a standard TIP3P water model, and ions were added randomly to maintain electrical neutrality and mimic physiological conditions. It is worth noting that we did not apply any additional refinements to these initial predictions.

In addition to the AlphaFold3 predictions, we also utilized models from Elofsson. For this, we commenced with the highest-ranked AlphaFold3 model. Following the same protocol, we solvated this model and incorporated ions using the default GROMACS setup[22]. After establishing the initial conditions, we allowed the system to undergo a relaxation phase to optimize the structure, which is crucial for achieving stable simulation conditions. Once relaxation was completed, the system was subjected to MD simulations for a total duration of 47.4 nanoseconds. During this simulation, we extracted 473 snapshots, capturing ten snapshots per nanosecond, which were subsequently submitted to the prediction center. This thorough approach ensures a detailed understanding of the dynamic behaviours present in the modelled system.

**Supplemental Text 7: GromihaLab TS272 - Ambuj Srivastava, K. Harini, Sowmya Ramaswamy Krishnan, and M. Michael Gromiha**

Our pipeline begins with an intense literature survey to gather experimental data on the target RNA. We have predicted the secondary and three-dimensional structures of the RNA utilizing various existing tools. Final models were selected with human expertise based on structural homologs and experimental evidence from the literature such as the presence of higher-order structures including pseudoknots. For the target R1260, we predicted the secondary structures using RNAFold[40], RedFold[41], and IPKnot++[42]. Further, these predictions were used to generate tertiary structures of the RNA with trRosettaRNA[43] along with predictions from the AlphaFold3 server[15]. The generated models were compared to the cryo-EM structure of a template of the RNA (PDB 7EZ0[1]) for the final selection. In comparison, predictions from AlphaFold3 showed the least deviation from the known structure (RMSD ~3.5 Å), which was selected for further simulations.

The predicted structure was simulated using RNA.OL3 force field for RNA molecules in the AMBER22 program[13]. The structure was solvated in a cubic box using the TIP3P water model with a box boundary of 15 Å from the surface of the RNA molecule. The number of $Na^+$ and $Cl^-$ ions was calculated using the SLTCAP server[44] to maintain a salt concentration of 150 µM in the simulation box. The Langevin thermostat was used to gradually increase the temperature of the system from 0 K to 300 K with a step size of 2 fs for 100,000 steps. The system was further subjected to constant volume and pressure for 500,000 and 100,000 steps, respectively, each with a 2 fs timestep. Brenden barostat was used to maintain constant pressure, and the SHAKE algorithm was used to constrain hydrogen-involving bonds. Final minimization was performed for 10 ns with a step size of 2 fs, and trajectories were saved for every 5000 steps. Furthermore, the minimized trajectories were aligned and clustered into 10 groups using k-means clustering. We obtained an ensemble of folded RNA ribozyme structures within a water solvent shell. The structures showed variations mainly in the P6 and P9 domains of the RNA[1].

**Supplemental Text 8: KiharaLab TS294 - Jacob Verburgt, Yuki Kagaya, Tsukasa Nakamura, Anika Jain, Genki Terashi, Pranav Punuru, Emilia Tugolukova, Joon Hong Park, Anouka Saha, David Huang, and Daisuke Kihara**

We used a MD simulation approach to model the solvent shell in CASP16 target R1260. To provide our simulation with good initial ion placement for persistent ions, we fetched and superimposed several deposited models of the structure from the PDB. The specific PDB IDs were 7EZ0, 7YCI, 7YCH, 7YCG, 7YC8, 7YGD, 7YGC, 7YGB, 7YGA, 7YG9, 7YG8, 7XD7, 7XD4, 7XD3, 7XD6, 7XD5, 7XSN, 7XSM, 7XSL, 7XSK, 6WLS, 7UVT, 7EZ2, 8I7N, 8HD7, 8HD6, 7R6M, 7R6N, and 7R6L[1,45–51]. From these PDB structures, places where magnesium ions consistently exist across the majority of structures were noted and extracted. In total, we moved forward with 27 Mg positions. Using 7EZ0, we prepared a Desmond MD simulation[52] through the Schrodinger Maestro interface (Schrödinger Release 2024-3), ensuring that all the identified magnesium positions were still present after preparation. We had attempted to use an AlphaFold3[15] structure of the target but had complications in aligning the target with the predetermined placement of the ions. The system was prepared with explicit SPC water, and a 0.15 M sodium chloride buffer with neutralizing ions. We used sodium chloride instead of magnesium chloride as the system builder in Maestro 2024.3 does not support $Mg^{2+}$ as an ion. No restraints were placed on the RNA, and the system was subjected to a 10 ns simulation at 300 K. A 10 ns simulation was used due to time and resource constraints. We extracted separate PDB files for each of the resultant 1000 frames and fed them individually to a script to extract a 5 Å solvent shell. We further processed these 1000 solvent shell PDB files to match with standard PDB naming conventions and submitted the ensemble as our R1260 submission.

**Supplemental Text 9: GeneSilico TS338 - Masoud Amiri Farsani, Eugene Baulin, and Janusz M. Bujnicki**

For the R1260 target, our team submitted two models. The first model was generated by our in-house tool SimRNA-Sol (an extension of SimRNA), and the second model was built manually via template modeling.

SimRNA-Sol was run using the 7EZ0[1] structure, with frozen RNA coordinates. The simulation included 10 $Mg^{2+}$ ions, 10 $Na^{+}$ ions, and 10 water molecules. The simulation was run with default parameters, namely eight independent runs of a Monte Carlo Replica Exchange of 16,000,000 steps, 10 replicas, covering a temperature range from 1.35 to 0.9. The 1% best energy decoys were selected (8,000 structures) and subjected, for each solvent moiety, to HDBSCAN clustering, with a minimum cluster size set to 1% of the selected decoys (80 structures). From each cluster, the best-scored representative structure was selected for consideration. In the case of overlapping sites, the choice of the preferred solvent moiety at a given site was based on the inspection of the local environment.

The manual model was prepared using the ions and water molecules from the closest template structures determined experimentally. Overall we used eight templates of the P4-P6 domain (1GID[53] chains A and B, 1HR2[54] chains A and B, 1L8V[55] chains A and B, 2R8S[56], and 6D8O[57]) and 13 templates of the full ribozyme (7EZ0, 7R6L, 7XD5, 7XD6, 7YCI, 7YG8, 7YG9, 7YGA, 7YGB, 7YGC, 8I7N, 8TJV, 8TJX[1,45,46,48,49,58]). Each template was first superimposed onto 7EZ0 using ARTEMIS[59], followed by manual local adjustments performed in ChimeraX[60]. Then, all the templates were merged into a single model that included RNA from 7EZ0 and ions and water molecules from all the aligned templates. The merged model included 348 $Mg^{2+}$ ions, 762 water molecules, and no $Na^{+}$ ions. The P4-P6 domain templates contributed all 762 water molecules and 90 of 348 $Mg^{2+}$ ions. Subsequently, redundant ions were removed manually by visually identifying clusters of close atoms and removing all but one solvent moiety for each cluster. The final manually edited model included 75 $Mg^{2+}$ ions and 553 water molecules.

**Supplemental Text 10: Cheatham-lab TS349, Cheatham-lab_villa T412 - Olivia Fisher and Thomas Cheatham**

RNA structure is notoriously sensitive to its environment- namely, the presence of solvents, ions, and other biomolecules in the system. Water and ions play an especially important role in RNA folding, dynamics, and structure. Computational methods such as MD simulations can be an asset in elucidating these intricate RNA system dynamics as they facilitate the observation of conformational sampling over time and reveal inter/intramolecular interactions. MD simulation of these interactions at an atomistic level has proved difficult without preexisting experimental data. RNA structure has proven to be highly sensitive to its ionic environment, especially in the presence of $Mg^{2+}$, requiring ions for many roles, most notably for stabilizing folding and mediating ligand binding[61,62]. Recent experimental studies have discovered the reliance on $Mg^{2+}$ for riboswitch activity based on how ion presence shifts conformation populations towards a predominately folded state and increases the compactness of the structure. However, a major obstacle in understanding RNA dynamics using MD is current forcefield limitations in simulating $Mg^{2+}$ affinity with RNA, as the ion can get trapped in the wrong place and take longer than the realistic simulation length to become unstuck, effectively freezing the system in an anomalous conformation[63,64]. Inappropriate sampling caused by the chelation effect can lead to a sampling of conformational space that is biologically inaccurate and can be misleading in data interpretation of ion and water placement prediction.

Past work has shown that the selection of the ion parameters plays a critical role in representing accurate dynamics and structural changes of RNA[64]. We selected two commonly used divalent ion models to represent $Mg^{2+}$ in our CASP16 simulations, the Li Merz and the Villa ion parameters[65,66]. These models were selected because they have previously shown low levels of direct magnesium chelation that often happen during MD simulation, causing structural alterations and trapping the RNA for the length of the trajectory[64].

*Methods*. The experimental cryo-EM structure (PDB 7EZ0[67]) was used for the initial *Tetrahymena* Ribozyme RNA conformation. The hydrogen atoms and initial $Mg^{2+}$ ions were removed from the file before parametrizing the system. The RNA was described with the Amber OL3 RNA force field parameters[36] and solvated with a truncated octahedron 10.0 Å box using the OPC water model[68]. The system was neutralized using 194 $Mg^{2+}$ ions and 2 $Cl^-$ ions and solvated with 82,494 water molecules. To generate the 10 mM $MgCl_2$ experimental conditions, an additional 18 $Mg^{2+}$ and 35 $Cl^-$ ions were added to the system based on the initial volume. Two different parameter sets were used to model the $MgCl_2$ atoms: the Li Merz divalent ion parameters and the Villa divalent ion parameters. The van der Waal parameters were modified in the resulting topology files according to the Leonard-Jones backbone parameters, which increased the radii of the OP1, OP2, O5', and O3' oxygen atoms by 0.0884 Å[8,69]. Four replicas were run for each system to ensure reproducibility.

Each system was minimized with 1000 steps steepest descent minimization with strong restraints on heavy atoms followed by NTV MD at 300 K and SHAKE on hydrogens for 15 ps. Steepest descent minimization was done again first with relaxed restraints on heavy atoms, then minimal restraints, and then no restraints. Lastly, NTP MD was done with SHAKE, first with low restraints and then with minimal restraints on heavy atoms for 5 ps, then with minimal backbone restraints for 10 ps, and finally with no heavy atom restraints for 10 ps. After minimization, hydrogen mass repartitioning was done for both systems, which increased non-solvent hydrogen masses to 3.02 Da by redistributing weight from adjacent heavy atoms, allowing for an increased time step in production (2 fs to 4 fs)[70].

MD production was run with constant pressure and volume, with the temperature set to 300 K. The Villa systems ran for a combined total of 2.12 μs, and the Li Merz systems ran for 2.2 μs. Following production, trajectories from each replica were combined, and then the trajectories from each divalent ion model were clustered according to the K-means algorithm using CPPTRAJ[71]. The clusters with over 1000 frames were selected (6 clusters from Li Merz and 4 from Villa) to represent the most populated conformational states. For CASP submission, 50 frames were extracted evenly across each cluster and saved as PDBs, the hydrogens were removed to save space, and all 500 PDBs were submitted.

**Supplemental Text 11: bussilab_replex TS391, bussilab_plain_md TS485 - Elisa Posani, Olivier Languin Cattoen, and Giovanni Bussi**

We submitted two predictions, one using conventional MD simulations and the other with enhanced sampling to accelerate the rearrangement of $Mg^{2+}$ coordination shells. Simulations were done using GROMACS2 2022.3[22] and PLUMED3 2.8.1[72].

*Topology and force field.* The structure from PDB 7EZ0[1] was sanitized with PDBFixer. All 27 $Mg^{2+}$ ions present in the original structure were kept. We used the TIP3P[9] water model and the AMBER force field for RNA[30,32,36], microMg parameters for $Mg^{2+}$[73], and compatible NaCl parameters[74].

*Buffer preparation.* The structure was solvated in a rectangular box (~17×12×13 nm3) and neutralized with $Na^+$ and $Mg^{2+}$ in a 3-to-1 ratio (including the already present 27 $Mg^{2+}$) to match the ion competition observed in experiments[75]. Further ions were added in the bulk to reach experimental buffer conditions (10 mM MgCl2, and 50 mM Na-HEPES treated as NaCl). All extra ions (124 $Mg^{2+}$, 121 $Na^+$, 91 $Cl^-$) were added with gmx genion, with independent random seeds for each replica mentioned hereafter.

*Equilibration.* After energy minimization, the system was equilibrated for 5 ns in the NPT ensemble (1 bar[76], 300 K[77]), using positional restraints on the RNA heavy atoms and the 27 PDB $Mg^{2+}$ ions.

*RMSD restraint in production runs.* Restraints were applied to prevent RNA rotation and to limit its conformational dynamics. Two different settings were used for the two submissions. In the conventional MD simulation, we used GROMACS positional restraints to all RNA heavy atoms and the 27 PDB $Mg^{2+}$ ions, with a stiffness of 1000 kJ $mol^{-1}$ $nm^{-2}$. In the replica exchange simulation, the RMSD of a subset of RNA atoms (C1′, C2, P) to its native coordinates was computed with simple translational alignment, and a harmonic restraint centered at 0 nm was applied, using a harmonic constant of 200 kJ $mol^{-1}$ $nm^{-2}$. This value was empirically selected to obtain an average RMSD around 4 Å. The harmonic constants are not directly comparable because they are applied to different sets of atoms. In addition, the restraint on RMSD acts on the average deviation, so that it would correspond to GROMACS positional restraints with an even lower harmonic constant. Importantly, the 27 $Mg^{2+}$ ions present in the reference structure were not restrained in the enhanced sampling simulation.

*Conventional MD submission (group 485).* 16 replicates of the system were prepared independently and run for 165 ns each, for a total of 2640 ns. For each replicate, 62 equally spaced frames were extracted with a pace of 2.5 ns, skipping the initial 10 ns. This resulted in a total of 992 frames submitted for assessment.

*Accelerated Mg-binding dynamics.* Bias potentials were applied to each Mg ion as a function of its distance from the closest phosphate non-bridging oxygen and its coordination number to water oxygens[78]. The bias functional form was parameterized on a simple system composed of a diuridine molecule and a single magnesium ion in water, as explained in PLUMED tutorial 24.017[79]. Parameters were chosen by hand to compromise between the following goals: minimize barrier height on the observed reaction pathway; maximize overlap with the original ensemble; avoid the generation of new, spurious metastable states; maintain the overall binding affinity.

*Replica-exchange phase*. Ions' sampling was enhanced by coupling the bias-accelerated approach to a Hamiltonian replica-exchange setup[80,81], simulating a ladder of 16 replicas with various bias amplitudes. The first 5 replicas were kept bias-free, while the following 11 got increasing scaling factors for the bias potential. Spacings between the non-zero scaling factors were optimized to enforce homogeneous acceptance rates.

*Relaxation phase*. Hamiltonian replica-exchange phases were interspersed with conventional MD for the first 5 (bias-free) replicas. Our method thus alternates costly sampling with accelerated $Mg^{2+}$-binding kinetics, and cheaper sampling with conventional MD.

*Replica exchange submission (group 391)*. A total of 12 sampling rounds were simulated, each comprising 5 ns of replica exchange followed by 10 ns of conventional MD. Two independent replicates were simulated, totaling 16 + 16 = 32 replicas with different initial coordinates for bulk ions. The total accumulated simulation time was $2\times12\times(11\times5+5\times15)$ ns = 3120 ns. 10 equally spaced frames were extracted from each unbiased replica from the replica-exchange phases after discarding the first two rounds. This resulted in a total of $2\times10\times5\times10=1000$ frames submitted for assessment.

*Comparison of the two approaches.* The two submissions were done using identical force fields and equivalent initial conditions. The only differences are: (a) the replica exchange simulation samples the $Mg^{2+}$-RNA binding/unbinding events faster and (b) the conventional MD simulation used a stiffer restraint on RNA and an additional restraint on the already known $Mg^{2+}$ ions.

**Supplemental Text 12: Vfold TS481 - Yuanzhe Zhou, Sicheng Zhang, Jun Li, and Shi-Jie Chen**

The VFOLD group employed physics- and machine learning-informed MD simulations to model water and ion distributions around the target RNA. The computation involves three steps.

First, as the initial condition, we configure $Mg^{2+}$ ions according to ion positions in the PDB template structures. Since the target RNA, the *Tetrahymena* ribozyme, folds to conformations similar to PDB 7EZ0[1], we adopted the 27 $Mg^{2+}$ ions observed in this PDB structure. We then searched the PDB database for other templates that contain metal ions. The aligned templates show nearly identical global folds, with only the 5' and 3' ends (i.e., P1, P10 helix, internal guide sequence (IGS), and IGS extension) exhibiting large conformational flexibility. The alignment of 13 experimentally solved *Tetrahymena* ribozyme structures (PDB codes: 7EZ2[1], 7R6L[45], 7R6M[45], 7R6N[45], 7XD3[46], 7XD4[46], 7XD5[46], 7XD6[46], 7XD7[46], 7XSN[47], 7YGD[48], 8HD6[46], 8I7N[46] along with PDB 7EZ0 suggested three additional $Mg^{2+}$ binding sites. In total, 30 $Mg^{2+}$ ions were retrieved from the PDB structures.

Next, we use our in-house software tools, the physics-based Monte Carlo tightly bound ion (MCTBI) model[82–84] and the machine learning-based MgNet[85], to predict additional $Mg^{2+}$ ions. MCTBI is a statistical mechanics-based model that accounts for ion correlation and fluctuation effects in nucleic acids. The model predicts metal ion binding fractions, the most probable bound ion distribution, the electrostatic free energy including the free energy components. These results provide mechanistic insights into the role of metal ions in RNA structure formation and folding stability. Additionally, MgNet is a recently developed deep learning method that treats RNA-ion complexes as images, predicting $Mg^{2+}$ binding sites that are tightly coordinated with RNA atoms. MgNet is capable of identifying critical atoms for inner- and outer-sphere coordination and uncovering new ion binding motifs, making it a valuable complement to X-ray crystallography for precise metal ion site identification. We placed $Mg^{2+}$ ions to 43 MCTBI-predicted binding sites and 17 MgNet-predicted sites. In total, 60 additional $Mg^{2+}$ ions were produced in the second step.

Finally, we performed AMBER all-atom MD simulations to predict the solvent shell of the target RNA. The target RNA structure and $Mg^{2+}$ ions were solvated in a truncated octahedron box with a 15 Å buffer. To match the number of excess ions (i.e., the ions in excess of bulk buffer ions) predicted by MCTBI, 73 $Mg^{2+}$ and 30 $Na^+$ ions were added, along with 15 $Mg^{2+}$ and 73 $Na^+$ buffer ions based on the experimental conditions (i.e., 50 mM $Na^+$ and 10 mM $Mg^{2+}$). Additionally, 73 $Cl^-$ ions were also added to neutralize the system. The all-atom MD simulation was performed with the leaprc.RNA.OL3 force field for RNA[32,36], leaprc.water.OPC3 for water[86], and frcmod.ionslm_126_opc3 for ions (Li/Merz ion parameters for -1 to +4 in OPC3 water, 12-6 normal usage set[87–90]). The system underwent three consecutive 5000-step minimization, where movement was progressively allowed for hydrogen, water and ions, and parts of the RNA. Subsequently, the system was heated from 100 K to 298.15 K over 100 ps, with only hydrogen and water free to move. This was followed by a 200 ps simulation in the NPT ensemble. After the equilibration, the system was simulated in the NPT ensemble at a temperature of 298.15 K, with a simulation time step of 2 fs. Distance restraints were applied to the following nucleotides to maintain the canonical base pairs: 52, 54-60, 108, 110-112, 169-172, 209-213, 219-223, 263-265, 271-273, 311, 323-334, 346, 348-354, 358-360, 369-371, 375-381. Two ~550 ns simulations were performed with the same initial structures and different random seeds, and 1,000 snapshots were evenly extracted from the two simulation trajectories.

**Supplemental Text 13: Additional analysis by RNA Dojo TS006 - Ikuo Kurisaki, Junichi Iwakiri, Kengo Sato, and Michiaki Hamada**

The RMSD values for the core regions (sequential numbering: 1-41, 68-207, 225-262, 275-310, 315-342, 382-386; biological numbering: 22-62, 89-228, 246-283, 296-331, 336-363, 403-407) are ~4 Å (grey lines in left panels in **Supplemental Figure 11C-D**). Since the magnitude of RMSD values are similar to the CASP organizer-informed criterion, we use these trajectories to examine the number of $Mg^{2+}$ ions within the 5 Å solvation shell of the RNA. In the *3D-RISM-derived system*, not only experimentally-resolved 27 $Mg^{2+}$ molecules but also almost all of 3D-RISM-derived $Mg^{2+}$ molecules are found within the solvation shell (right panels in **Supplemental Figure 11C-D**). Our 3D RISM calculations accurately predict $Mg^{2+}$ molecules, which stably interact with the RNA but have been overlooked in the cryo-EM structures.

As for the influence of these ions on RNA, the 3D RISM-predicted ions may stabilize the flexible region of the RNA. This speculation comes from the following two observations (*see* red lines in left panels in **Supplemental Figure 11C-D**). First, RMSD values of the flexible regions are suppressed in the *3D-RISM-derived system* compared with the *reference system*. Second, RNA in the *3D-RISM-derived system* appears to be structurally robust. Actually, the RMSD value transiently increases around the time point of 80 ns, while this change is recovered by 100 ns in the MD trajectory. We can find a part of the flexible region that binds to $Na^{+}$ molecules, which are predicted by the 3D-RISM calculation (*see* **Supplemental Figure 11B**). The above analyses suggest that MD simulations combined with RNA-ion binding prediction with 3D-RISM calculations have technical potential to identify experimentally invisible RNA-bound ions and then provide further structural and dynamic insights into RNA molecules in aqueous solution.

**Supplemental Text 14: Additional analysis by Coogs2 TS077, Coogs3 TS446 - Ayush Gupta, Karim Malekzadeh, and Gül H. Zerze**

For both production runs, we computed the RMSD of the heavy atoms in the RNA structure (**Supplemental Figure 12A-B**). In the restrained simulation, all the sampled frames had an RMSD value of less than 0.4 nm from the native structure (7EZ0[1]). In the unrestrained simulation, frames with an RMSD above 0.4 nm were excluded for submission (Group number: TS446) as they are deemed unsuitable for predicting the surrounding solvent shell; however, we kept all the frames for the structure and transport properties below since they were useful for the analysis.

_Water structure_: The radial distribution functions, g(r), from both simulations (**Supplemental Figure 12C**) exhibit identical profiles, indicating that the spatial organization of water molecules around RNA remains unchanged regardless of whether positional restraints are applied. This suggests that restraining the heavy atoms of RNA does not significantly alter the hydration shell (while RMSD stays less than 0.8 nm), implying that the local water structure around phosphate groups and heavy atoms is primarily governed by the direct RNA-water interactions rather than RNA flexibility.

_Ion structure:_ To determine the arrangement of ions in the solvation shell, we calculated the g(r) of $Mg^{2+}$ and $Na^{+}$ ions with respect to the heavy atoms and phosphate atoms of the RNA (**Supplemental Figure 13A**). The earlier onset of the first peak of the g(r) for $Mg^{2+}$ compared to $Na^{+}$ ions shows that $Mg^{2+}$ ions are more closely arranged around the RNA than $Na^{+}$ ions. Our trajectories also showed that $Mg^{2+}$ ions remained bound to the RNA, while $Na^{+}$ ions are distributed more evenly, including in the bulk. $Mg^{2+}$ ions remained in the same pockets as in their initial condition for the unrestrained simulation, whereas one $Mg^{2+}$ ion (and only one $Mg^{2+}$ ion) switched its bound pocket in the unrestrained simulation (**Supplemental Figure 14** also see below for further discussion of this).

We also calculated the mean-square displacement (MSD) of ions (**Supplemental Figure 13B**). The MSD plots for $Mg^{2+}$ show that their displacement was higher in the simulation with unrestrained RNA structure, which is due to the close presence of $Mg^{2+}$ ions around the RNA (although $Mg^{2+}$ ions were not restrained). In the simulation with restraining the RNA atoms though, we observed a significant movement of one of the $Mg^{2+}$ ions (and only one of the $Mg^{2+}$ ions) from their original pocket (RNA nucleotides A21, U22, A150, and U146) in the native structure (7EZ0) to a different pocket with RNA nucleotides A102, C103, G173, G174, and A175 (**Supplemental Figure 14**). This newly found binding pocket of the $Mg^{2+}$ ion is not present in the experimental structure 9CBX.

We also found that $Na^{+}$ and $Cl^{-}$ ions were displaced to a much larger extent compared to $Mg^{2+}$, which is expected as $Na^{+}$ and $Cl^{-}$ ions occupy the bulk (in addition to the proximity of RNA). The MSD plots also show an order of magnitude difference between the displacement of $Na^{+}$ and $Cl^{-}$ ions in both production runs.

**Supplemental Text 15: Additional analysis by SoutheRNA TS156 - Fabrizio Pucci and Simón Poblete**

From our results, we observed that imposing the structural restraints was needed to avoid the distortion of the molecule's shape along the simulation. An unrestrained simulation of the reference structure 9CBU in TIP3P water reached an RMSD of 27 Å from its initial conformation in less than 5 ns of simulation. We also compared the correlation between the density generated by our ensemble structures and the cryo-EM map EMD-42499_2.2A_sharpened structure using Chimera[91] after optimal alignment with the reference structure 9CBU. The correlation for each frame of the ensemble, shown in **Supplemental Figure 15A-B** does not show a substantial difference for both RNA structures using SPC/E or TIP3P water models. However, the reference structure 9CBU with and without restraints shows relevant differences compared to the AF3 models as plotted in **Supplemental Figure 15C**. The thickness of the water layer considered around the RNA molecule also was found to increase marginally, but systematically, the correlation between our generated ensemble and the cryo-EM map as shown in **Supplemental Figure 16A-B**. Finally, the structure of the water around the RNA molecule was observed by plotting the Minimum Distance Distribution Function (MDDF) using the package ComplexMixtures.jl[92]. The plot of **Supplemental Figure 16C** shows no difference between the water distribution around the first AF3 model and 9CBU when using the TIP3P water model, but it differs appreciably when the SPC/E model is employed.

**Supplemental Text 16: Additional analysis by GromihaLab TS272 - Ambuj Srivastava, K. Harini, Sowmya Ramaswamy Krishnan, and M. Michael Gromiha**

We compared these predicted structures to the released cryo-EM derived models (PDB: 9CBW, 9CBU) using the RNA-align web server[93]. Interestingly, our predictions showed a RMSD in the range of 4.77-5.79 Å. Except for the P6 domain of the ribozyme (residues 204-227), all the other domains had relatively low deviation (2.28-4.45 Å) from the experimental structure. Similar to the crystal structure, the P6 domain protrudes into the solution without interacting with the other domains[94]. However, the simulations could not reproduce the water networks observed around the P6a domain in the cryo-EM model. With a longer simulation timescale, the diffusion of surface water molecules to these interior domains can be captured with high accuracy.

Based on the prediction experiments conducted in CASP16, we observed significant progress in protein structure prediction in recent years. However, RNA structure prediction still faces challenges, particularly on more complex higher-order structures or with larger RNA molecules. This opens the field with huge opportunities to improve RNA secondary/tertiary structure prediction algorithms. Although few tools are available in the literature, many web servers did not provide results in a stipulated time frame. Moreover, it is difficult to replicate the known homologous structures during the predictions. The development of the AlphaFold3 web server has addressed some of these issues. However, the accuracy of predictions still tends to be lower when it comes to novel folds or structures containing pseudoknots[95]. Additionally, there is a need for robust and user-friendly protein-nucleic acid complex prediction models.

**Supplemental Text 17: Additional analysis by GeneSilico TS338 - Masoud Amiri Farsani, Eugene Baulin, and Janusz M. Bujnicki**

We compared the two models to each other and to the reference models (automatic 9CBX and consensus 9CBU). For that, in each pairwise comparison, we greedily matched the atoms of interest ($Mg^{2+}$ ions or water molecules) with a threshold of 1.0, 2.0, 3.0, 4.0, or 5.0 Å (**Supplemental Table 2**). For $Mg^{2+}$ ions, the SimRNA-Sol model agreed better with our manual model (F-score 0.3 at 3.0 Å) than with the reference models (F-scores 0.18 and 0.21 at 3.0 Å). The manual model agreed better with the automatic reference 9CBX than with the consensus reference 9CBU, F-scores 0.20 and 0.14 at 3.0 Å for $Mg^{2+}$ ions and F-scores 0.18 and 0.11 at 3.0 Å for water molecules, respectively. The SimRNA-Sol model showed the opposite trend, agreeing better with the consensus reference. Notably, for water molecules, the SimRNA-Sol and manual models diverged greatly (F-score 0.02 at 3.0 Å), with the manual model being slightly closer to the references.

Overall, considering the large differences in absolute numbers of solvent moieties included in our two models, we can conclude that the SimRNA-Sol showed a reasonable conservative performance in terms of $Mg^{2+}$ ions, while the prediction of water molecules proved to be a harder task for SimRNA-Sol.

**Supplemental Text 18: Additional analysis by Cheatham-lab TS349, Cheatham-lab_villa T412 - Olivia Fisher and Thomas Cheatham**

Analysis of trajectories was done using CPPTRAJ and visualized in VMD and ChimeraX[96]. Hydrogen bond analysis of $Mg^{2+}$ interactions was done using a 4.0 Å distance cutoff, and RMSD was calculated in reference to the first frame.

*Results*

RMSD histograms of both divalent ion models include each individual replica, as shown in **Supplemental Figure 17**. Results confirm that the two models sample similar structural populations and indicate convergence. Longer simulation time would likely show further equilibration.

The representative structures for each model obtained from K-means clustering are shown in **Supplemental Figure 18**, overlayed with the R1260 EMD-42499 cryo-EM map to show the differences in conformation. Clear changes in structure are shown with overall surrounding ion placement.

The radial distribution function was used to calculate $Mg^{2+}$ ion density as a function of distance from RNA, as shown in **Supplemental Figure 19**. Compared to each other, the two ion models follow very similar distributions with only very slight differences. It is interesting to see overall distribution patterns, but when looking at the sum of fraction occupancies of $Mg^{2+}$ ions with RNA, as shown in **Supplemental Figure 20**, we see clear differences between the two ion models. Fraction occupancies were calculated for all four simulations in both ion models and separated into interactions of $Mg^{2+}$ with three groups: phosphate oxygen atoms, base oxygen and nitrogen atoms, and sugar oxygen atoms following protocols from Bergonzo et. al.[4] Within each interaction group, there are more higher occupancy locations in the Li Merz ion model than the Villa model. The Li Merz MG-phosphate group consistently had higher summed fraction occupancies dispersed across the RNA, which is also seen in the base and sugar groups although not as starkly.

*Discussion*

Our results using traditional MD simulation of the R1260 ribozyme show ion placement differences between two commonly used divalent ion models- Li Merz and Villa. CASP16 assessment shows slightly increased accuracy of the Villa model in cross correlation than the Li Merz model. This difference can be explained by the $Mg^{2+}$-RNA fractional occupancy bindings that show a consistent trend toward higher occupancy and more potential long-scale chelation events. Further studies may be useful to solidify these findings as well as improved on by enhanced sampling techniques.

**Supplemental Text 19: Additional analysis by bussilab_replex TS391, bussilab_plain_md TS485 - Elisa Posani, Olivier Languin Cattoen, and Giovanni Bussi**

The average RNA RMSD from native (7EZ0) was 0.05 nm in the conventional MD and 0.45 nm in the replica exchange simulation, consistent with the different stiffness of the applied restraints. **Supplemental Figure 21** shows that the equilibration of the bulk region was similar in the two simulations. However, the sampling of directly (inner shell) binding states was significantly faster in the replica exchange simulation. **Supplemental Table 3** reports the average number of ions in selected distance ranges from phosphate non-bridging oxygens, corresponding to inner-shell binding, outer-shell binding, and bulk regions, as well as the corresponding concentrations in the bulk. The conventional MD simulation displays a larger number of outer-shell bound ions compared to the replica exchange simulation. These ions were attracted by the RNA negative charge but could not undergo inner-shell binding in the simulated time.

**Supplemental Text 20: Additional analysis by Vfold TS481 - Yuanzhe Zhou, Sicheng Zhang, Jun Li, and Shi-Jie Chen**

Analysis of the simulation trajectories revealed that the MCTBI- or MgNet-predicted $Mg^{2+}$ ions show larger fluctuations than those in PDB 7EZ0[1], with ~2.43 Å for the 27 $Mg^{2+}$ ions in 7EZ0 and ~4.64 Å for the 90 $Mg^{2+}$ ions added according to MCTBI- or MgNet-predictions. This suggests that the $Mg^{2+}$ ions in the PDB structures, compared to the model-predicted ones, are more tightly coordinated and bound to the RNA.

17. Lin Z, Akin H, Rao R, et al. Evolutionary-scale prediction of atomic-level protein structure

with a language model. *Science*. 2023;379(6637):1123-1130.

33. Bonomi M, Branduardi D, Bussi G, et al. PLUMED: A portable plugin for free-energy

calculations with molecular dynamics. *Comput Phys Commun*. 2009;180(10):1961-1972.

49. Zhang X, Li S, Pintilie G, Palo MZ, Zhang K. Snapshots of the first-step self-splicing of

Tetrahymena ribozyme revealed by cryo-EM. *Nucleic Acids Res*. 2023;51(3):1317-1325.

65. Li P, Roberts BP, Chakravorty DK, Merz KM Jr. Rational design of Particle Mesh Ewald

compatible Lennard-Jones parameters for +2 metal cations in explicit solvent. *J Chem Theory Comput*. 2013;9(6):2733-2748.
